# A whole-animal phenotypic drug screen identifies suppressors of atherogenic lipoproteins

**DOI:** 10.1101/2024.11.14.623618

**Authors:** Daniel J. Kelpsch, Liyun Zhang, James H. Thierer, Adrian G. Rivera Cruz, Kobe Koren, Urmi Kumar, Yuki Lin, Monica R. Hensley, Mira Sohn, Jun O. Liu, Thomas Lectka, Jeff S. Mumm, Steven A. Farber

## Abstract

Lipoproteins are essential for lipid transport in all bilaterians. A single Apolipoprotein B (ApoB) molecule is the inseparable structural scaffold of each ApoB-containing lipoprotein (B-lps), which are responsible for transporting lipids to peripheral tissues. The cellular mechanisms that regulate ApoB and B-lp production, secretion, transport, and degradation remain to be fully defined. In humans, elevated levels of vascular B-lps play a causative role in cardiovascular disease. Previously, we have detailed that human B-lp biology is remarkably conserved in the zebrafish using an *in vivo* chemiluminescent reporter of ApoB (LipoGlo) that does not disrupt ApoB function. Thus, the LipoGlo model is an ideal system for identifying novel mechanisms of ApoB modulation and, due to the ability of zebrafish to generate many progeny, is particularly amenable to large-scale phenotypic drug screening. Here, we report a screen of roughly 3000 compounds that identified 49 unique ApoB-lowering hits. Nineteen hits passed orthogonal screening criteria and seven were subjected to extensive phenotyping. A licorice root component, enoxolone, significantly lowered B-lps only in animals that express a functional allele of the nuclear hormone receptor Hepatocyte Nuclear Factor 4⍺ (HNF4⍺). Consistent with this result, inhibitors of HNF4⍺ also reduce B-lp levels. These data demonstrate that mechanism(s) of action can be rapidly determined from a whole animal zebrafish phenotypic screen. Given the well documented role of HNF4⍺ in human B-lp biology, these data validate the LipoGlo screening platform for identifying small molecule modulators of B-lps that play a critical role in a leading cause of worldwide mortality.

## Introduction

Lipoproteins are critical particles that transport lipids in the aqueous plasma of animals and profoundly impact human physiology and disease. One subset of lipoproteins, the (ApoB)-containing lipoproteins (B-lps), transport lipids from sites of absorption and synthesis (e.g., intestine and liver) through the bloodstream to peripheral tissues. Excess serum B-lps are a well-established causative factor in the initiation and progression of cardiovascular disease (CVD) [1,2] and are associated with elevated risk for insulin resistance and hepatic steatosis [3–5]. CVD encompasses heart failure, stroke, and heart-related abnormalities and accounts for ∼1 million lost lives in the US every year [6]. In cells that synthesize B-lps, each particle is formed by the transfer of lipids to a single ApoB molecule via the activity of microsomal triglyceride transfer protein (MTP) at the endoplasmic reticulum [7–9]. Since each B-lp contains only one ApoB molecule, the total ApoB level is directly proportional to the number of B-lps. Despite a rich literature on the factors that regulate MTP, ApoB, and B-lp levels, many mechanisms that modulate B-lp production, secretion, transport, uptake, and turnover remain unknown [10,11].

Clinically, several effective therapies are currently used to reduce circulating B-lps in individuals with hyperlipidemia or CVD. Statins directly inhibit cholesterol biosynthesis, ultimately promoting hepatic uptake of cholesterol rich circulating B-lps reducing the atherogenic potential of excess B-lps (LDL) in circulation [12,13]. However, statins can also cause side effects, with the most severe being autoimmune myopathy and rhabdomyolysis and, more rarely, stroke, liver disease, and diabetes [14–19]. Statin therapy is often used in conjunction with the intestinal cholesterol absorption inhibitor ezetimibe [20–22]. However, ezetimibe is rarely used as a monotherapy due to its relatively modest effect on serum B-lps. Moreover, some individuals do not respond to statin-based therapies [16,23]. One recently approved compound, Bempedoic acid, inhibits cholesterol biosynthesis similar to statins and demonstrates a B-lp lowering effect among statin-intolerant individuals [24]. However, long-term studies of bempedoic acid are ongoing. These compounds effectively reduce the risk of cardiovascular events, but often, patients on these compounds still die of CVD [12,24].

One strategy to reduce B-lp levels is through a small molecule inhibitor of MTP activity (e.g., lomitapide) [25]. While lomitapide effectively reduces B-lp levels, its use is associated with adverse side effects, including hepatic steatosis and endoplasmic reticulum stress, eventually causing hepatic fibrosis, and increased intestinal triglyceride accumulation, leading to elevated fecal lipids and gastrointestinal symptoms. Due to these severe side effects, lomitapide is prescribed mainly to individuals with familial hypercholesterolemia, which is characterized by elevated plasma B-lps and cholesterol from birth. Mipomersen, an antisense oligonucleotide targeting ApoB, reduces B-lps synthesis but is associated with severe hepatotoxicity [26]. A newer therapeutic strategy to reduce circulating B-lps is to block the activity of PCSK9 (Proprotein Convertase Subtilisin/Kexin type 9). PCSK9 promotes low-density lipoprotein (LDL)-receptor (LDLR) degradation by blocking the recycling of this receptor back to the plasma membrane [27,28]. Thus, PCSK9 inhibitors promote liver LDLR recycling and increase B-lp uptake. Current PCSK9 therapies are monoclonal antibody-based and must be injected every several weeks, feature some adverse effects in patients, and are costly. Small molecule inhibitors and other approaches that target PCSK9 are currently in development [29–31].

Despite an array of treatment options for hyperlipidemia, CVD is still the leading cause of death worldwide [6]. Moreover, the strategies mentioned above are rarely effective in reducing another ApoB-containing lipoprotein, Lipoprotein(a), affecting approximately ∼20% of individuals in the US [32,33]. Patients with genetic mutations in the Lipoprotein(a) encoding gene have a 2 to 4-fold increased risk of sudden heart attack or stroke. Taken together, there remains a pressing need for new strategies that reduce circulating B-lps.

One factor that makes identifying B-lp-reducing compounds difficult is the challenge of replicating the multiple cell types and multiorgan physiology that are known to impact plasma levels of B-lps in a simple, easy-to-screen cell culture system. This issue is circumvented by using an intact whole vertebrate, though this is difficult with mammalian models due to cost and scale. We previously established the larval zebrafish (*Danio rerio*) as an excellent model for ApoB biology in that most of the genes known to regulate human B-lp synthesis and metabolism are present [34,35]. This includes the gene that encodes cholesteryl ester transfer protein (CETP), a central regulator of human lipoprotein homeostasis that is deficient in mouse and rat models [36]. Thus, the zebrafish plasma lipid profile is more similar to humans than many rodent models in that the bulk of plasma cholesterol is carried in B-lps (as opposed to ApoA1-containing lipoproteins like HDL) [37,38]. To study ApoB biology in fish, we generated a reporter of B-lps called LipoGlo, where NanoLuciferase (NL) was fused to the gene that encodes nearly all ApoB protein in the zebrafish, *apoBb.1*, at its endogenous locus [35]. This fusion does not disrupt normal B-lp biology. Since each B-lp is associated with a single ApoB molecule, the luminescence generated by LipoGlo-expressing animals is directly proportional to the total number of B-lps.

Another major advantage of the larval zebrafish model system is that during early development, zebrafish do not ingest food, and B-lps transport lipids from the nutrient-dense maternally deposited yolk to peripheral tissues [39]. Thus, any variability in B-lp levels due to the amount and type of food consumed is eliminated in early zebrafish development. Moreover, the zebrafish has developed all major digestive organs by five days post fertilization (dpf), a period where growth is not dependent on exogenous feeding. The larval zebrafish is also quite small (∼5 mm in length), making it amenable to large-scale robotic screening, and several drugs from zebrafish screens are currently in clinical development [40–42].

Using these unique attributes of zebrafish larvae, we performed a whole-animal phenotypic drug screen to identify chemical modulators of B-lps. We screened a library of ∼3000 compounds and identified 49 unique compounds that lowered B-lp in larval zebrafish and validated 19 as lead compounds that are the basis of ongoing studies. One lead compound, enoxolone, has many reported potential activities that includes inhibition of Hepatocyte Nuclear Factor 4⍺ (HNF4⍺), an established transcriptional regulator of lipoprotein metabolism [43–45]. We report that known HNF4⍺ inhibitors reduce B-lp levels to levels equivalent to genetic depletion of HNF4⍺ in larval zebrafish. Further, HNF4⍺ expression is required for enoxolone to exert its B-lp lowering effect in the fish. The finding that enoxolone suppresses B-lps in the zebrafish through modulation of HNF4⍺ demonstrates the remarkable conservation of B-lp metabolism in the zebrafish system and supports our phenotypic drug screening paradigm.

## Results

### A whole-animal drug screen to identify B-lp-reducing compounds

Using the previously characterized B-lp zebrafish reporter line, LipoGlo [35], we undertook a small-molecule screen to identify B-lp modulating compounds *in vivo*. The screen design was guided by the normal developmental profile of B-lp metabolism in the zebrafish (Figure 1A) [35]. The embryonic zebrafish begins to mobilize lipids stored within the yolk and starts to secrete B-lps from 1 to 3 dpf. After 4 dpf, as yolk lipids are depleted, B-lp numbers concomitantly decrease (Figure 1A). During embryonic and larval stages that are yolk-dependent (lecithotrophic period), zebrafish do not require feeding, reducing the variability introduced by the amount and type of food consumed when experiments are performed at stages after the yolk is depleted. Thus, we developed an automated screening platform where animals are treated at 3 dpf for 48 h with test compounds, and then at 5 dpf they are fixed and assayed for whole-animal luminescence as a readout for total B-lp level. Animals treated on 3 dpf with lomitapide, an MTP inhibitor, for 48 hours typically exhibited a >50% reduction in B-lp levels making it a strong positive control for a drug screen (Figure 1B) [35].

**Figure 1.**
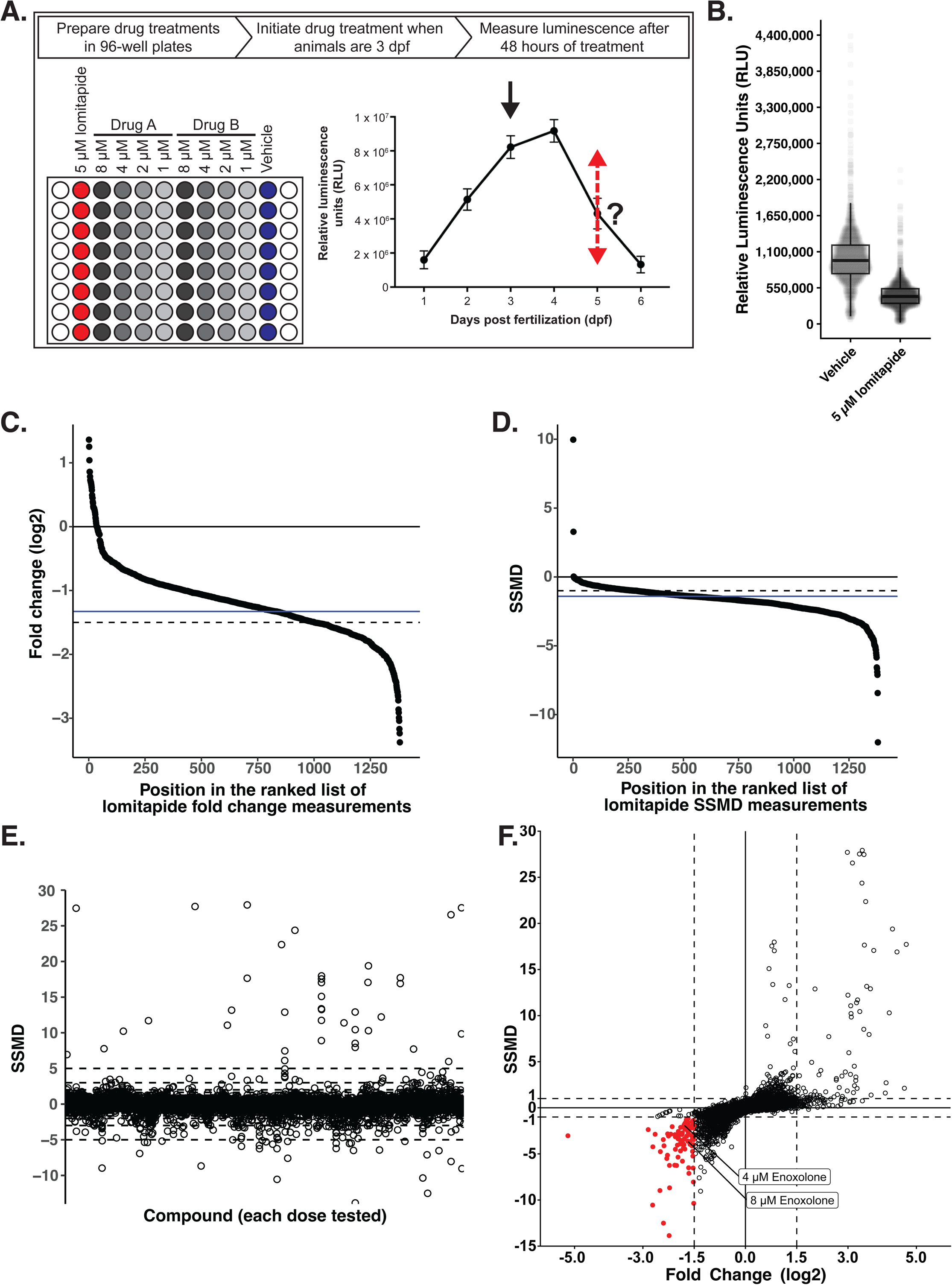
A whole-animal drug screen identifies LipoGlo-reducing compounds. **(A)** Schematic summarizing drug screening paradigm. Drug treatments were prepared in 96-well plates, each plate with a negative control (vehicle), positive control (5 µM lomitapide), and serial dilutions (four-fold dilution; 8 µM, 4 µM, 2 µM, and 1 µM) of two different drugs of interest. Each treatment was prepared with 8 replicates. When animals were 3 dpf, when B-lp levels are relatively high (black arrow), they were dispensed into their drug treatment for a 48-hour incubation when luminescence was measured (red dashed arrow). **(B)** Boxplot of average Relative Luminescence Units (RLU) measured from fixed 5 dpf *Fus(ApoBb.1-NanoLuciferase); Tg(ubi:mcherry-2A-FireflyLuciferase)* animals treated for 48 hours with either negative (vehicle) or positive (5 µM lomitapide) control. Each data point represents the average of 8 independent samples measured from a single 96-well plate from 1381 independent experiments across the entire screen. We measured a 55.6% reduction in RLU in 5 µM treated animals. **(C)** An ordered plot of each average fold change of luminescence (log2 scale) measured from 5 µM lomitapide treated animals from each 96-well plate (n = 1381) relative to respective vehicle treatment. The solid black line at y=0 represents the divide in increased and decreased luminescence levels, the solid blue line at y = -1.33 represents the curve’s inflection point, and the dashed black line at y = -1.5 represents the fold change cutoff used to define a hit. **(D)** An ordered plot of each SSMD score measured from positive control (5 µM lomitapide) treated animals from each 96-well plate (n = 1381) relative to respective vehicle treatment. The solid black line at y = 0 represents the divide in increased and decreased SSMD score, the solid blue line at y = -1.41 represents the curve’s inflection point, and the dashed black line at y = -1 represents the SSMD (open circles) cutoff used to define a hit. **(E)** A plot of SSMD scores measured from each drug at each dose tested, each open circle represents the SSMD score of an individual drug at an individual dose. A total of 2762 drugs were tested, each at 4 different doses (8, 4, 2, and 1 µM; n = 11048). Dashed lines at y = ±1, ±1.25, ±1.645, ±2, ±3, ±5 represent common defining cutoffs of SSMD scores. **(F)** A dual flashlight plot of each dose of each drug (open circles) SSMD score (y-axis) against fold change (log2-scale, x-axis). Dashed lines at y = -1, y = 1, x = -1.5, and x = 1.5 represent the cut-off to define hits that significantly affect luminescence levels; all significant luminescence-reducing compounds are highlighted in red (n = 50).

One established drug screening approach is a repositioning strategy, which identifies whether any approved or investigational drugs have new or unintended advantageous biological effects [46]. This approach has identified new indications for numerous drugs [46–48]. Drug repositioning screens are advantageous over novel screening as they feature a lower risk of failure, reduced time frame to development, and reduced cost (for review, see [46]).

To accomplish such a large-scale effort, we utilized the ARQiv-HTS platform (Automated Reporter Quantification *in vivo* coupled to High-Throughput Screening robotics) [49] to array and dilute drugs from the Johns Hopkins Discovery Library (JHDL) in a 4-point serial dilution (8 µM, 4 µM, 2 µM, and 1 µM), in addition to positive (5 µM lomitapide) and negative (vehicle) controls (Figure 1A). The JHDL comprises roughly 3000 compounds – many of which have known mechanisms of action and are approved for human use [50]. Each treatment condition included 8 independent replicates, a number selected based on the typical variation observed in the B-lp assay observed in prior experiments and power calculations. LipoGlo-expressing larvae (3 dpf) were dispensed into their respective drug treatment and reared for 48 hours at 27°C on a 14:10 h light:dark cycle. Previously, we have characterized whole-animal B-lp levels from whole-animal homogenates [35]. However, in order to simplify the screening process, we turned to fixing whole animals in place before measuring total luminescence. At 5 dpf, animals were briefly fixed with paraformaldehyde, incubated in NanoLuciferase substrate, and B-lp levels were measured with a plate reader (Figure 1A).

### Identification of B-lp-reducing compounds

We measured the effect of drug treatments across 1381 96-well plates, each plate containing negative controls, positive controls, and a serial dilution of two independent drugs (2762 total compounds) from the JHDL (Figure 1A, Supplemental Table 1). To provide insight into our B-lp screen sensitivity in relation to effect size we quantified the effect of the positive control (5 µM lomitapide) treatment across all 1381 plates. On average, lomitapide produced 55.6% ± 23.2% reduction in B-lp levels (Figure 1B), consistent with our prior studies [35]. We then rank-ordered the fold change of B-lp levels for each 96-well plate following lomitapide treatment (Figure 1C). The rank-ordered list of fold change further confirms that lomitapide reduces B-lps across many samples (Figure 1C, values < 0 and below solid black line), but there is a modest level of plate-to-plate variation. Thus, we defined compounds that reduce B-lps as those compounds with a fold change of < -1.5 (log_2_ scale; Figure 1C, dashed line). Interestingly, we find that the inflection of rank-ordered fold change is -1.33 (Figure 1C, solid blue line) and likely represents a more specific hit-defining cutoff for this dataset.

We then set out to deploy a statistical measure to help us define a hit. In numerous prior high-throughput screens with many replicates per group, the Strictly Standardized Mean Difference (SSMD) was an ideal statistical measure [51,52]. SSMD estimates the fold change penalized by the variability of that fold change and is robust to variation and outliers in screens. We calculated the SSMD for lomitapide treatment from each plate. We confirmed a reduction in B-lp levels (Figure 1D, values < 0 and below solid black line) with an average SSMD of -1.76 (n = 1381), which is considered “fairly strong” by the developers of this metric [51,52]. Prior research uses an SSMD of 1.0 as a cutoff to define the significance threshold of a hit. Thus, we defined compounds that reduce B-lps as those compounds with an SSMD of < -1.0 (Figure 1D, dashed line). Interestingly, we determined that the inflection of rank-ordered SSMD scores for lomitapide treatments is -1.41 (Figure 1D, solid blue line), which likely represents a more specific hit-defining cutoff for this dataset. Ultimately, the robust reduction of B-lps following 5 µM lomitapide treatment on a large scale demonstrates proof of concept for a large scale drug screen to identify pharmacological agents that modulate ApoB levels in a whole animal.

We then quantified SSMD of each dose of each drug treatment to determine the overall effect size of that drug treatment (Figure 1E) and identified 487 unique drugs that lower B-lp levels with an SSMD cutoff of < -1.0 (Figure 1E, Supplemental Table 2). While the SSMD score is typically very robust, it is sensitive to weak effects with low variation, so biological effect size was also considered when calling screen hits [51,52]. We assessed the SSMD score and fold change of each drug dose (Figure 1F). Taken together, we classified hits as those drugs with an SSMD < -1 and fold change of < -1.5 (log_2_ scale), in line with cutoffs previously defined [51,52]. Using these cutoffs, we identified 50 (49 unique as fendiline was tested twice) B-lp-reducing compounds (Figure 1F, Figure 2, Supplemental Table 3, Supplemental Figure 1). However, as noted above, we identified the inflection of rank-ordered fold change values and SSMD scores across 1381 positive control experiments as -1.33 and -1.41, respectively (Figures 1C and 1D). Using the new hit cutoffs, there are minor changes in the profile of hit compounds; calcipotriene and diphenylboric acid are no longer hits, and twelve compounds (2-phenpropylamino-5-nitrobenzoic acid, cetalkonium chloride, clioquinol, clomipramine, dioctylnitrosamine, docusate sodium, gliclazide, guaiacwood, quipazine maleate, rose bengal, sodium anthraquinone, and vitamin K1) are classified as hits.

**Figure 2.**
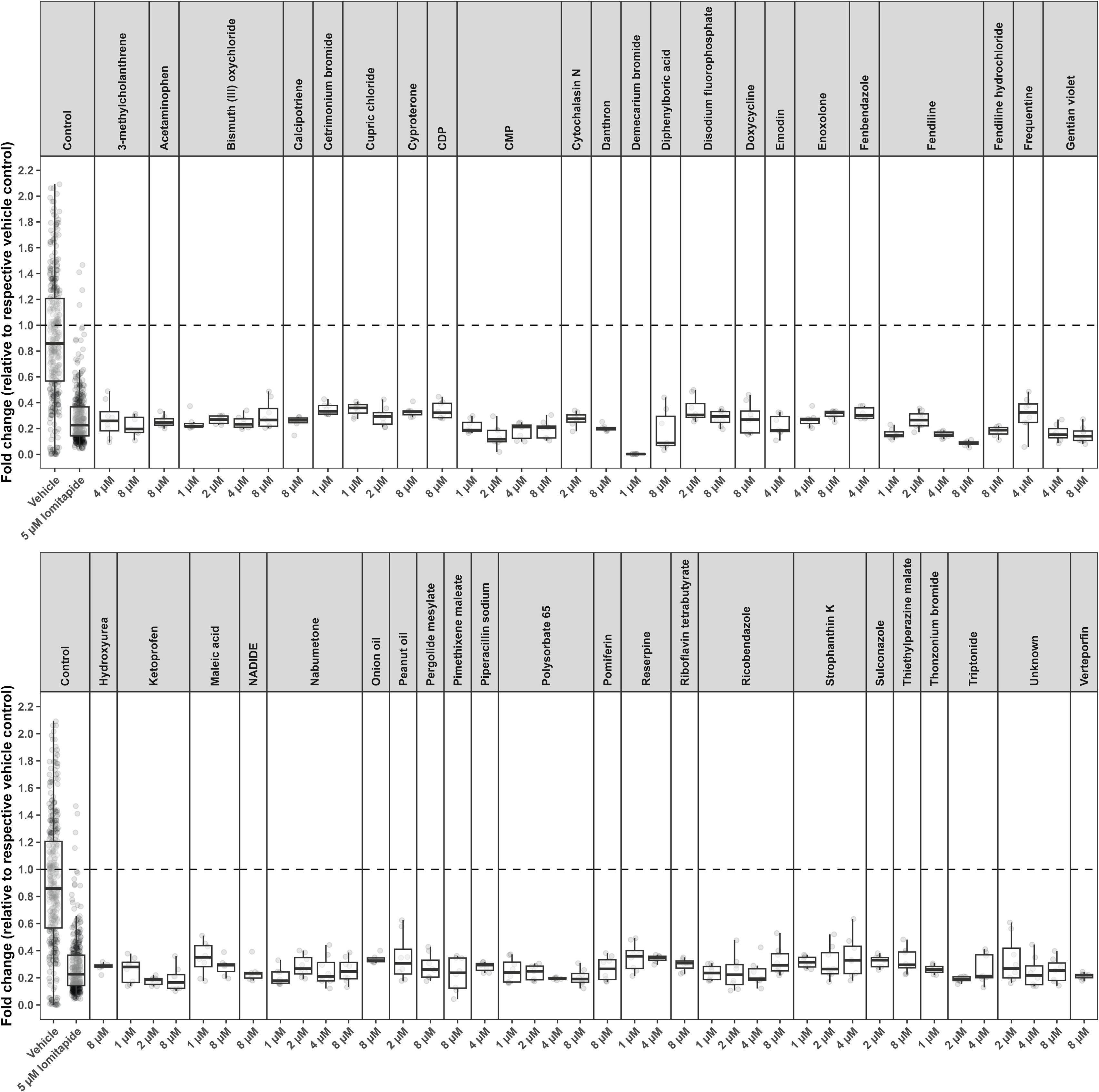
Fifty total compounds significantly reduce LipoGlo levels. Boxplot of the fold change for each of the 50 hit compounds from the Johns Hopkins Drug Library (JHDL) at the respective dose they met hit criteria (fold change (log2) ≤ -1.5 and SSMD ≤ -1). The luminescence of each drug at each dose was measured (n = 8), and fold change was calculated against the mean of vehicle treatment.

### Validating the B-lp lowering effects of 49 unique hits

We identified 49 unique compounds with SSMD < -1 and fold change of < -1.5 (log_2_ scale) and aimed to validate the B-lp lowering effects of each of these hits. Drug screening libraries are typically prepared in batches and stored for long periods of time. Thus, validating the effect of a fresh batch of each hit identified from a drug screen is essential. Of our 49 unique hits, one compound could not be identified (Supplemental Table 3), and 11 have yet to be obtained and/or validated (Supplemental Table 3). The remaining 37 hit compounds were purchased from commercial suppliers (Supplemental Table 3) and LipoGlo-expressing animals were treated with each compound using an 8-fold serial dilution with between 1 and 3 biological replicates, each with 8 technical replicates (Supplemental Table 3, Supplemental Figure 2).

Gentian violet caused lethality at all doses tested (Supplemental Table 3 and Supplemental Figure 2). Thirteen more compounds, acetaminophen, cytidine 5’-monophosphate, fenbendazole, ferron, hydroxyurea, ketoprofen, medroxyprogesterone acetate, maleic acid, NADIDE, piperacillin sodium, prochlorperazine dimaleate, ricobendazole, and strophanthin K failed to show any effect on B-lp levels (Supplemental Table 3 and Supplemental Figure 2). Surprisingly, two compounds, sulconazole (Supplemental Figure 2AE) and thonzonium bromide (Supplemental Figure 2AH), caused an increase in B-lps levels (Supplemental Table 3). Notably, 21 compounds, 3-methylcholanthrene, cetrimonium bromide, cyproterone, calcipotriene, danazol, danthron, demecarium bromide, doxycycline, emodin, enoxolone, fendiline, nabumetone, pergolide mesylate, pimethixene maleate, pomiferin, reserpine, riboflavin tetrabutyrate, triptonide, thiethylperazine malate, and verteporfin each significantly reduced B-lps at ≥1 dose examined, thus validating the primary screen effect (Supplemental Table 3 and Supplemental Figure 2).

While these hits reduce B-lp levels as measured by luminescence generated by the LipoGlo reporter, it is unclear whether this is due to a reduction of B-lps or if a hit interferes with the enzymatic activity of NanoLuciferase. To determine if a hit was a luciferase inhibitor, we generated a tissue homogenate from untreated 3 dpf LipoGlo expressing animals and briefly incubated aliquots of homogenate with vehicle or a serial dilution of each hit. Only verteporfin showed a dose-dependent decrease in luminescence activity, indicating it is a NanoLuciferase inhibitor and likely does not affect B-lp levels (Supplemental Table 3). Ultimately, our screening efforts identified 19 compounds that reduce B-lp levels through unknown mechanisms (Supplemental Table 3).

We classified these remaining hits according to their Medical Subject Heading (MeSH) terms [53]. We found amines (cetrimonium bromide, demecarium bromide, and fendiline), biological pigments (riboflavin tetrabutyrate), heterocyclic compounds (pergolide mesylate, pomiferin, and reserpine), hormones, hormone substitutes, and hormone antagonists (cyproterone and danazol), hydrocarbons (3-methylcholanthrene, doxycycline, enoxolone, and triptonide), ketones (nabumetone), micronutrients (calcipotriene), quinones (danthron and emodin), and sulfur compounds (pimethixene maleate and thiethylperazine malate).

### Secondary characterization of selected validated B-lp lowering hits

To further evaluate the biological relevance and potential mechanisms of selected validated hits, we performed secondary analyses assessing total B-lp levels, larval morphology, and lipoprotein size distribution. Our lab previously defined the method to measure total B-lp levels in a whole animal by using the homogenate of a single zebrafish larva [35]. We confirmed that treatment with 4 µM pomiferin significantly reduced total B-lp levels measured from homogenates collected after treatment (p = 8.6×10^−4^; Figure 3A). However, pomiferin treatment (4 µM) produced animals with reduced body length and lethality at higher doses, suggesting developmental toxicity may confound interpretation of its B-lp lowering effect.

**Figure 3.**
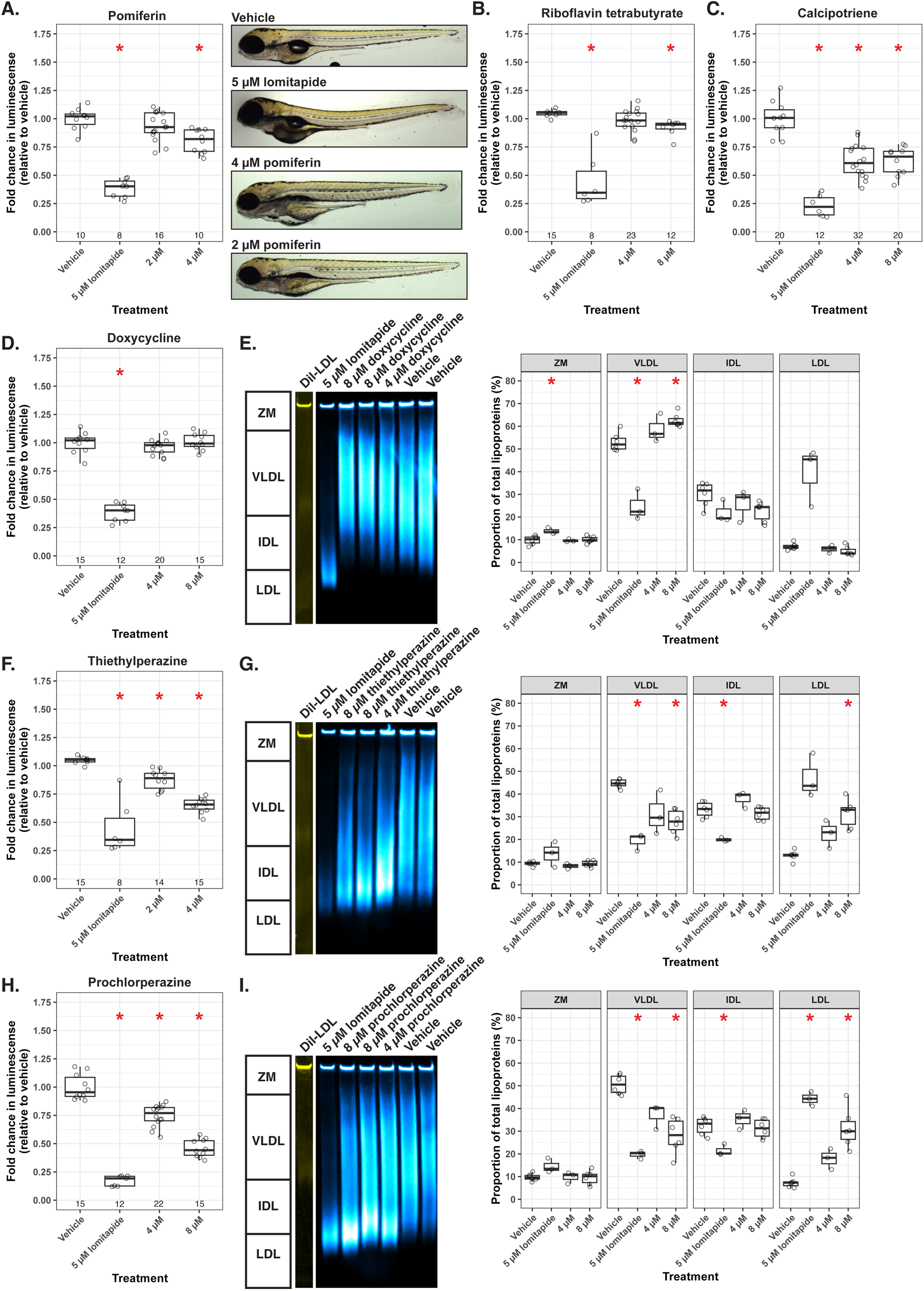
Validation of hits reveals different phenotypes for different drugs. **(A-D, F, H)** Average fold change of Relative Luminescence Units (RLU) measured from homogenates of 5 dpf Fus(ApoBb.1-NanoLuciferase); Tg(ubi:mCherry-2A-FireflyLuciferase) animals treated for 48 hours with either negative (vehicle), positive (5 µM lomitapide) control, or a serial dilution of the indicated compound. Each data point represents a measurement from an independent animal collected from two **(A)** or three (B, D, F, H) or four **(C)** independent experiments and normalized to the average vehicle RLU from each individual experiment. Sample size (n) is listed below each treatment. Statistical significance was determined by one-way ANOVA followed by Dunnett’s test against vehicle, with Bonferroni correction. Significant treatments (red asterisks) and their adjusted p-values are reported below. **(A)** Pomiferin treatment significantly reduced RLU levels (one-way ANOVA, F(3,40) = 61.85, p = 4.4×10^−15^). Significant Dunnett’s comparisons: 5 µM lomitapide (p = 7.6×10^−11^) and 4 µM pomiferin (p = 8.6×10^−4^). Representative whole-mount images of vehicle, 5 µM lomitapide, 2 µM pomiferin, and 4 µM pomiferin-treated animals are shown. **(B)** Riboflavin tetrabutyrate: one-way ANOVA F(3,54) = 68.85, p < 2×10^−16^; significant comparisons: 5 µM lomitapide (p = 5.0×10^−7^) and 8 µM riboflavin tetrabutyrate (p = 5.2×10^−3^). **(C)** Calcipotriene: one-way ANOVA F(3,80) = 61.47, p < 2×10^−16^; significant comparisons: 5 µM lomitapide (p = 8.0×10^−17^), 4 µM calcipotriene (p = 2.9×10^−8^), and 8 µM calcipotriene (p = 1.6×10^−6^). **(D)** Doxycycline: one-way ANOVA F(3,58) = 126.5, p < 2×10^−16^; significant comparison: 5 µM lomitapide (p = 1.7×10^−14^). (F) Thiethylperazine: one-way ANOVA F(3,48) = 82.76, p < 2×10^−16^; significant comparisons: 5 µM lomitapide (p = 5.0×10^−7^), 4 µM thiethylperazine (p = 1.4×10^−10^), and 2 µM thiethylperazine (p = 4.3×10^−3^). **(H)** Prochlorperazine: one-way ANOVA F(3,60) = 66.42, p < 2×10^−16^; significant comparisons: 5 µM lomitapide (p = 1.6×10^−11^), 8 µM prochlorperazine (p = 1.2×10^−6^), and 4 µM prochlorperazine (p = 7.0×10^−7^). **(E, G, I)** Representative native-PAGE images of luminescent B-lps from homogenates of 5 dpf animals treated with vehicle, 5 µM lomitapide, or the indicated compound for 48 hours. Each image is a composite of chemiluminescence (B-lps, cyan hot) and fluorescence (DiI-LDL, yellow). B-lps were binned into one of four classes: zero mobility (ZM), very-low-density lipoproteins (VLDL), intermediate-density lipoproteins (IDL), or LDL, and visualized via boxplot. Each gel image is representative of one of three independent experiments. Statistical significance was determined by one-way ANOVA and post-hoc Dunnett’s test within each lipoprotein class. Significant comparisons (red asterisks) for each panel: **(E)** doxycycline ZM class: one-way ANOVA F(3,14) = 5.38, p = 1.1×10^−2^; significant comparison: 5 µM lomitapide (p = 1.3×10^−2^). Doxycycline VLDL class: one-way ANOVA F(3,14) = 47.69, p = 1.3×10^−7^; significant comparisons: 5 µM lomitapide (p = 3.0×10^−2^), 8 µM doxycycline (p = 3.5×10^−3^). **(G)** Thiethylperazine VLDL class: one-way ANOVA F(3,14) = 19.0, p = 3.3×10^−5^; significant comparisons: 5 µM lomitapide (p = 1.0×10^−2^), 8 µM thiethylperazine (p = 8.4×10^−4^). Thiethylperazine IDL class: one-way ANOVA F(3,14) = 20.4, p = 2.2×10^−5^; significant comparison: 5 µM lomitapide (p = 2.6×10^−4^). Thiethylperazine LDL class: one-way ANOVA F(3,14) = 24.8, p = 7.2×10^−6^; significant comparison: 8 µM thiethylperazine (p = 1.2×10^−3^). **(I)** Prochlorperazine VLDL class: one-way ANOVA F(3,14) = 24.8, p = 7.4×10^−6^; significant comparisons: 5 µM lomitapide (p = 3.8×10^−6^), 8 µM prochlorperazine (p = 9.1×10^−4^). Prochlorperazine IDL class: one-way ANOVA F(3,14) = 7.5, p = 3.0×10^−3^; significant comparison: 5 µM lomitapide (p = 2.4×10^−3^). Prochlorperazine LDL class: one-way ANOVA F(3,14) = 33.8, p = 1.1×10^−6^; significant comparisons: 5 µM lomitapide (p = 1.2×10^−3^), 8 µM prochlorperazine (p = 2.3×10^−3^).

Treatment of animals with riboflavin tetrabutyrate (Figure 3B) and calcipotriene (Figure 3C) reduced total B-lp levels (p < 2×10^−16^) but did not affect larval morphology. A key feature of B-lps is their size, often a proxy for the total amount of lipid in the particle [35]. Particle size can impact the particle’s lifetime (e.g. in metabolically healthy humans, small particles are cleared rapidly by the liver) [54–56]. Thus, we also assessed whether these compounds alter B-lp size distribution. Animals were treated for 48 h with vehicle, 5 µM lomitapide, or a drug of interest, and whole-animal homogenates were prepared and subjected to native polyacrylamide gel electrophoresis followed by chemiluminescent imaging. B-lps were classified into four classes based on gel migration: zero mobility (ZM), very low-density lipoproteins (VLDL), intermediate-density lipoproteins (IDL), and low-density lipoproteins (LDL) as previously described [35]. Lomitapide treatment effectively reduces VLDL particles and increases LDL particles [35] (Figure 3E), whereas riboflavin and calcipotriene did not affect lipoprotein classes. Thus, riboflavin tetrabutyrate and calcipotriene reduce total B-lp levels without overt developmental toxicity or changes in lipoprotein subclass distribution, suggesting they may act through mechanisms that decrease overall particle abundance rather than altering lipoprotein turnover or catabolism.

Although doxycycline treatment lowered B-lp levels in whole animals in the primary screen and validation studies, we did not observe a reduction in total B-lps in whole-animal homogenates (Figure 3D). However, we detected a slight increase in VLDL levels (p < 2×10^−16^; Figure 3E), suggesting that doxycycline may alter lipoprotein composition or distribution rather than total particle abundance.

Alternatively, two structurally related compounds, thiethylperazine and prochlorperazine, at 4 µM significantly reduced (p < 1.4×10^−10^ and p < 1.2×10^−6^ respectively), B-lp levels measured from whole-animal homogenates (Figure 3F and 3H). Furthermore, both 8 µM thiethylperazine and 8 µM prochlorperazine increased relative LDL (p = 1.2×10^−3^ and p = 2.3×10^−3^, respectively) and decreased relative VLDL levels (p = 8.4×10^−4^ and p = 9.1×10^−4^, respectively; Figure 3G and 3I) suggesting a shift toward smaller lipoprotein particles and a potential alteration in lipid processing or clearance pathways. Together, these results highlight a diversity of mechanisms among validated hits, ranging from compounds that reduce total B-lp abundance without affecting B-lp class composition to those that shift B-lp class distribution, while also underscoring the importance of secondary assays to distinguish true B-lp modulators from those that likely produce a B-lp effect through generalized toxicity.

### Enoxolone significantly reduces B-lps in the larval zebrafish

Hit compounds were prioritized for follow-up studies based on reproducible dose-dependent responses, minimal toxicity as indicated by normal morphology over development, lack of direct NanoLuciferase inhibition, and prior reports suggesting potential links to lipid metabolism. One compound meeting these criteria was enoxolone, also known as 18β-Glycyrrhetinic acid, (Figure 2, Supplemental Table 1, Supplemental Figure 1T, Supplemental Figure 2L). Enoxolone is a triterpenoid derived from licorice root (*Glycyrrhiza glabra*) [57]. Evidence suggests that enoxolone perturbs several biological pathways and has reported anti-inflammatory, anti-oxidant, anti-viral, and antimicrobial effects [44,58–61]. Further, several studies suggest enoxolone can modulate lipid metabolism, although the mechanisms mediating these effects are poorly described or remain unclear [44,45,62]. For example, recent studies demonstrate that enoxolone treatment can ameliorate diet-induced hyperlipidemia, rescuing the effects of a high-fat diet on plasma lipoproteins and lipids [44]. Consistent with this finding, we observed a significant reduction in B-lp levels following enoxolone treatment with two different experimental endpoints. First, we validated our primary screen assay and found a significant reduction in B-lp levels following 8 µM, 4 µM, 2 µM, 1 µM, 0.5 µM, and 0.0625 µM enoxolone treatments from whole fixed animals compared to vehicle treatment (Figure 4A). We confirmed that 8 µM enoxolone treatment significantly reduced total B-lp levels measured from homogenates collected from whole animals after treatment (Figure 4B). Given that we assay B-lp levels by measuring NanoLuciferase activity, it was necessary to confirm that enoxolone specifically reduces B-lp levels and does not directly inhibit NanoLuciferase (Figure 4C).

**Figure 4.**
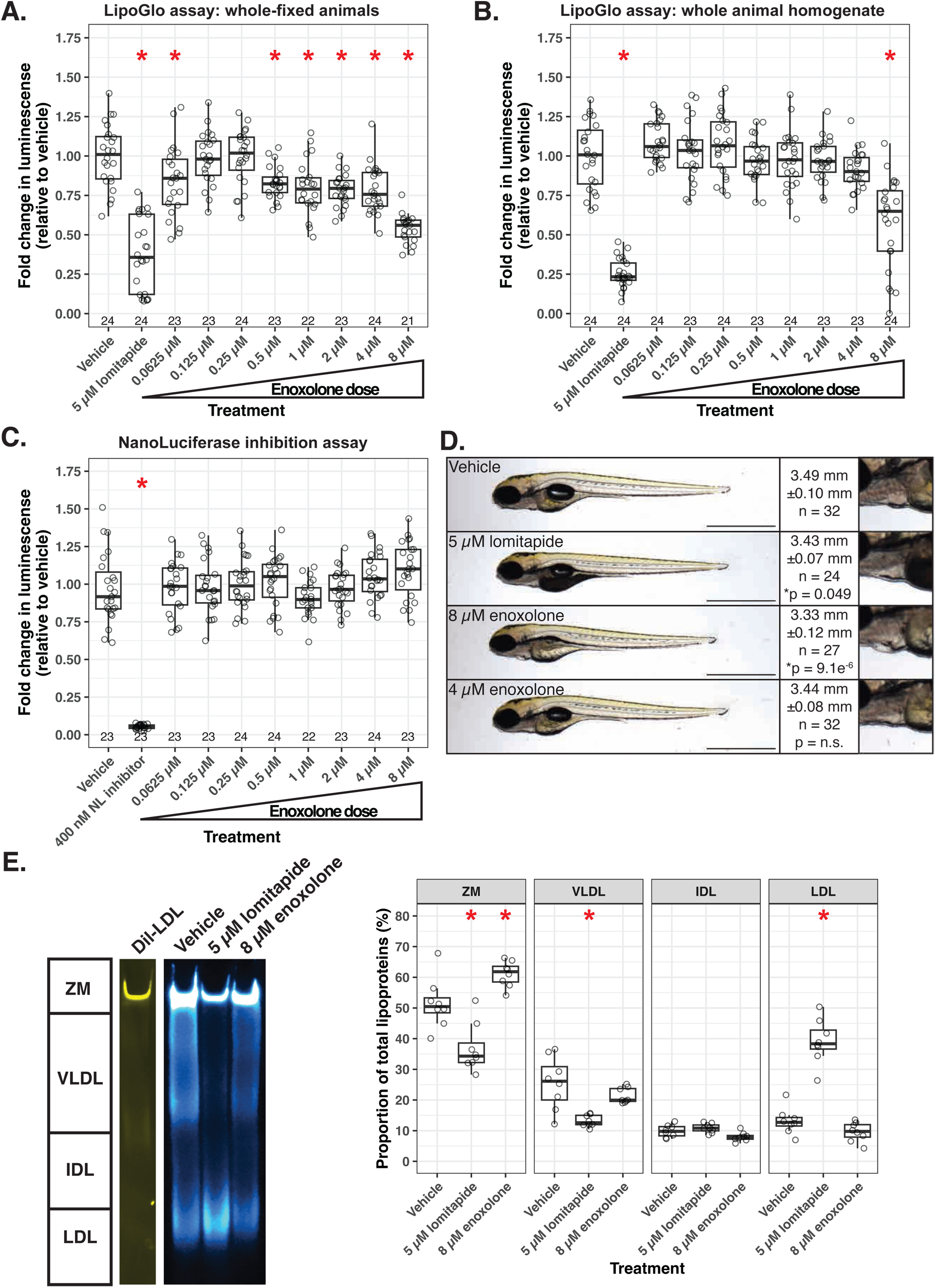
Enoxolone reduces B-lps in larval zebrafish. **(A)** Boxplot of the average fold change of Relative Luminescence Units (RLU) measured from fixed 5 dpf *Fus(ApoBb.1-NanoLuciferase); Tg(ubi:mcherry-2A-FireflyLuciferase)* animals treated for 48 hours with either negative (vehicle), positive (5 µM lomitapide) control, or an 8-fold serial dilution of enoxolone. Each data point represents a measurement from an independent animal collected from three independent experiments and normalized to the average vehicle RLU from each individual experiment. Several treatments significantly altered RLU levels (one-way ANOVA, *F*(9,221) = 32.79, *p* < 2×10^−16^). Lomitapide treatment (n = 24) significantly reduced RLU levels compared to vehicle treatment (n = 24, Dunnett’s test *p* = 5.6×10^−10^). Treatment with 8 µM enoxolone (n = 21, Dunnett’s test *p* = 7.8×10^−11^), 4 µM enoxolone (n = 24, Dunnett’s test *p* = 1.2×10^−3^), 2 µM enoxolone (n = 23, Dunnett’s test *p* = 3.2×10^−4^), 1 µM enoxolone (n = 22, Dunnett’s test *p* = 5.2×10^−3^), and 0.5 µM enoxolone (n = 23, Dunnett’s test *p* = 7.2×10^−3^) also reduced total RLUs. **(B)** Boxplot of the average fold change of RLUs measured from homogenized 5 dpf *Fus(ApoBb.1-NanoLuciferase); Tg(ubi:mcherry-2A-FireflyLuciferase)* animals that were treated for 48 hours with either vehicle, 5 µM lomitapide, or an 8-fold serial dilution of enoxolone. Several treatments significantly altered RLU levels (one-way ANOVA, *F*(9,226) = 54.2, *p* < 2×10^−16^). Lomitapide treatment (n = 24) significantly reduced RLU levels compared to vehicle treatment (n = 24, Dunnett’s test *p* = 1.7×10^−15^), as did 8 µM enoxolone treatment (n = 24, Dunnett’s test *p* = 6.1×10^−6^). **(C)** Boxplot of the average fold change of RLUs measured from untreated homogenates of 5 dpf *Fus(ApoBb.1-NanoLuciferase); Tg(ubi:mcherry-2A-FireflyLuciferase)* animals that were briefly treated with either vehicle, 400 nM NanoLuciferase inhibitor, or an 8-fold serial dilution of enoxolone to determine if enoxolone is an inhibitor of NanoLuciferase enzymatic activity. Only one treatment altered RLU levels (one-way ANOVA, *F*(9,221) = 78.31, *p* < 2×10^−16^), which was the positive control of 400 nM NanoLuciferase inhibitor (n = 22) when compared to vehicle treatment (n = 23, Dunnett’s test *p* = 3×10^−14^). No enoxolone treatment significantly altered RLU levels when compared to vehicle treatment. **(D)** Representative whole-mount images of 5 dpf *Fus(ApoBb.1-NanoLuciferase); Tg(ubi:mcherry-2A-FireflyLuciferase)* larvae following treatment of vehicle, 5 µM lomitapide, or 8 µM enoxolone for 48 hours. Lomitapide treatment induced a dark yolk phenotype, while no notable phenotypes followed enoxolone treatment. Scale bar represents 1 mm. **(E)** Representative image of a native-PAGE gel of luminescent B-lps from homogenates of 5 dpf animals treated with vehicle, 5 µM lomitapide, or 8 µM enoxolone for 48 hours. The image is a composite of chemiluminescence (B-lps, cyan hot) and fluorescence (DiI-LDL, yellow). For quantifications, B-lps were binned into one of 4 classes (ZM (zero mobility), very-low-density lipoproteins (VLDL), intermediate-density lipoproteins (IDL), or LDL), and these values were visualized via boxplot. The gel image is a representative image of representative samples from one of the three independent experiments performed. * <0.05 as compared to vehicle

Immersing a live, developing animal in a compound has the potential to profoundly alter overall tissue development that could secondarily result in attenuating B-lp levels. To address this concern, we examined the effect of enoxolone on overall gross larval morphology. Animals treated with 8 µM enoxolone are shorter than vehicle-treated animals, lack inflated swim bladders, and have mild cardiac edema (Figure 4D, Supplemental Figure 3). Whether the morphological phenotypes observed at 8 µM enoxolone resolve over time was not assessed here. However, the noted phenotypes are absent from 4 µM enoxolone treated animals, which appear strikingly similar to vehicle-treated animals. Treatment of larvae with 5 µM lomitapide (the positive control compound in our screen), a potent inhibitor of the lipid transfer activity of MTP, results in short animals and an observable phenotype that we have termed the ‘dark yolk’ (Figure 4D, Supplemental Figure 3). Typically, MTP protein in the yolk syncytium is responsible for packaging lipids from the yolk into B-lps destined for circulation [63]. When MTP activity is disrupted, yolk lipids are not secreted in B-lps and instead are stored in cytoplasmic lipid droplets within the yolk syncytium. These lipid droplets increase the opacity of the yolk, causing it to darken under transmitted light (Figure 4D). Notably, enoxolone does not cause a dark yolk phenotype.

We also measured the effect of drug treatment on lipoprotein classes via native-PAGE. While enoxolone treatment reduces total B-lps (Figure 4A and 4B), there are no overt changes in B-lp size distribution compared to vehicle-treated animals other than a slight increase in the zero mobility (ZM) fraction which contains very large particles and/or tissue aggregates (Figure 4E).

### Pharmacological inhibitors of HNF4⍺ reduce B-lp levels in the larval zebrafish

Enoxolone effectively lowers B-lp levels in larval zebrafish and appears to do so in a mechanism unlike that of lomitapide (Figure 4). Recent work has suggested that enoxolone may directly bind and inhibit the transcription factor HNF4⍺, contributing to changes in plasma lipids in the mouse [44]. To test if HNF4⍺ mediates the effect of enoxolone, we first evaluated whether pharmacological inhibition of HNF4⍺ alters B-lp levels. BIM5078 was recently defined as a novel and selective HNF4⍺ antagonist and was chemically modified to improve bioavailability (BI6015) [64]. Treatment (48 h) of animals (3 dpf) with 8 µM BI6015 reduced LipoGlo levels in fixed animals compared to their respective vehicle controls (Figure 5A). Total luminescence from whole-animal homogenates collected after treatment was also reduced following BIM5078 (8 µM, 4 µM, 1 µM, 0.5 µM, 0.25 µM, 0.125 µM, and 0.0625 µM) and BI6015 (8 µM, 4 µM, 2 µM, 1 µM, and 0.125 µM) treatments (Figure 5B). Importantly, neither BIM5078 nor BI6015 directly inhibit NanoLuciferase’s enzymatic activity (8 µM, 4 µM, and 0.05 µM; Figure 5C). We also assessed each compound’s effect on B-lp size distribution. Besides a small but significant increase in VLDL particles caused by BIM5078 treatment, BIM5078 and BI6015 do not cause any significant overall changes in B-lp size distributions (Figure 5D).

**Figure 5.**
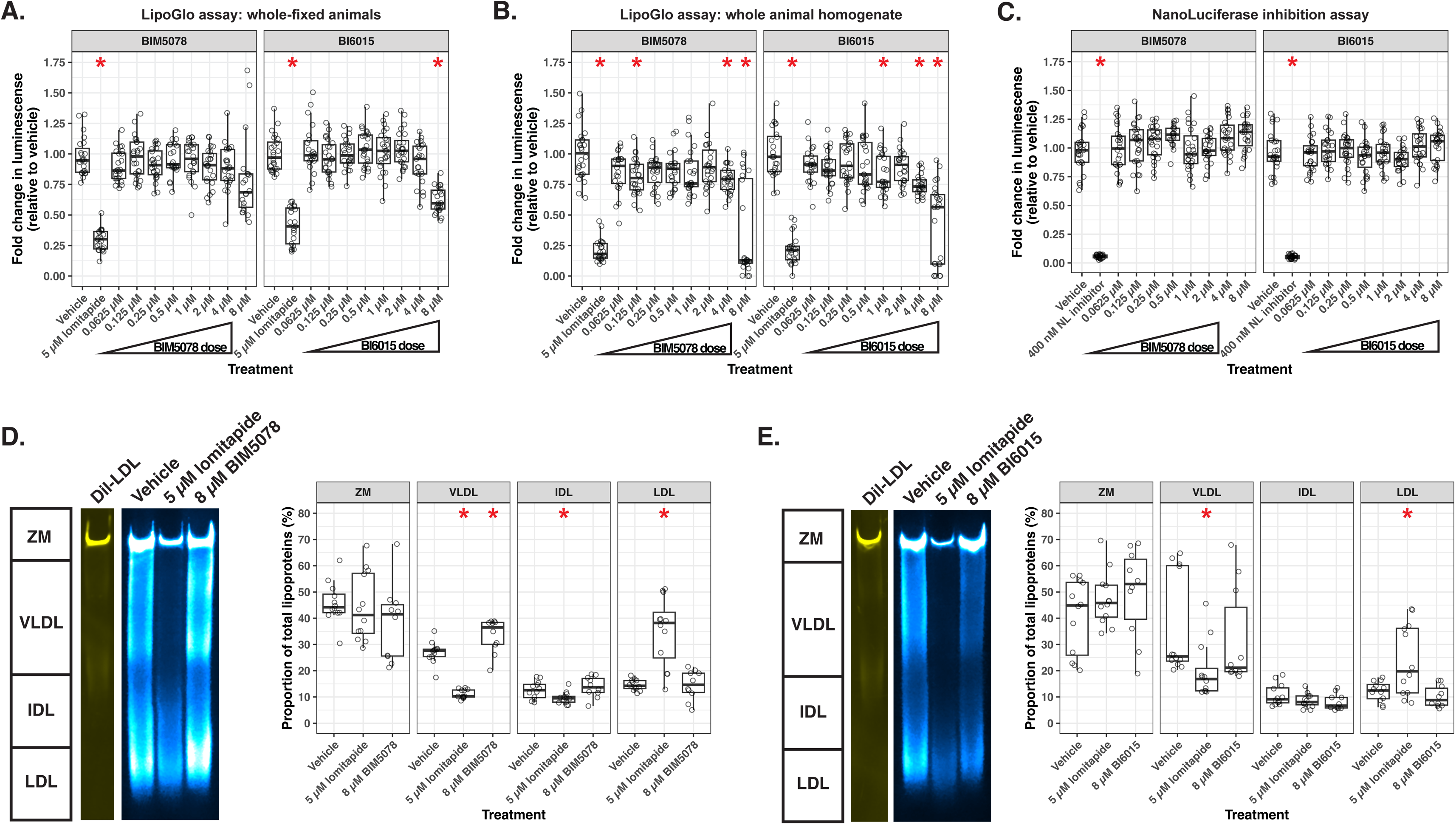
Pharmacological inhibition of HNF4⍺ reduces lipoproteins in the larval zebrafish. **(A)** Boxplot of the average fold change of Relative Luminescence Units (RLU) measured from fixed 5 dpf *Fus(ApoBb.1-NanoLuciferase); Tg(ubi:mcherry-2A-FireflyLuciferase)* animals treated for 48 hours with either negative (vehicle), positive (5 µM lomitapide) control, or an 8-fold serial dilution of BIM5078 or BI6015. Each data point represents a measurement from an independent animal collected from three independent experiments and normalized to the average vehicle RLU from each individual experiment. Several treatments significantly altered RLU levels in the BIM5078 experiment (one-way ANOVA, *F*(9,216) = 29.61, *p* < 2×10^−16^) and BI6015 experiment (one-way ANOVA, *F*(9,219) = 43.6, *p* < 2×10^−16^). In the BIM5078 experiment, only lomitapide treatment (n = 21) significantly reduced RLU levels compared to vehicle treatment (n = 23, Dunnett’s test *p* = 1.1×10^−18^). In the BI6015 experiment, lomitapide treatment (n = 21) significantly reduced RLU levels compared to vehicle treatment (n = 24, Dunnett’s test *p* = 8.3×10^−16^). Treatment with 8 µM BI6015 (n = 21, Dunnett’s test *p* = 6.9×10^−12^) also reduced total RLUs. **(B)** Boxplot of the average fold change of RLUs measured from homogenized 5 dpf *Fus(ApoBb.1-NanoLuciferase); Tg(ubi:mcherry-2A-FireflyLuciferase)* animals that were treated for 48 hours with either vehicle, 5 µM lomitapide, or an 8-fold serial dilution of BIM5078 or BI6015. Several treatments significantly altered RLU levels in the BIM5078 experiment (one-way ANOVA, *F*(9,221) = 39.61, *p* < 2×10^−16^) and BI6015 experiment (one-way ANOVA, *F*(9,223) = 42.7, *p* < 2×10^−16^). In the BIM5078 experiment, lomitapide treatment (n = 22) significantly reduced RLU levels compared to vehicle treatment (n = 24, Dunnett’s test *p* = 2×10^−16^). Treatment with 8 µM BIM5078 (n = 24, Dunnett’s test *p* = 9.1×10^−8^), 4 µM BIM5078 (n = 24, Dunnett’s test *p* = 1.2×10^−3^), and 0.125 µM BIM5078 (n = 23, Dunnett’s test *p* = 8.3×10^−3^) also reduce total RLUs. In the BI6015 experiment, lomitapide treatment (n = 22) significantly reduced RLU levels compared to vehicle treatment (n = 24, Dunnett’s test *p* = 2.3×10^−18^). Treatment with 8 µM BI6015 (n = 24, Dunnett’s test *p* = 8.8×10^−8^), 4 µM BI6015 (n = 23, Dunnett’s test *p* = 1.2×10^−5^), and 1 µM BI6015 (n = 24, Dunnett’s test *p* = 2.1×10^−2^) also reduce total RLUs. **(C)** Boxplot of the average fold change of RLUs measured from untreated homogenates of 5 dpf *Fus(ApoBb.1-NanoLuciferase); Tg(ubi:mcherry-2A-FireflyLuciferase)* animals that were briefly treated with either vehicle, 400 nM NanoLuciferase inhibitor, or an 8-fold serial dilution of BIM5078 or BI6015 to determine if HNF4⍺ inhibitors interfere with NanoLuciferase enzymatic activity. Several treatments significantly altered RLU levels in the BIM5078 experiment (one-way ANOVA, *F*(9,218) = 90.21, *p* < 2×10^−16^) and BI6015 experiment (one-way ANOVA, *F*(9,224) = 108.7, *p* < 2×10^−16^). In the BIM5078 experiment, only the NanoLuciferase inhibitor treatment (n = 22) significantly reduced RLU levels compared to vehicle treatment (n = 23, Dunnett’s test *p* = 1.4×10^−16^) and BIM5078 treatment did not alter RLU levels. In the BI6015 experiment, only the NanoLuciferase inhibitor treatment (n = 24) significantly reduced RLU levels compared to vehicle treatment (n = 23, Dunnett’s test *p* = 7.3×10^−17^) and BI6015 treatment did not alter RLU levels. **(D)** Representative image of a native-PAGE gel of luminescent B-lps from homogenates of 5 dpf animals treated with vehicle, 5 µM lomitapide, or 8 µM BIM5078 or 8 µM BI6015 for 48 hours. The image is a composite of chemiluminescence (B-lps, cyan hot) and fluorescence (DiI-LDL, yellow). For quantifications, B-lps were binned into one of 4 classes (ZM (zero mobility), very-low-density lipoproteins (VLDL), intermediate-density lipoproteins (IDL), or LDL), and these values were visualized via boxplot. The gel image is a representative image of representative samples from one of the two independent experiments performed.

### HNF4⍺ is required for normal B-lp levels in the larval zebrafish

We demonstrated that the pharmacological inhibition of HNF4⍺ is sufficient to reduce B-lp levels, suggesting this transcription factor is required to maintain normal B-lp levels in the fish. We next aimed to determine whether HNF4⍺ is genetically required for normal B-lp levels throughout larval development. We obtained a loss of function allele of HNF4⍺, *rdu14* [65], and crossed the allele into a *Fus(ApoBb.1-NanoLuciferase)/+* background to assess the effect of genetic loss of HNF4⍺ on B-lp quantity and size. We quantified total B-lp levels from 1 to 5 dpf from wild-type, heterozygous (*HNF4*⍺*^rdu14/+^*), and homozygous (*HNF4*⍺*^rdu14/rdu14^*) animals. In wild-type animals, the levels of B-lps rise as the animal utilizes the lipids found within the maternally deposited yolk (Figure 6A) [35]. As the yolk lipids are depleted, there is a concurrent decrease in total B-lp levels (Figure 6A). We found that homozygous animals have reduced B-lp levels at 1, 2, 3, and 4 dpf (Figure 6A). Further, we observed a minor but significant reduction of B-lp levels at 4 dpf in heterozygous animals (Figure 6A). By 5 dpf when yolk lipids are largely depleted, wild-type, heterozygous, and homozygous animals have similar B-lp levels (Figure 6A). Thus, HNF4⍺ expression is required for proper B-lp production.

**Figure 6.**
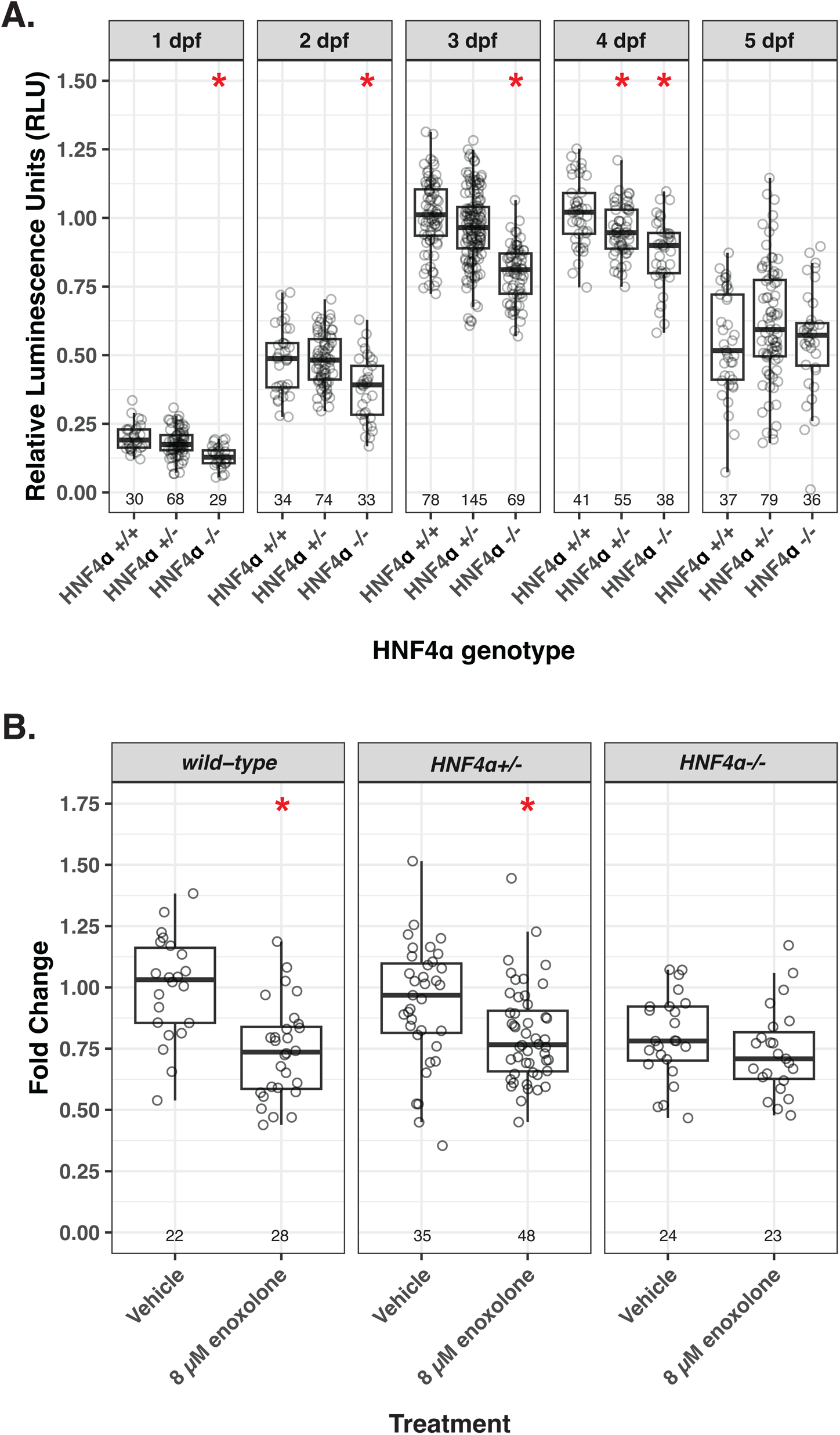
HNF4⍺ is required for lipoproteins throughout larval development and for the lipoprotein-reducing effect of enoxolone. **(A)** Boxplot of normalized Relative Luminescence Units (RLU) measured from homogenized *Fus(ApoBb.1-NanoLuciferase)/+* whole animals that were either *HNF4*⍺*^+/+^*, *HNF4*⍺*^rdu14/+^*, or *HNF4*⍺*^rdu14/rdu14^*, collected at 1, 2, 3, 4, and 5 dpf. Data were collected from at least three independent experiments and normalized to the mean of 3 dpf *HNF4*⍺*^+/+^* animals. Lipoprotein levels change throughout development (two-way ANOVA, *F*(1) = 261.206, *p* < 2×10^−16^) and due to the loss of HNF4⍺ (*F*(2) = 12.13, *p* = 6.4×10^−6^). Compared to their wild-type siblings, HNF4⍺ homozygotes have reduced lipoproteins at 1 dpf (Dunnett’s test, *HNF4*⍺*^rdu14/rdu14^* n = 29 versus *HNF4*⍺*^+/+^* n = 30, *p* = 1.6×10^−6^), 2 dpf (Dunnett’s test, *HNF4*⍺*^rdu14/rdu14^* n = 33 versus *HNF4*⍺*^+/+^* n = 34, *p* = 6×10^−3^), 3 dpf (Dunnett’s test, *HNF4*⍺*^rdu14/rdu14^* n = 69 versus *HNF4*⍺*^+/+^* n = 78, *p* = 1.8×10^−19^), 4 dpf (Dunnett’s test, *HNF4*⍺*^rdu14/rdu14^* n = 38 versus *HNF4*⍺*^+/+^* n = 41, *p* = 1.7×10^−5^). HNF4⍺ mutants have unchanged lipoproteins at 5 dpf. **(B)** Boxplot of normalized RLUs measured from homogenized *Fus(ApoBb.1-NanoLuciferase)/+* whole animals that were either wild-type, heterozygous, or homozygous (*HNF4*⍺*^+/+^*, *HNF4*⍺*^rdu14/+^*, or *HNF4*⍺*^rdu14/rdu14^* respectively) and treated with either vehicle or 8 µM enoxolone for 48 hours. The data were collected from two independent experiments and normalized to the mean of vehicle-treated *HNF4*⍺*^+/+^* animals. Lipoprotein levels were significantly altered by the HNF4⍺ genotype (Two-way ANOVA, *F*(2) = 3.385, *p* = 3.6×10^−2^), drug treatment (*F*(1) = 22.736, *p* = 3.9×10^−6^), and the interaction of the HNF4⍺ genotype and drug treatment (*F*(2) = 3.136, *p* = 4.6×10^−2^). Enoxolone treatment reduced lipoproteins in *HNF4*⍺*^+/+^* (Dunnett’s test, 8 µM enoxolone n = 28 versus vehicle n = 22, *p* = 1.7×10^−4^) and *HNF4*⍺*^rdu14/+^* (Dunnett’s test, 8 µM enoxolone n = 48 versus vehicle n = 35, *p* = 4.2×10^−2^). However, enoxolone treatment did not significantly alter lipoprotein levels in *HNF4*⍺*^rdu14/rdu14^* animals (Dunnett’s test, 8 µM enoxolone n = 23 versus vehicle n = 24, *p* = 0.8).

### HNF4⍺ is required for the B-lp-reducing effect of enoxolone treatment

Enoxolone reduces total B-lps when treating 3 dpf animals for 48 hours (Figure 4A and Figure 4B). We wanted to determine if this effect of enoxolone requires HNF4⍺. Thus, we treated a mixed population of wild-type, heterozygous, and homozygous 3 dpf animals with 8 µM enoxolone for 48 hours, homogenized individual animals, and quantified total B-lp levels (Figure 6B). Enoxolone treatment of wild-type and heterozygous animals resulted in a significant decrease in total B-lp levels. Notably, homozygous animals failed to respond to enoxolone treatment (Figure 6B), demonstrating that HNF4⍺ is required for the B-lp-reducing effect of enoxolone treatment.

### Enoxolone treatment alters expression patterns of the cholesterol biosynthetic pathway

To further characterize how enoxolone treatment modulates B-lp levels, we took an unbiased approach to identify the transcriptional changes following enoxolone treatment. We treated LipoGlo expressing animals with 8 µM enoxolone or vehicle for 4, 8, 12, 16, or 24 hours. Following each treatment duration, we collected total RNA from whole animals for differential expression analysis using RNAseq (Supplemental Tables 4 and 5). After 4 and 8 hours of enoxolone treatment, 39 and 34 genes were differentially expressed, respectively, with most genes upregulated (Figure 7A, Supplemental Table 5). Following 12, 16, and 24 hours of treatment, we identified 57, 118, and 402 differentially expressed genes, respectively (Figure 7A, Supplemental Table 5). The variations of these samples deviate from each other by both treatment and treatment duration, as shown by PCA, while biological replicate (n = 3 per treatment and treatment duration) variation remained low (Supplemental Figure 4A). These data indicate that enoxolone treatment induces a transcriptional response in animals as early as 4 hours post-treatment, and the breadth of that response increases over treatment duration.

**Figure 7.**
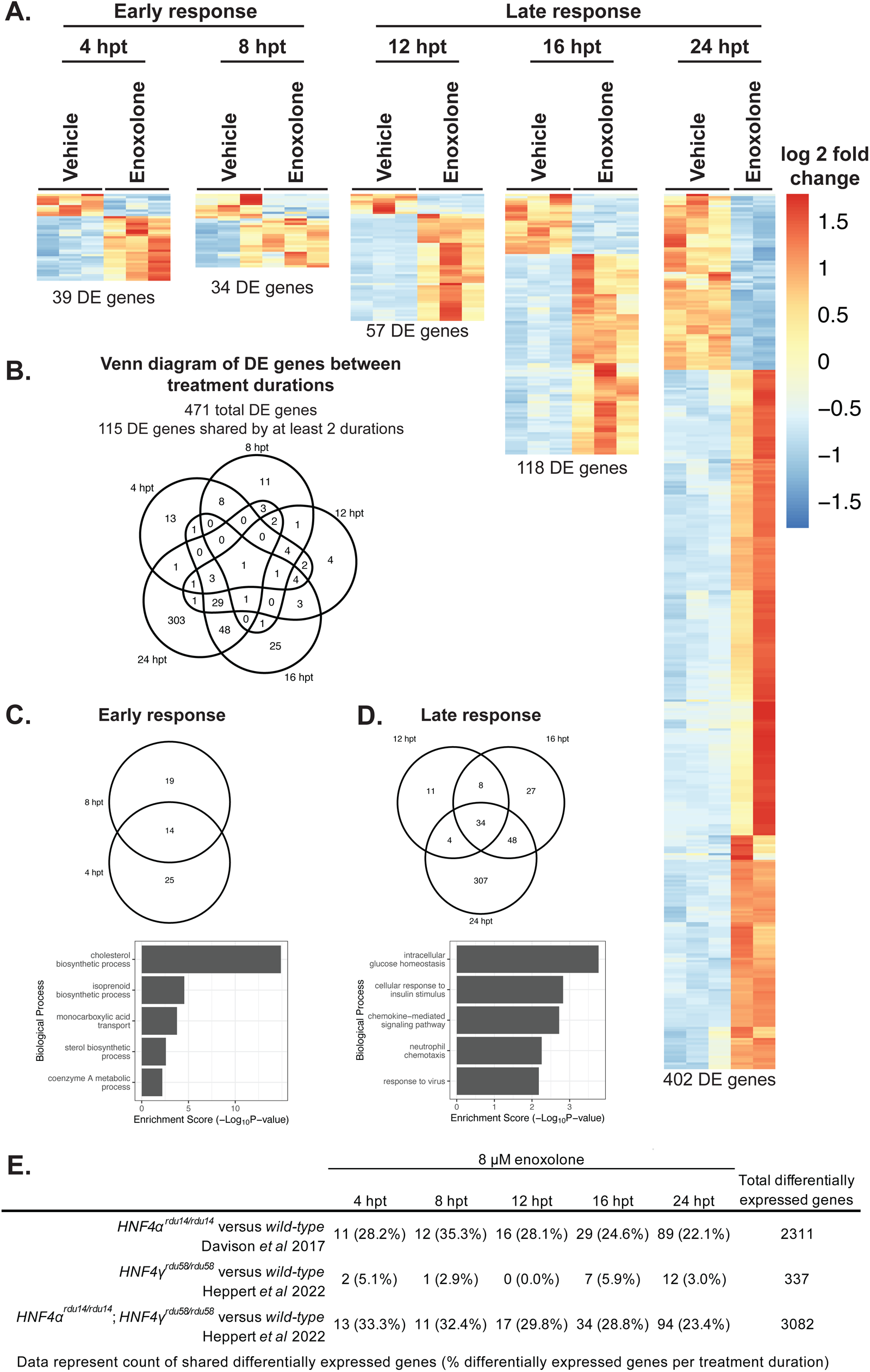
Differential expression analysis throughout enoxolone treatment affects lipid regulatory genes and is similar to the genetic loss of HNF4⍺. **(A)** Heat map of differentially expressed (DE) genes following 4, 8-, 12-, 16-, and 24-hours post-enoxolone treatment (hpt), respectively, 39, 34, 57, 118, and 402 genes were differentially expressed with red colors depicting increased and blue colors depicting decreased relative expression levels (log2 fold change). Each column of each heatmap represents a single replicate. **(B)** Venn diagram of overlapping differentially expressed genes from each treatment duration. Of the total 471 differentially expressed genes, 115 are shared between at least two treatment durations, and only one gene, *insig1*, is shared by all durations. **(C)** The early response to enoxolone treatment features 14 differentially expressed genes. Gene ontology analysis of these 14 genes reveals enrichment of lipid regulating pathways. **(D)** The late response to enoxolone treatment features 34 differentially expressed genes. Gene ontology analysis of these 34 genes reveals carbohydrate-regulating and cell signaling pathway enrichment. **(E)** Table comparing differentially expressed genes following 4, 8-, 12-, 16-, and 24-hours post-enoxolone treatment to HNF4⍺ knockout, HNF4Ɣ knockout, and HNF4⍺/HNF4Ɣ double knockout. There is considerable overlap between differentially expressed genes following enoxolone treatment and HNF4⍺ knockout, but little overlap with HNF4Ɣ knockout.

Of the 471 differentially expressed genes across all treatment durations, we found that 115 genes are shared among at least two durations, indicating overlap between these different treatment timepoints (Figure 7B). Interestingly, a single gene, *insulin induced gene 1* (*insig1*), was transcriptionally upregulated at all durations following enoxolone treatment (Figure 7B and Supplemental Figure 4B). *insig1* is a cholesterol sensor and critical regulator of lipid and carbohydrate metabolism [66,67]. We further characterized differentially expressed genes using gene ontology analysis at each treatment duration (Supplemental Figure 4C-4G, Supplemental Table 6) and by the early and late responses (Figure 7C and 7D). Fourteen genes were shared among the total 58 differentially expressed genes from 4 and 8 hours post-treatment, and these 14 genes are strongly associated with cholesterol biosynthesis and other lipid-associated pathways (Figure 7C, Supplemental Table 6).

Of the 439 differentially expressed genes from 12, 16, and 24 hours post-treatment, 34 differentially expressed genes are shared between all three treatment durations and are associated with gene ontology terms related to carbohydrate metabolism and signaling pathways (Figure 7D). We expanded this analysis using the bioinformatic tool Enrichr [68–70], which largely recapitulated gene ontology results described above. However, Enrichr analysis revealed significant overlap between several late enoxolone-responsive gene sets and transcriptional signatures associated with prochlorperazine, another compound identified in our screen (Figure 2, Figure 3, Supplemental Figure 1AO, Supplemental Figure 2Z). Enrichment of prochlorperazine-associated signatures was observed at 12 (prochlorperazine MCF7 up, 4/58 genes [INSIG1;IRF7;ISG15;ATF3], adjusted p-value = 0.006), 16 (prochlorperazine MCF7 up, 6/58 genes [INSIG1;DDIT4;IRF7;PMAIP1;ISG15;ATF3], adjusted p-value = 0.00008; prochlorperazine PC3 up, 3/29 genes [INSIG1;DDIT4;ATF3], adjusted p-value = 0.006), and 24 hours post-treatment (prochlorperazine PC3 up, 6/29 genes [DUSP5;INSIG1;DDIT3;TRIB3;SQSTM1;ATF3], adjusted p-value = 0.0001; prochlorperazine MCF7 up, 7/58 genes [DDIT3;INSIG1;IRF7;PMAIP1;ISG15;SAT1;ATF3], adjusted p-value = 0.0005). These results suggest that enoxolone and prochlorperazine may perturb overlapping molecular pathways, an observation that warrants further investigation. Ultimately, these data demonstrate distinct early and late responses to enoxolone treatment, and the early response modulates key lipid metabolism pathways.

### Enoxolone treatment causes changes in expression patterns similar to HNF4⍺ knockout

The roles of HNF4⍺ in larval zebrafish development have recently been characterized [65,71]. HNF4⍺ loss is viable in the fish, likely due to partially redundant function with HNF4β and HNF4Ɣ [71]. Interestingly, the transcriptional targets of HNF4⍺, HNF4β, and HNF4Ɣ only partially overlap, indicating that each transcription factor has specific and non-specific molecular targets. Thus, we hypothesized that if the molecular target of enoxolone is HNF4⍺, then enoxolone treatment would phenocopy the effect of HNF4⍺ knockout. We compared the differentially expressed genes following 4, 8, 12, 16, and 24 hours after enoxolone treatment to the subset of differentially expressed genes identified from intestines extracted from 6 dpf HNF4⍺ homozygous (relative to *wild-type* controls) animals (Figure 7E, Supplemental Table 7) [71]. Of all differentially expressed genes for each enoxolone treatment duration, 22.1% to 35.3% were shared with differentially expressed genes from the HNF4⍺ knockouts (Figure 7E). To assess the specificity of this comparison, we also compared differentially expressed genes for each enoxolone treatment duration to the subset of differentially expressed genes identified from intestines extracted from 6 dpf HNF4ɣ knockouts (*HNF4ɣ^rdu58/rdu58^*; relative to *wild-type* controls) animals (Figure 7E). Importantly, the comparison of overlapping differentially expressed genes between enoxolone treatment and HNF4ɣ knockouts was substantially lower than the comparison to HNF4⍺ knockouts, with between 0.0%-5.9% of enoxolone differentially expressed genes shared with HNF4ɣ knockouts (Figure 7E). Further demonstrating the specificity of significant overlap between enoxolone treatment and HNF4⍺ knockout, we additionally found substantial overlap of differentially expressed genes between enoxolone treatment and HNF4⍺ HNF4ɣ double knockouts (23.4%-33.3%; Figure 7E). Ultimately, these data support that enoxolone treatment, at least partially, phenocopies the effect of HNF4⍺ knockout.

## Discussion

Currently, the only known means to improve the outcomes of hyperlipidemia and reduce the risk of CVD-associated mortality is to reduce circulating levels of B-lps. The two major strategies for reducing B-lp levels are blocking B-lp synthesis (e.g., lomitapide and mipomersen) or promoting circulating B-lp uptake (e.g., statins, bempedoic acid, and PCSK9 targeting therapies). Continual statin use for more than 1 year in individuals with high B-lp levels reduced circulating B-lps and lowered all-cause mortality by ∼30% [12,13]. However, that still leaves significant mortality in people taking statins. While newer alternatives to statins are effective at reducing B-lp levels, they are costly and feature some undesirable off-target effects, limiting their use to the most severe forms of dyslipidemia, and their long-term outcomes have yet to be fully determined [25–28]. Thus, there is still a need to identify new mechanisms for reducing B-lps, thereby improving CVD therapeutics.

Recent work from our lab identified a new hypomorphic allele of MTP, reducing the lipid transfer activity of triglyceride but not phospholipid from the endoplasmic reticulum [63]. Animals expressing this mutant MTP synthesize triglyceride-poor B-lps and are phenotypically normal in adulthood, unlike animals carrying a loss of function MTP mutation. The revelation that MTP lipid transfer activities can be modulated demonstrates the potential for developing a selective small molecule triglyceride transfer inhibitor to reduce B-lps lipid content. Further, B-lps are regulated by many different mechanisms and cell types, calling into question why B-lp lowering therapies are still limited to targeting B-lp synthesis and uptake. Consistent with the goal of finding new targets to reduce plasma B-lps, recent efforts have identified a number of new genes that regulate B-lps in the larval zebrafish [72,73]. Thus, our understanding of B-lp regulation is not yet saturated, and we expect many druggable targets in the pathways of B-lp regulation that have yet to be defined.

We aimed to identify new strategies for lowering circulating B-lp levels using drug screening and directly measuring total B-lp levels in a whole-animal system. While limited, there have been incremental advances in identifying B-lp modulators in complex systems. Prior work from the Hekimi lab screened novel compounds that affect a slow defecation phenotype that correlates with lowering B-lp levels in *C. elegans* and identified one highly effective molecule, CHGN005 [74]. More recently, the Duncan lab screened for ApoB-lowering compounds using iPSC-derived hepatocytes, identifying 46 ApoB-lowering compounds and confirming one, DL-1, in rodent models [75]. However, until now, there have not been any large-scale drug screens to identify B-lp modulators in a vertebrate whole-animal system.

Using zebrafish expressing the LipoGlo reporter, we examined the effects of 2762 compounds of the JHDL to identify new B-lp lowering drugs. An obstacle in drug screening is defining significant compounds or hits, especially considering the variance of biological replicates [52]. Researchers commonly use arbitrarily defined cutoffs to identify the largest reasonable number of significant screen hits [52,76]. To identify significant B-lp-reducing compounds, we used a well-described statistical measure, SSMD, and considered the directly measured effect size of B-lp reduction (Figure 1F). Using these methods, we identified 49 unique B-lp-reducing compounds. With our screening paradigm, we frequently examined our positive control (n = 1381), which consistently reduced B-lp levels. However, when we applied our hit cutoffs, we found that lomitapide treatment infrequently would have been called a hit (26.4%), suggesting that our hit cutoffs are very stringent. Further demonstrating this stringency, if we performed a Student’s t-test for each 5 µM lomitapide treatment we examined (n = 1381), 81.7% of experiments (1129/1381) would have reached statistical significance (p < 0.05). Rank-ordered analysis of lomitapide fold change and SSMD defines the midpoints of these measurements as -1.33 and -1.41, respectively. We found that using this approach to define hit cut-offs increases the frequency that lomitapide treatments are called as a hit increases (34.4%), and the number of B-lp lowering compounds increases to 59 unique drugs (12 additional compounds that are the subject of ongoing studies). Further, we note that we did not independently re-test compounds falling below our SSMD and fold-change cutoffs, thus a systematic assessment of the false-negative rate of our screening paradigm awaits future work. As many of our hit compounds validated, our approach to use the positive control in a drug screen to help define hit cutoffs is advantageous. However, this approach is possible only when the positive control is utilized frequently (e.g. on every plate) throughout the screening paradigm.

Our screen identified numerous B-lp-increasing compounds in addition to B-lp-reducing compounds. While each of these compounds may not be clinically advantageous, their mechanisms of action may define novel biological pathways that may provide new therapeutic targets to manipulate B-lps *in vivo.* When further examining B-lp-increasing compounds, we must be careful to consider that B-lp levels change throughout larval zebrafish development [35]. Thus, the B-lp increasing effect of these compounds may act indirectly on B-lps as a result of the drug slowing larval development, delaying the eventual decrease in B-lps that happens over development time – a hypothesis we have yet to examine for each of these B-lps-increasing compounds.

We confirmed the B-lp-lowering effects of 19 hits from our screen with several orthogonal studies. Further validation of these hits demonstrated a wide range of potential mechanisms of lipoprotein regulation. We identified hits that affected larval development, some hits that reduced total B-lp levels, and several structurally related compounds that directly reduced B-lp particle size (Figure 3). We then reviewed the literature for each of these compounds and prioritized hits for further study based on their likelihood to modulate B-lp levels. We selected the licorice root derivative enoxolone, because many compounds from this plant are used in traditional Chinese medicine to treat a myriad of disorders, including non-alcoholic fatty liver disease, dyslipidemia, and obesity [45,57,77–79]. However, mechanistic insight into how licorice root and its components produce these clinically favorable outcomes is largely lacking.

While enoxolone reduces B-lp levels, its effects on larval morphology and B-lp profile differ from that of MTP inhibition. More specifically, treatment of larvae with the MTP inhibitor, lomitapide, causes lipid droplet accumulation in the yolk syncytium, resulting in a ‘dark yolk’ phenotype (Figure 4D). While the highest concentration of enoxolone reduced larval standard length, no concentration of enoxolone tested produced a dark yolk. Further, lomitapide treatment results in only small B-lps [35], while enoxolone treatment does not affect the profile of B-lp size (Figure 4E). These data demonstrate that enoxolone reduces B-lp levels by a mechanism distinct from MTP inhibition.

Prior studies of enoxolone have found it to be protective from high-fat diet-induced hyperlipidemia and alcohol-induced hepatic injury [44,45]. However, it has also been found that enoxolone treatment is associated with inhibition of 11β-hydroxysteroid dehydrogenase activity, which can promote hypertension in some patients [43]. While this effect reduces enthusiasm for the clinical use of enoxolone, it may be possible to separate the B-lp lowering and hypertensive mechanisms of enoxolone by altering its structure.

Recent studies found enoxolone was capable of inhibiting HNF4⍺-dependent transcription of genes necessary for B-lp biosynthesis in cells and mice and protected animals from high-fat diet-induced hyperlipidemia [44]. Further, enoxolone docking analysis suggests that this drug interacts with HNF4⍺, which is a potent lipid and carbohydrate metabolism regulator in the liver and gut [80–83]. However, those authors could not directly show that HNF4⍺ was required to suppress hyperlipidemia in an animal model [44]. Here, we demonstrate that HNF4⍺ is required for the B-lp lowering effect of enoxolone and suggest that enoxolone directly targets HNF4⍺ or is a potent modulator of HNF4⍺ activity.

HNF4⍺ regulates many transcriptional targets [71,80]. Thus, we performed differential expression analysis to define the altered transcriptional pathways following enoxolone treatment. The early response to enoxolone treatment is enriched in differentially expressed genes related to lipid regulatory processes like cholesterol and isoprenoid biosynthesis, lipid transport, and the metabolism of coenzyme A (Figure 7C). Interestingly, many differentially expressed genes enriched at times immediately after treatment do not overlap with late response genes, and these late genes are associated with glucose/insulin signaling and other cellular pathways. However, one gene, *insig1*, is transcriptionally upregulated at every treatment duration measured (Supplemental Figure 4B). Insulin-induced gene 1 (Insig1) is a cellular cholesterol sensor that controls fatty acid and cholesterol biosynthesis by regulating the activity of sterol regulatory element binding proteins (SREBPs) and levels of HMG CoA reductase protein [84,85]. Not surprisingly, insig1 is closely correlated to dyslipidemia and other metabolic disorders, and Insig1 targeting therapies have been proposed [66,84,85]. We expect that *insig1* upregulation following enoxolone treatment signifies a response to the limited cholesterol mobilization associated with reduced B-lp levels following treatment.

Several transcriptional targets of enoxolone have recently been described; for example, enoxolone treatment downregulates the expression of genes key to B-lp synthesis, including MTP, ApoB, and PLA2G12B [44,86,87]. Interestingly, we do not detect any changes in expression for MTP, ApoB, and PLA2G12B following enoxolone treatment (Supplemental Table 5). However, it is possible that enoxolone would affect the expression of these genes in a fish model of hyperlipidemia.

We also examined whether enoxolone treatment in the fish causes transcriptional changes similar to the genetic depletion of HNF4⍺. The transcriptional targets of HNF4⍺ and HNF4Ɣ in the larval zebrafish intestine were recently characterized [71]. Many differentially expressed genes following enoxolone treatment were also differentially expressed in the intestines of HNF4⍺ knockout animals (22.1% to 35.3%, Figure 7E). This similarity is lost when comparing HNF4Ɣ knockouts and enoxolone treatment. These data demonstrate that enoxolone treatment affects many of the transcriptional targets of HNF4⍺. It should be noted that these datasets do not entirely overlap; we hypothesize this may stem from differences between whole-animal enoxolone treatments and intestines from HNF4⍺ knockouts. Alternatively, the perturbation of the HNF4⍺-dependent transcriptional network may be one of several downstream effects of enoxolone treatment. However, this hypothesis is not well supported by the consistency of overlapping differentially expressed genes between different durations of enoxolone treatment. Ultimately, these data demonstrate that enoxolone treatment causes many transcriptional changes, and these changes phenocopy the effect of genetic loss of HNF4⍺, supporting that HNF4⍺ is a target of enoxolone treatment.

Here, we not only demonstrate that enoxolone reduces B-lp levels in the larval zebrafish through inhibition of HNF4⍺ activity but also show that phenotypic drug screens for compounds that alter lipid biology using a whole animal model is achievable. The conservation of enoxolone’s HNF4⍺ inhibition in the zebrafish provides strong proof of concept that our screen was successful. We also identified and confirmed many other B-lp-lowering compounds for which we have yet to define their mechanisms of action. Further, the drug screening paradigm we developed using the LipoGlo system is highly scalable and can be deployed to screen large novel drug libraries to identify many additional B-lp-lowering compounds.

## Materials and Methods

### Zebrafish husbandry and maintenance

All Zebrafish (*Danio rerio*) protocols were approved by the Carnegie Institution Department of Embryology Animal Care and Use Committee (Protocol #139). Adult zebrafish were maintained at 27°C on a 14:10 h light:dark cycle and fed once daily with ∼3.5% body weight Gemma Micro 500 (Skretting). Embryos were obtained by natural spawning, raised in embryo medium at 28.5°C, and kept on a 14:10 h light:dark cycle. All embryos used for experiments were obtained from pair-wise crosses and were staged according to [88]. Exogenous food was provided starting at 5.5 days post fertilization (dpf). Larvae were fed with GEMMA Micro 75 (Skretting) 3x a day until 14 dpf, GEMMA Micro 150 3x a day + Artemia 1x daily from 15 dpf–42 dpf and then GEMMA Micro 500 daily supplemented once a week with Artemia. The nutritional content of GEMMA Micro is as follows: Protein 59%; Lipids 14%; Fiber 0.2%; Ash 14%; Phosphorus 1.3%; Calcium 1.5%; Sodium 0.7%; Vitamin A 23000 IU/kg; Vitamin D3 2800 IU/ kg; Vitamin C 1000 mg/kg; Vitamin E 400 mg/kg. Zebrafish sex is not determined until the juvenile stage, so sex is not a variable in experiments with embryos and larvae. All larvae were maintained in E2 (15 mM NaCl/0.5 mM KCl/1 mM CaCl/0.15 mM KH_2_PO_4_/0.05 mM Na_2_HPO_4_/1 mM MgSO_4_) or E3 (5 mM NaCl/0.17 mM KCl/0.33 mM CaCl/0.33 mM MgSO_4_) embryo media as noted.

### Drug screen

The Johns Hopkins Drug Library (JHDL) (2934 total compounds supplied at 10 mM in DMSO) [89] was screened across a four-fold serial dilution (8 µM to 1 µM). Drug dilutions were prepared using the Automated Reporter Quantification *in vivo* (ARQiv) screening system [49]. Briefly, each JHDL compound was prediluted to a concentration of 160 µM in a 2% DMSO/100 ppm Tween-20/E3 solution to make a working drug stock. Each 96-well drug treatment plate (Perkin Elmer 6005299) was pre-loaded 225 µL 100 ppm Tween-20/E3 in columns 3 and 7, while the remaining columns were pre-loaded with 125 µL 0.2% DMSO/100 ppm Tween-20/E3 using a Micro10x microplate dispenser (Hudson Robotics). A 25 µL aliquot of drug working stock was added to each well of either column 3 or 7 of the microplate, mixed, and serially diluted by a SOLO automated pipettor (Hudson Robotics). Each drug treatment plate was prepared with a positive control in each well of column 2 with a concentration of 10 µM lomitapide (Aegerion Pharmaceuticals, #AEGR-733)/0.2% DMSO/100 ppm Tween-20/E3. To prepare for the addition of 3 dpf larvae, 75 µL of E3 was added to each well of each drug treatment plate.

Concurrently, *Fus(ApoBb.1-NanoLuciferase); Tg(ubi:mCherry-2A-Firefly Luciferase)* embryos were collected from homozygous in-crosses, manually cleaned daily, and reared in E2 at 28.5°C and kept on a 14:10 h light:dark cycle until 3 dpf. Once 3 dpf, animals were transferred to E3 and loaded into each well of each drug treatment plate in 50 µL using the Complex Object Parametric Analyzer and Sorter (COPAS-XL, Union Biometrica). The final concentration of each well of every drug treatment plate was as follows:

Wells of column 1: 1 fish per well in 250 µL 0.1% DMSO/50 ppm tween-20/E3

Wells of column 2: 1 fish per well in 250 µL 5 µM lomitapide/0.1% DMSO/50 ppm tween-20/E3 (positive control)

Wells of column 3: 1 fish per well in 250 µL 8 µM drug A/0.1% DMSO/50 ppm tween-20/E3

Wells of column 4: 1 fish per well in 250 µL 4 µM drug A/0.1% DMSO/50 ppm tween-20/E3

Wells of column 5: 1 fish per well in 250 µL 2 µM drug A/0.1% DMSO/50 ppm tween-20/E3

Wells of column 6: 1 fish per well in 250 µL 1 µM drug A/0.1% DMSO/50 ppm tween-20/E3

Wells of column 7: 1 fish per well in 250 µL 8 µM drug B/0.1% DMSO/50 ppm tween-20/E3

Wells of column 8: 1 fish per well in 250 µL 4 µM drug B/0.1% DMSO/50 ppm tween-20/E3

Wells of column 9: 1 fish per well in 250 µL 2 µM drug B/0.1% DMSO/50 ppm tween-20/E3

Wells of column 10: 1 fish per well in 250 µL 1 µM drug B/0.1% DMSO/50 ppm tween-20/E3

Wells of column 11: 1 fish per well in 250 µL 0.1% DMSO/50 ppm tween-20/E3 (negative control)

Wells of column 12: 1 fish per well in 250 µL 0.1% DMSO/50 ppm tween-20/E3

After dispensing, animals in treatment were incubated at 27°C and kept on a 14:10 h light:dark cycle for 48 hours. At 5 dpf, a solution of anesthetic, fixative, and NanoGlo substrate (Promega N1110) was added to each well of each drug plate at a final concentration of 0.014% MESAB/4.6% paraformaldehyde (Electron Microscopy Sciences)/12.2% NanoGlo Buffer/1% NanoGlo substrate. After a roughly 20-minute incubation, total well luminescence was measured using a Tecan M1000 Pro microplate reader with a 50 ms integration.

### LipoGlo assays with fixation endpoint

Each hit compound identified from our high-throughput screen was obtained from a vendor (Supplemental Table 3) and subjected to validation LipoGlo assays. In addition, BI6015 (Cayman Chemical 12032) and BIM5078 (Cayman Chemical 12031) were subjected to validation LipoGlo assays with the fixation endpoint. Validation assays were prepared as described above for high-throughput screening with a wider range of concentrations (eight-point serial dilution ranging from 8 µM to 62.5 nM) and using E2 embryo media. However, robotics were not used to prepare validation LipoGlo assays, and each experiment was prepared manually. After the 48-hour drug treatment, animals were fixed, and luminescence was measured using a BioTek Synergy H1 microplate reader with 500 ms integration or a Tecan Spark microplate reader with 500 ms integration. Each compound was examined with three independent experiments, with the results of each experiment normalized to the average RLU of the negative control (fold change) to control for experiment-to-experiment variation.

### LipoGlo assays with homogenization endpoint

Drug treatments of each hit compound, BI6015, and BIM5078 were prepared as described for LipoGlo assays with fixation endpoint. After the 48-hour drug treatment, animals were anesthetized and transferred to 96-well PCR plates (USA Scientific 1402-9598) in a homogenization buffer (10% sucrose/50 mM EGTA/1X cOmplete EDTA-free protease inhibitor cocktail tablet (Roche 11873580001)). Samples were homogenized via bath sonication in a shoehorn sonicator with an amplitude of 100, cycling pulses of 2 seconds on and 1 second off for a total cycle of 30 seconds. For each sample, an aliquot of 4 µL of homogenate was mixed with 66 µL phosphate-buffered saline (PBS), 10 µL NanoGlo buffer, and 0.2 µL NanoGlo substrate (Promega N1110) and incubated at room temperature for 5 minutes. Total luminescence was measured using a BioTek Synergy H1 microplate reader with 20 ms integration or a Tecan Spark microplate reader with 20 ms integration. Each compound was examined with three independent experiments, with the results of each experiment normalized to the average RLU of the negative control (fold change) to control for experiment-to-experiment variation.

### NanoLuciferase activity assay

We tested whether each hit compound, BI6015, and BIM5078 interfered with NanoLuciferase enzymatic activity. Untreated 3 dpf *Fus(ApoBb.1-NanoLuciferase); Tg(ubi:mCherry-2A-Firefly Luciferase)* animals were homogenized as described above. Here, 4 µL of homogenate was briefly incubated with each drug treatment (eight-fold serial dilution ranging from 8 µM to 62.5 nM) in a 0.1% DMSO/50 ppm tween-20/E2 solution in 96-well plates (Perkin Elmer 6005299). A solution of 30 µL PBS, 10 µL NanoGlo buffer, and 0.2 µL NanoGlo substrate (Promega N1110) was added to each sample and incubated at room temperature for 5 minutes. Total luminescence was measured using a BioTek Synergy H1 microplate reader with 20 ms integration or a Tecan Spark microplate reader with 20 ms integration. Each compound was examined with three independent experiments, with the results of each experiment normalized to the average RLU of the negative control (fold change) to control for experiment-to-experiment variation.

### Whole-mount bright-field microscopy and standard-length quantification

Fish were anesthetized following vehicle or drug treatment or at specific developmental points for microscopy. Individual animals were mounted in a solution of 3% methylcellulose and imaged using a Nikon SMZ1500 microscope with HR Plan Apo 1x WD 54 objective, Infinity 3 Lumenera camera, and Infinity Analyze 6.5 software. Images were rotated and cropped as necessary using FIJI (ImageJ V2.0.0, National Institutes of Health (NIH)). The standard length of individuals was measured in FIJI.

### Quantification of lipoprotein size distribution with LipoGlo electrophoresis

Native-PAGE gels (3% acrylamide:bis-solution 19:1 (BioRad, 1610144)/0.08% ammonium persulfate/0.006% tetramethylethylenediamine (TEMED)/1X TBE) were prepared for individual experiments. Gels were assembled into mini-protean electrophoresis rigs (BioRad) at 4 °C, filled with pre-chilled 1X TBE, and pre-run at 50V for 30 minutes to equilibrate the gel before sample addition. Twelve microliters of homogenate (prepared as described above) was then combined with 3 μL of 5X loading dye (40% sucrose/0.25% bromophenol blue/TBE), and 12.5 μL of the resulting solution was loaded per well (which corresponds to 10% of the larval homogenate per lane). A reference sample of DiI-labeled human LDL (Thermo Fisher Scientific, L3482) was mixed with 5X loading dye and loaded on each gel. Gels were then run at 50 V for 30 min, followed by 125 V for 2 h. Following resolution, gels were briefly coated in a 0.2% NanoLuciferase substrate/TBE solution or directly soaked in a 0.01% NanoLuciferase substrate/TBE solution before imaging using a LI-COR Odyssey Fc imaging system. The chemiluminescent channel (NanoLuciferase detection) was imaged for 2 minutes, and the 600 nm channel (DiI-LDL detection) was imaged for 30 seconds. Raw images were exported for further analysis using FIJI (ImageJ V2.0.0, National Institutes of Health (NIH)) and Excel (Microsoft). Briefly, each lane on the gel was converted to a plot profile and divided into LDL, IDL, VLDL and Zero Mobility (ZM) bins based on migration relative to the Di-I LDL standard [35]. Pixel intensity from the plot profile was summed within each bin for comparison between genotypes.

### Characterizing lipoproteins in HNF4⍺ mutants

To determine whether HNF4⍺ is required for normal lipoprotein levels, the *HNF4*⍺ allele *rdu14* was obtained from the Rawls laboratory [71] and crossed into the LipoGlo background. We then crossed *HNF4*⍺*^rdu14/+^; Fus(ApoBb.1-NanoLuciferase)/+* to *HNF4*⍺*^rdu14/+^* adults in pairs and collected embryos. Healthy embryos were maintained at 27°C on a 14:10 h light:dark cycle in E2 embryo media in 10-cm dishes in groups of 100 animals. At 1, 2, 3, 4, and 5 dpf, a dish of animals was anesthetized, and individual animals were homogenized as described above. Homogenates were subjected to LipoGlo analysis as described above. As only half of our animals are *Fus(ApoBb.1-NanoLuciferase)/+,* we defined heterozygous animals as those samples with luminescence above the background and wild-type animals as those samples with luminescence below the background. We genotyped for the rdu14 allele to de-identify samples. Briefly, genomic DNA was extracted in 50 mM sodium hydroxide from 10 µL of homogenate and finally buffered in 166 mM Tris pH 8.0. The locus containing the rdu14 allele was amplified by PCR, using the following primers (Forward: 5’-TGATTCACACTACTTACTTGTCTAG-3’, Reverse: 5’-GATTAAAAGTAGTTATCTCATCCTCAG-3’) and GoTaq polymerase (Promega M3001). The genotype of the locus was scored after resolving samples on 2.5% agarose gels as HNF4⍺*^+/+^, HNF4*⍺*^rdu14/+^, HNF4*⍺*^rdu14/rdu14^.* Samples were collected from at least 3 independent experiments, and each experiment was normalized to the mean of 3 dpf *HNF4*⍺*^+/+^; Fus(ApoBb.1-NanoLuciferase)/+* animals, as this time point was common across all experiments.

### RNA extraction, RNAseq, and downstream analysis

Individual 3 dpf *Fus(ApoBb.1-NanoLuciferase); Tg(ubi:mCherry-2A-Firefly Luciferase)* larvae were incubated in either vehicle (0.1% DMSO/50 ppm tween-20/E2) or 8 µM enoxolone (8 µM enoxolone/0.1% DMSO/50 ppm tween-20/E2). After 4, 8, 12, 16, or 24 hours, 5 larvae from each treatment were collected and pooled for RNA extraction. After 48 hours, total lipoproteins were measured from the remaining animals from each treatment duration, following the LipoGlo assay with the fixation endpoint described above, to confirm the efficacy of enoxolone treatment in the experiment. Samples were prepared and collected from three independent clutches.

Grouped samples were homogenized using 0.5 mm zirconium oxide beads (Next Advance, ZrOB05) with a Bullet Blender (Next Advance) in TRIzol reagent (Thermo Fisher, 15596026). Homogenates were mixed with 100% ethanol 1:1, and RNA was extracted using the Direct-zol RNA Microprep Kit (Zymo Research, R2060). RNA concentration and quality were assessed via the Thermo Scientific NanoDrop One and the Agilent 2100 Bioanalyzer. RNASeq libraries were prepared from approximately 500ng total RNA using the TruSeq Stranded mRNA Library Prep kit (Illumina, 20020595) and TruSeq RNA CD Index plate (Illumina, 20019792), according to the manufacturer’s protocol. Sequencing was done on the Illumina NextSeq 500, using a 75bp run with 8×8 indexing, yielding between 23 and 45 million reads per sample. RNA sequencing data were processed using the nf–core/rnaseq v3.11.1 pipeline [90]. Briefly, adapter sequences were trimmed from the reads, and ribosomal RNA reads were filtered. The remaining reads were mapped to the GRCz11 genome with Ensembl 110 annotation. Differential expression was examined using a custom R pipeline and the DEseq2 1.42 package for R. Each treatment duration was analyzed as an independent experiment to robustly analyze the difference between early and late effectors of enoxolone treatment. Samples from 8 hours post-treatment were run through a batch correction algorithm using the sva package [91]. Differentially expressed genes were identified as those with a fold change (log_2_ scale) beyond 1.5 (0.9 for 4 and 8 hours post-treatment) and adjusted p-value < 0.05. Gene ontology analysis was performed using the topGO package.

### Quantification and statistical analysis

All data processing and analyses were completed using custom scripts using R; see supplemental file 1 for all code used. Drug screen data were analyzed in near real-time, calculating the fold change and the Strictly Standardized Mean Difference (SSMD) [52] of each drug treatment compared to the negative control. Every drug-dose concentration that resulted in an SSMD score of ≤ -1.0 and fold change (log2 scale) of ≤ -1.5 was considered a significant luminescence-reducing hit. Statistically significant differences between drug treatments in the remaining experiments were identified by a one-way analysis of variance (ANOVA) test followed by a Dunnett’s test (with the negative control as the reference group) with a Bonferroni correction for multiple comparisons.

## Acknowledgments

We thank Dr. Jun Liu for contributing the JHDL for drug screening. We thank James Dweck, Jake Griffin, Kobe Koren, Urmi Kumar, Katelyn Macholl, Shajae Pinnock, and Maxwell Veiga for their help in animal husbandry and embryo cleaning for the drug screen. We thank Allison Pinder for library preparation and sequencing and Dr. Frederick Tan for pipelines for sequencing analysis via nf–core. We thank the members of Dr. John Rawls’ lab for their input of our differential expression analysis. We also thank the members of Dr. Steve Farber’s lab, especially Dr. Meredith Wilson for her editing of this manuscript. Support was provided by the National Institutes of Health (R01DK116079 [S.A.F.] and F32DK126297 [D.J.K.]). The Carnegie Institution for Science endowment and the G. Harold and Leila Y. Mathers Charitable Foundation (S.A.F) provided additional support for this work. This content is solely the responsibility of the authors and does not necessarily represent the official views of NIH.

---

**Supplemental Figure 1. Forty-nine unique compounds reduce B-lp levels in a high-throughput drug screen to identify modulators of B-lps in larval zebrafish.** Boxplots of each of the 50 (49 unique) B-lp lowering compounds from the initial drug screen of 2762 compounds, hits are compounds defined as having at least one dose test result in a fold change (log2 scale) of ≤ -1.5 and strictly standardized mean difference (SSMD) ≤ -1.0. Open circles represent each sample; sample size (n) and SSMD scores are listed below each treatment. The B-lp reducing hits are **(A)** unknown, **(B)** 3-methylcholanthrene, **(C)** acetaminophen, **(D)** bismuth (III) oxychloride, **(E)** cetrimonium bromide, **(F)** cyproterone, **(G)** cytochalasin N, **(H)** calcipotriene, **(I)** cinnamon oil, **(J)** cupric chloride, **(K)** cytidine 5’-monophosphate, **(L)** cytidine 5’-diphosphate trisodium salt, **(M)** danazol, **(N)** danthron, **(O)** demecarium bromide, **(P)** diphenylboric acid, **(Q)** disodium fluorophosphate, **(R)** doxycycline, **(S)** emodin, **(T)** enoxolone, **(U)** fenbendazole, **(V)** fendiline hydrochloride, **(W)** frequentine, **(X)** fendiline, **(Y)** ferron, **(Z)** gentian violet, **(AA)** hydroxyurea, **(AB)** ketoprofen, **(AC)** medroxyprogesterone acetate, **(AD)** maleic acid, **(AE)** myristic acid, **(AF)** NADIDE, **(AG)** nabumetone, **(AH)** onion oil, **(AI)** pergolide mesylate, **(AJ)** pimethixene maleate, **(AK)** piperacillin sodium, **(AL)** pomiferin, **(AM)** peanut oil, **(AN)** polysorbate 65, **(AO)** prochlorperazine dimaleate, **(AP)** reserpine, **(AQ)** riboflavin tetrabutyrate, **(AR)** ricobendazole, **(AS)** strophanthin K, **(AT)** sulconazole, **(AU)** triptonide, **(AV)** thiethylperazine malate, **(AW)** thonzonium bromide, **(AX)** verteporfin.

**Supplemental Figure 2. Validation of B-lp levels following treatment of 30 hits identified from a high-throughput drug screen of B-lp modulators.** Of the 49 identified B-lp-reducing compounds, we subjected 30 to further validation studies. We examined the effects of each compound with an 8-fold serial dilution from 8 µM to 0.0625 µM. Open circles represent each sample. Sample size (n) is listed below each treatment. Results were analyzed using one-way ANOVA to determine if the means of any treatment were significantly different from each other. When the one-way ANOVA was significant (p ≤ 0.05), a Dunnett’s test was conducted to determine which treatments differed significantly from vehicle treatment. Each p-value was adjusted for multiple comparisons with a Bonferroni correction and is listed on each graph. The compounds examined are **(A)** 3-methylcholanthrene, **(B)** acetaminophen, **(C)** cetrimonium bromide, **(D)** cyproterone, **(E)** calcipotriene, **(F)** cytidine 5’-monophosphate, **(G)** danazol, **(H)** danthron, **(I)** demecarium bromide, **(J)** doxycycline, **(K)** emodin, **(L)** enoxolone, **(M)** fenbendazole, **(N)** fendiline, **(O)** ferron, **(P)** hydroxyurea, **(Q)** ketoprofen, **(R)** medroxyprogesterone acetate, **(S)** maleic acid, **(T)** NADIDE, **(U)** nabumetone, **(V)** pergolide mesylate, **(W)** pimethixene maleate, **(X)** piperacillin sodium, **(Y)** pomiferin, **(Z)** prochlorperazine dimaleate, **(AA)** reserpine, **(AB)** riboflavin tetrabutyrate, **(AC)** ricobendazole, **(AD)** strophanthin K, **(AE)** sulconazole, **(AF)** triptonide, **(AG)** thiethylperazine malate, **(AH)** thonzonium bromide, **(AI)** verteporfin.

**Supplemental Figure 3. High-dose enoxolone-treated animals are shorter than vehicle-treated animals.** Boxplots of standard-length measurements of 5 dpf animals treated for 48 hours with vehicle, 5 µM lomitapide, and 8 µM or 4 µM enoxolone. After treatment, animals were imaged, and standard lengths were measured from the images. Animals treated with 5 µM lomitapide or 8 µM enoxolone were significantly shorter than vehicle-treated animals (one-way ANOVA, *F*(3,111) = 12.68, *p* = 3.5×10^−7^; Dunnett’s test 5 µM lomitapide n = 24 versus vehicle n = 32, *p* = 4.9×10^−2^; Dunnett’s test 8 µM enoxolone n = 27 versus vehicle n = 32, *p* = 9.1×10^−6^). Animals treated with 4 µM enoxolone lengths were unchanged compared to vehicle treatment (Dunnett’s test 4 µM enoxolone n = 32 versus vehicle n = 32, *p* = 0.095).

**Supplemental Figure 4. Summary analysis of differentially expressed genes following enoxolone treatment. (A)** Principal component analysis (PCA) of RNAseq samples for treatment and the duration of treatment. **(B)** Boxplot summarizing the expression pattern of *insig1* following the vehicle and 8 µM enoxolone treatment. I*nsig1* is highly expressed in enoxolone-treated animals at every duration measured. Each data point is the transcripts per million (TPM) of an individual sample. **(C)** Gene ontology (GO) analysis for the biological process of differentially expressed genes at 4 hours post-treatment (hpt). **(D)** GO analysis for biological process of differentially expressed genes at 8 hpt. **(E)** GO analysis for biological process of differentially expressed genes at 12 hpt. **(F)** GO analysis for biological process of differentially expressed genes at 16 hpt. **(G)** GO analysis for biological process of differentially expressed genes at 24 hpt.

**Supplemental Table 1. Raw data from JHDL primary screen.**

**Supplemental Table 2. Analyzed summary data from JHDL primary screen.**

**Supplemental Table 3. List of hits from primary screen with results of secondary screening validation studies.**

**Supplemental Table 4. Full RNAseq results at all timepoints following enoxolone treatment.**

**Supplemental Table 5. Differentially expressed genes at each time post enoxolone treatment.**

**Supplemental Table 6. Gene ontology at each time post enoxolone treatment.**

**Supplemental Table 7. Overlap of enoxolone-responsive differentially expressed genes with HNF4α and HNF4γ knockout datasets.**

