## Supplementary figures and images for "A whole-animal phenotypic drug screen identifies suppressors of atherogenic lipoproteins"

### 3-Aminobenzyl (___) Incomplete name_JHDL_17_015_055_2019_09_13_12_43.pdf

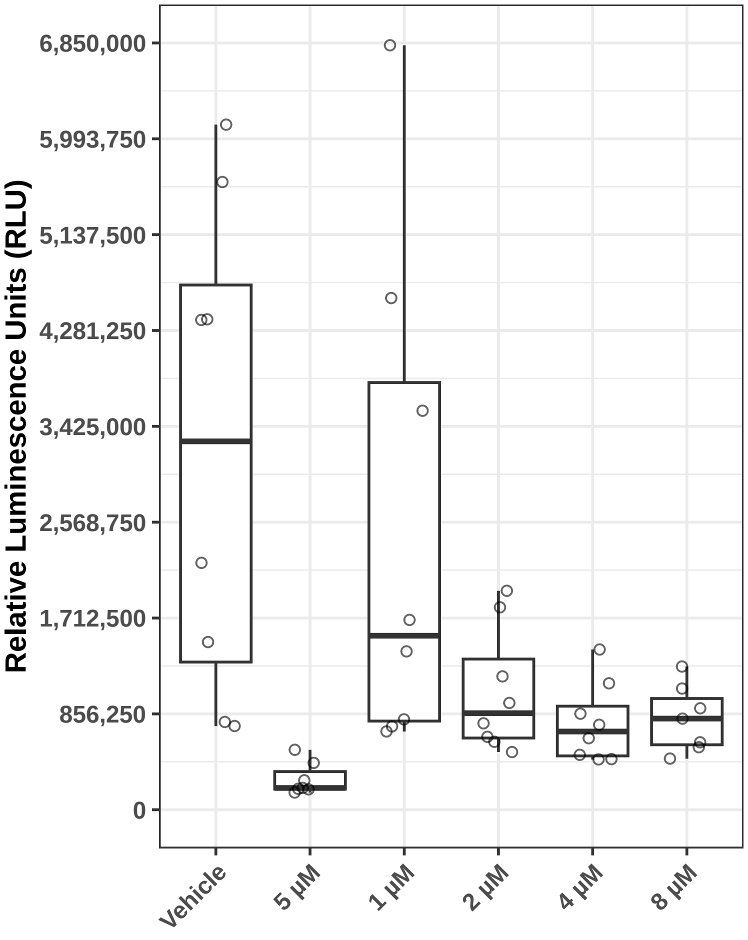

### 0000016_01_TH.jpg

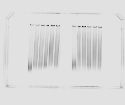

### 0000017_01_TH.jpg

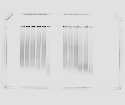

### 0000019_01_TH.jpg

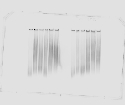

### 0000020_01_TH.jpg

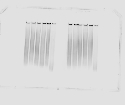

### 0000021_01_TH.jpg

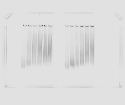

### 0000022_01_TH.jpg

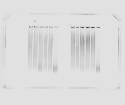

### 0000023_01_TH.jpg

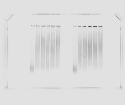

### 0000024_01_TH.jpg

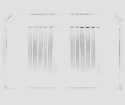

### 0000025_01_TH.jpg

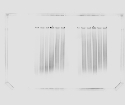

### 0000026_01_TH.jpg

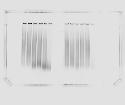

### 0000027_01_TH.jpg

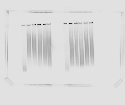

### 0000028_01_TH.jpg

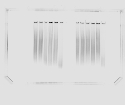

### 0000029_01_TH.jpg

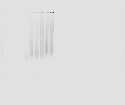

### 0000030_01_TH.jpg

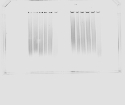

### 0000031_01_TH.jpg

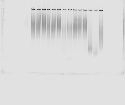

### 0000032_01_TH.jpg

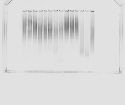

### CETRIMONIUM BROMIDE_JHDL_9_036_036_2019_06_28_11_39.pdf

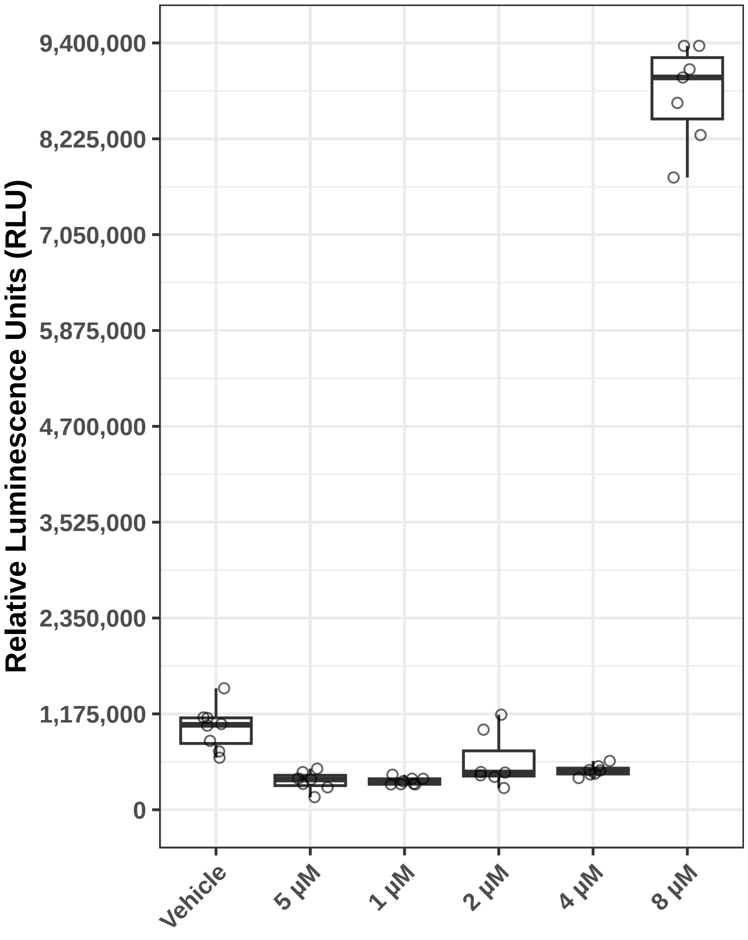

### Cinnamon oil (Cinnamon leaf oil)_JHDL_29_027_057_2019_12_06_13_27.pdf

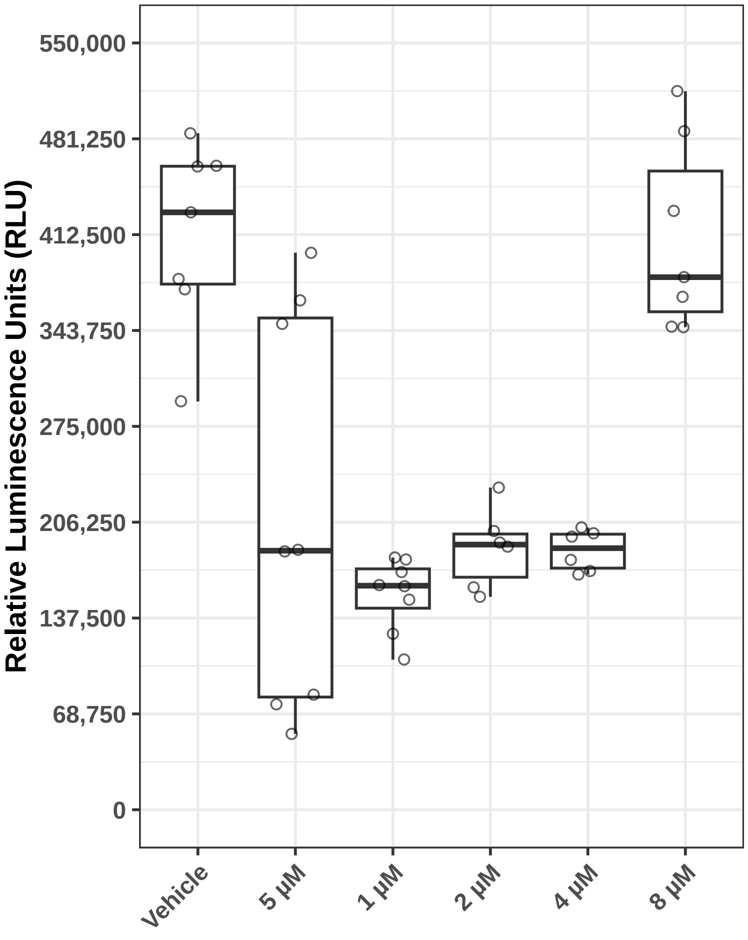

### CYPROTERONE_JHDL_8_034_024_2019_06_21_11_38.pdf

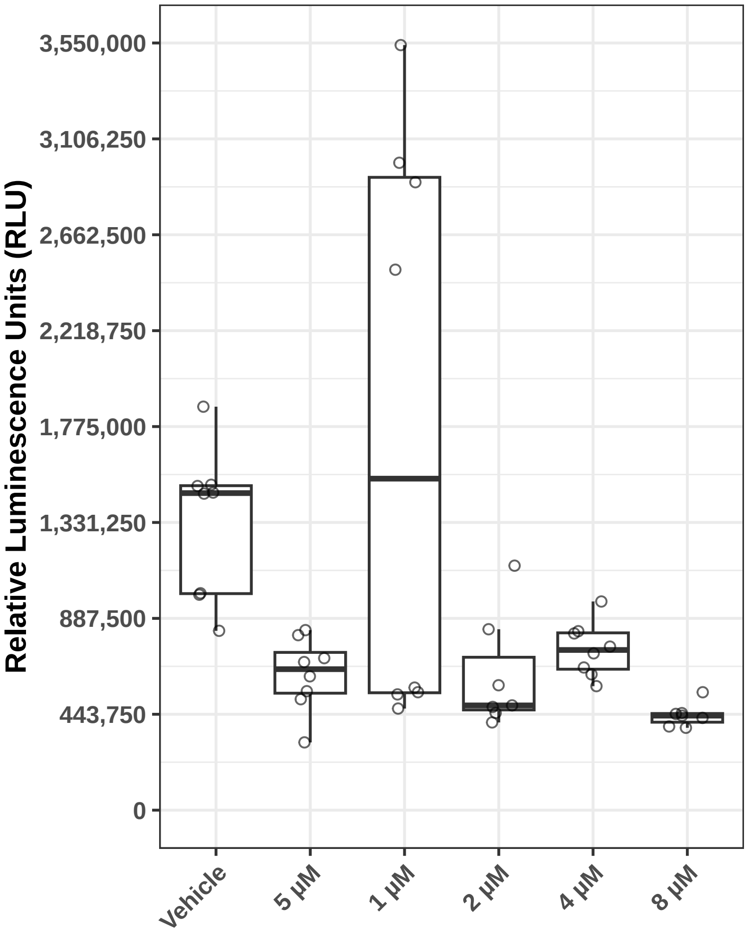

### Cytidine 5'-diphosphate┬átrisodium┬ásalt_JHDL_27_005_025_2019_11_08_11_10.pdf

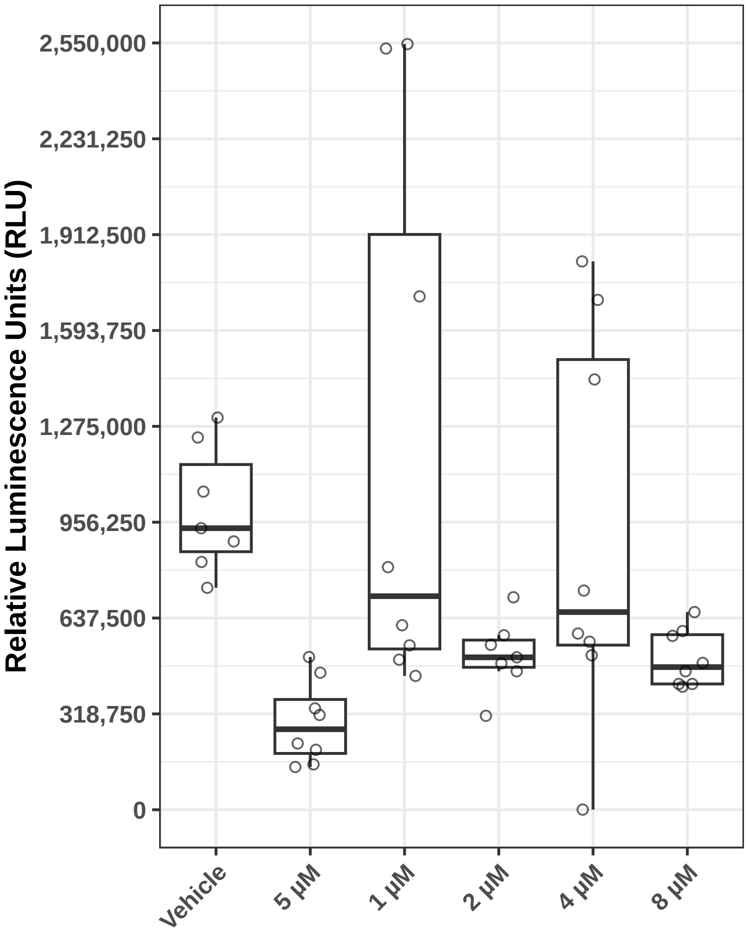

### Cytidine-5'-diphosphate trisodium salt_JHDL_27_005_025_2019_11_08_11_10.pdf

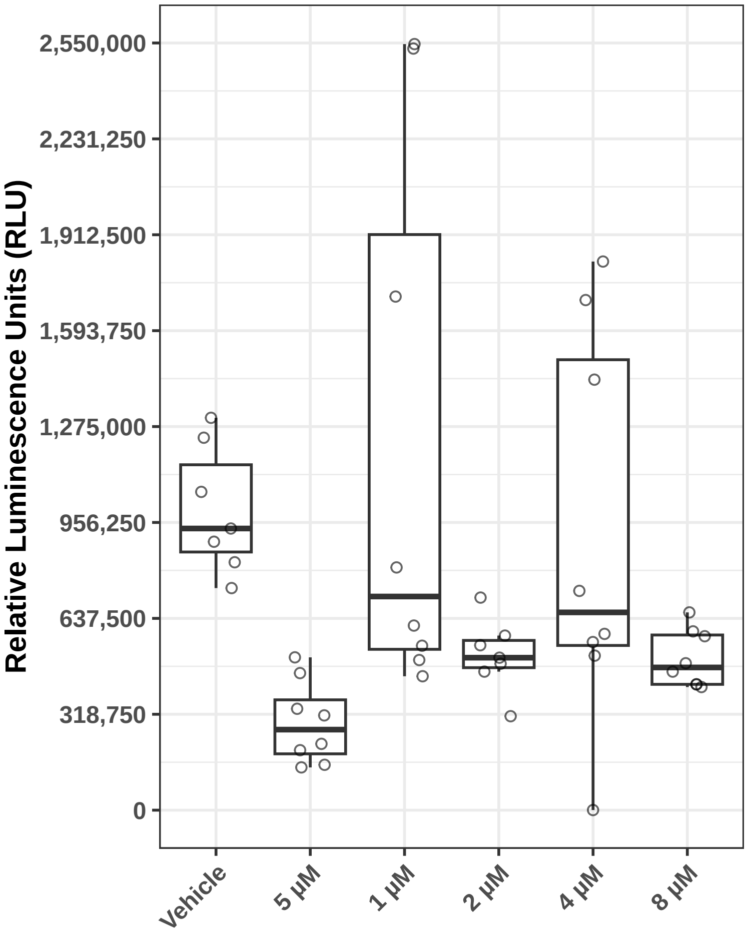

### CYTOCHALASIN N_JHDL_5_017_017_2019_05_17_11_22.pdf

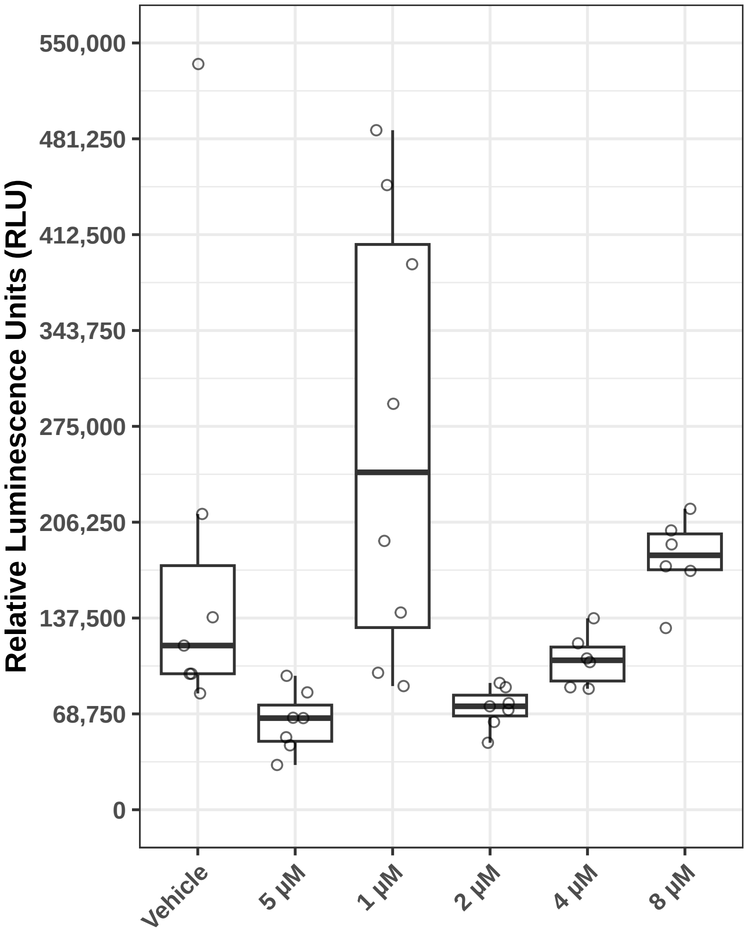

### Danazol_JHDL_16_008_008_2019_09_13_11_01.pdf

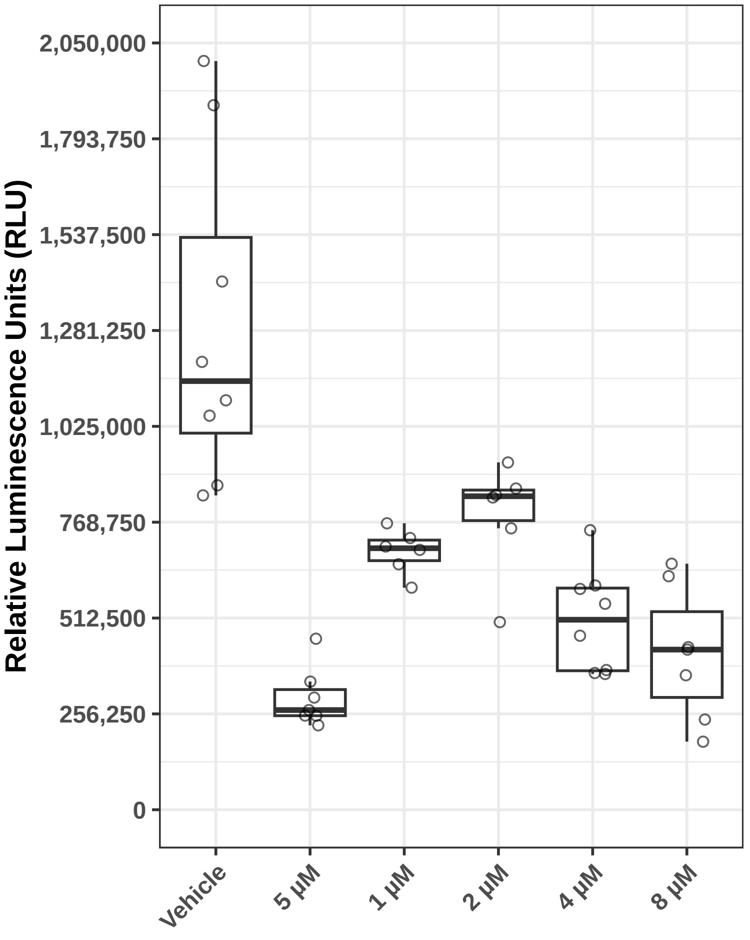

### Danazol_JHDL_35_026_071_2020_01_31_12_45.pdf

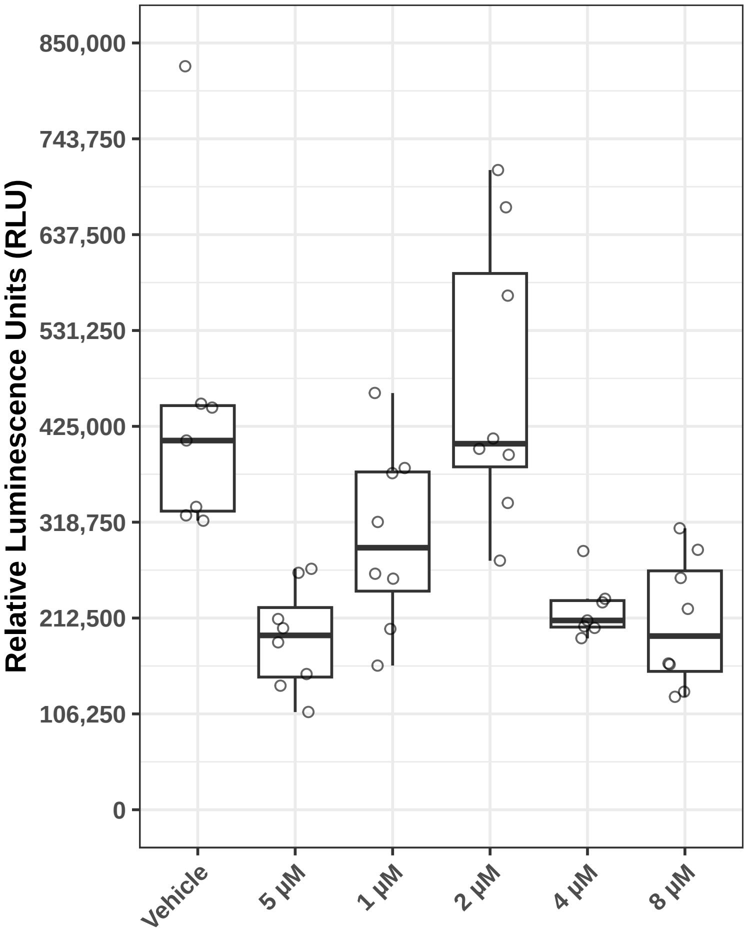

### Demecarium bromide_JHDL_35_024_069_2020_01_31_12_41.pdf

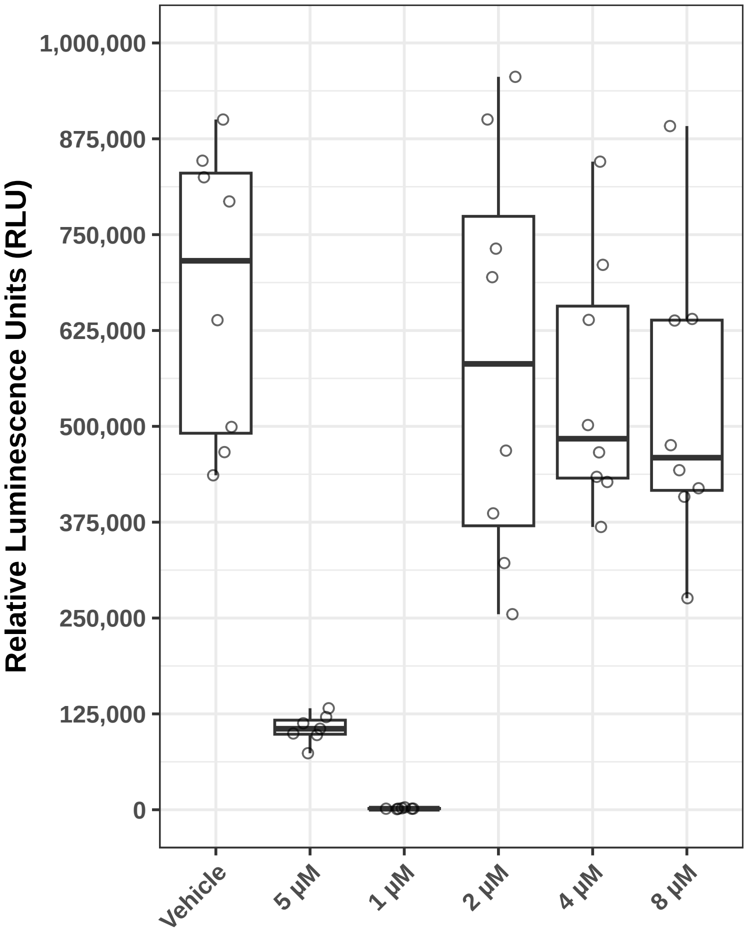

### DOX_0000027_01_600.png

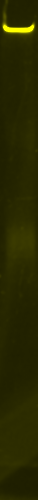

### DOX_0000027_01_Chemi.png

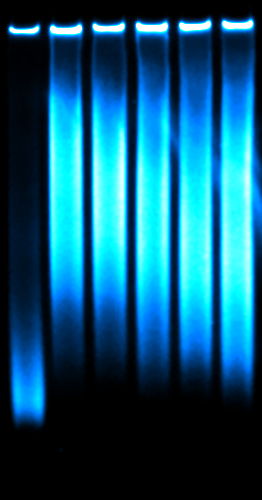

### Fendiline_JHDL_28_030_020_2019_12_06_11_34.pdf

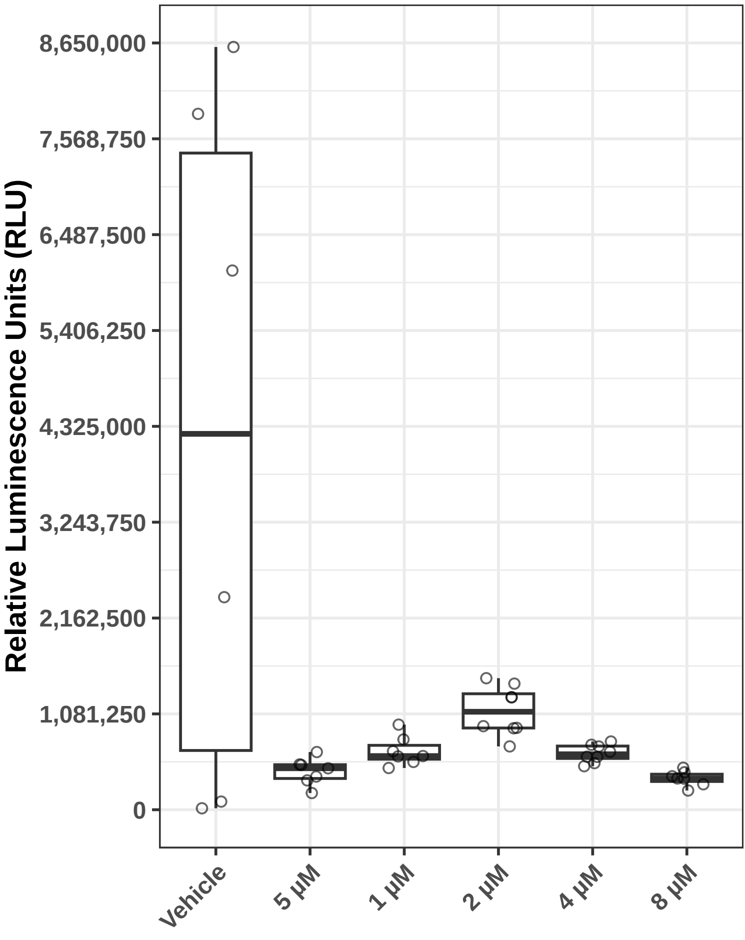

### Ferron (8-Hydroxy-7-iodo-5-quinolinesulfonic acid)_JHDL_29_009_039_2019_12_06_12_17.pdf

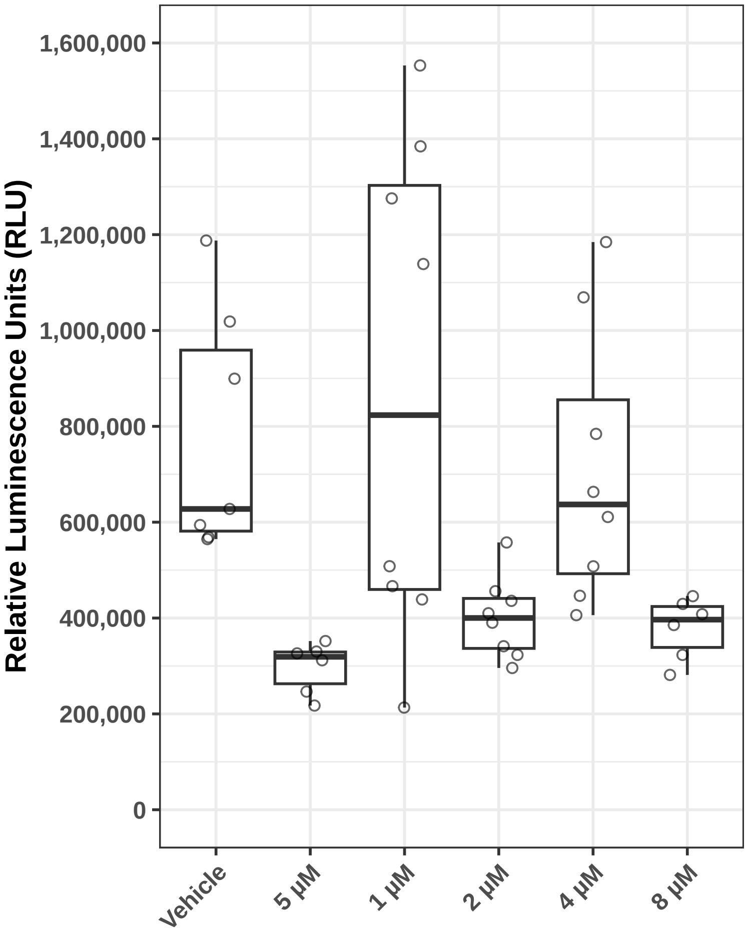

### Ferron_JHDL_15_032_047_2019_08_30_12_13.pdf

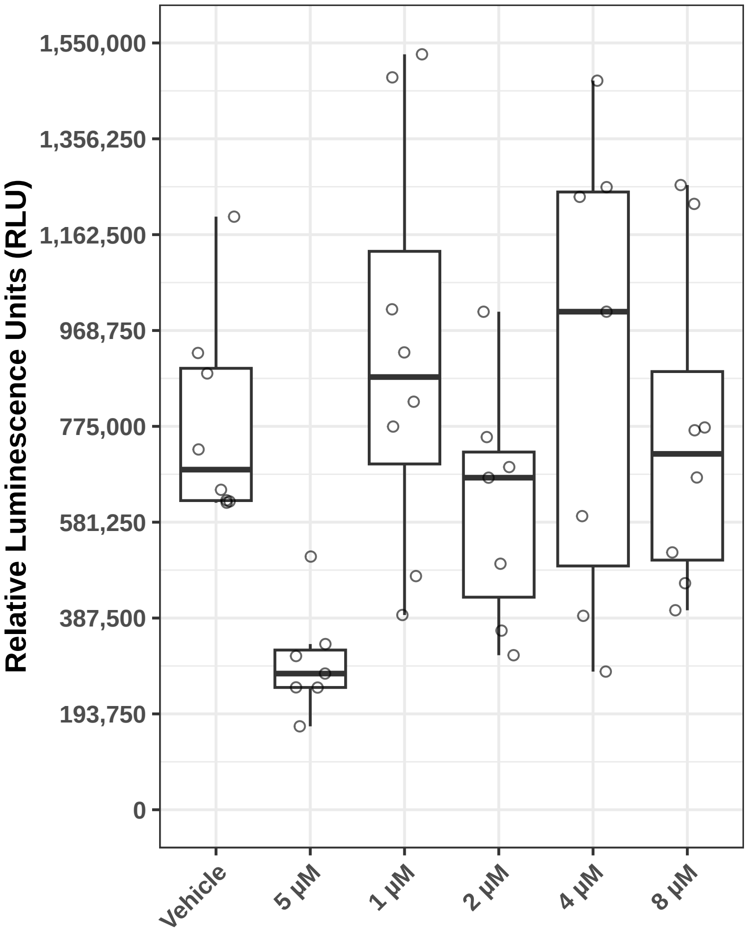

### Ferron_JHDL_29_009_039_2019_12_06_12_17.pdf

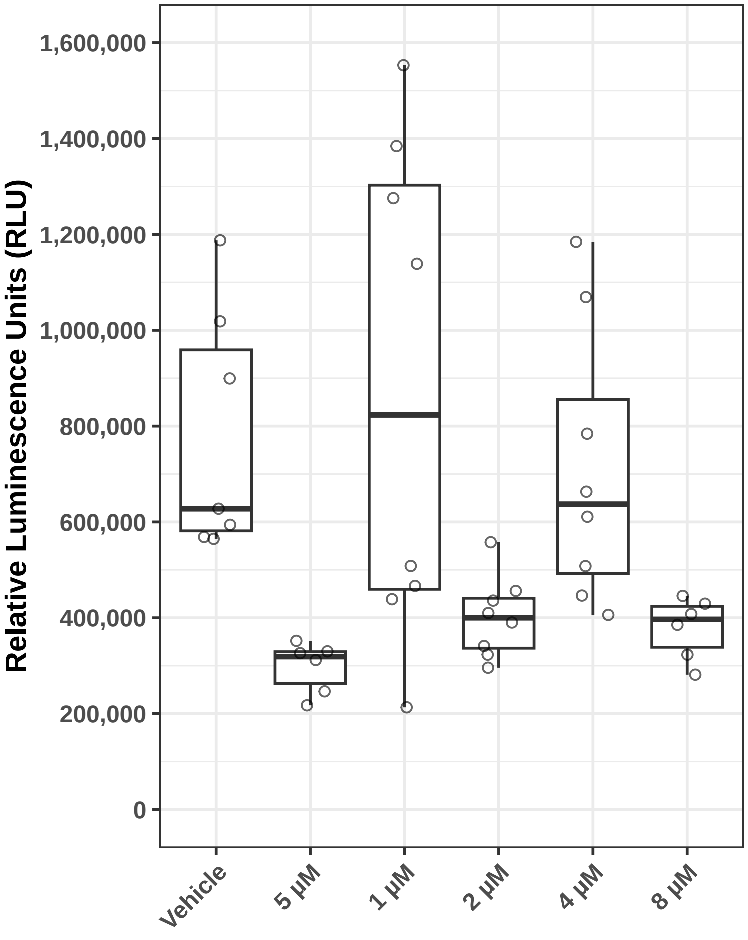

### FREQUENTIN_JHDL_3_040_007_2019_04_26_14_14.pdf

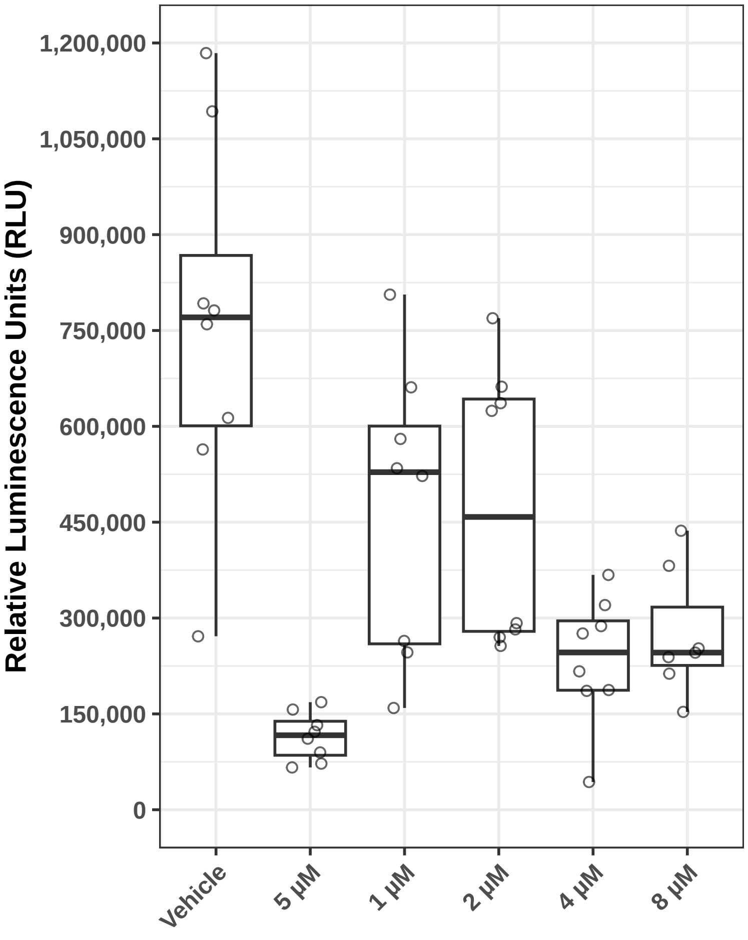

### Frequentine_JHDL_3_040_007_2019_04_26_14_14.pdf

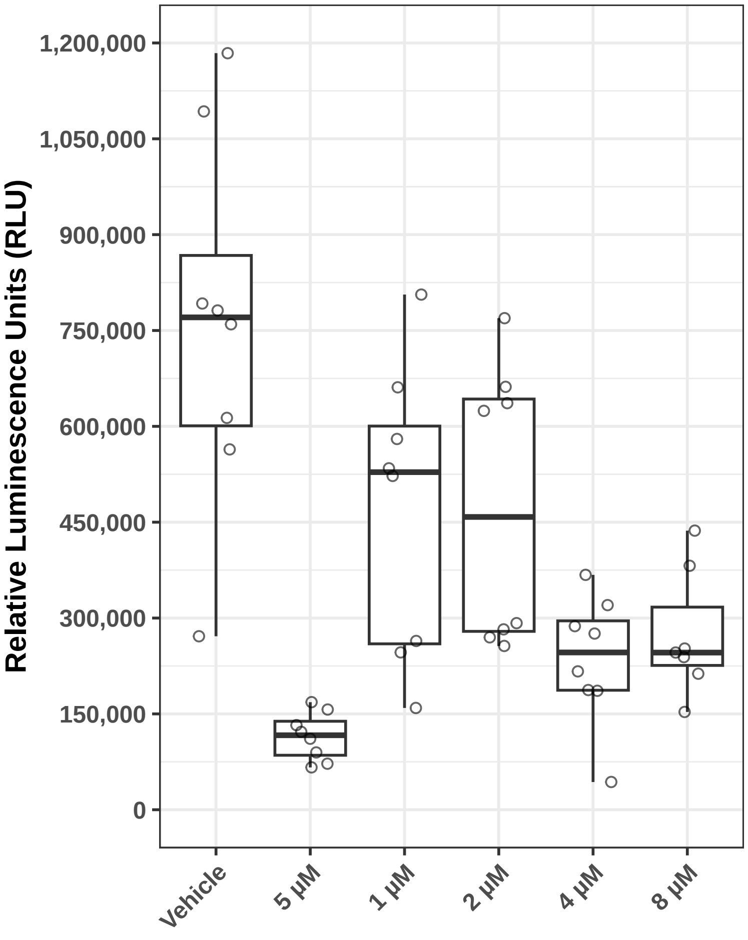

### GENTIAN VIOLET_JHDL_3_017_018_2019_04_26_12_48.pdf

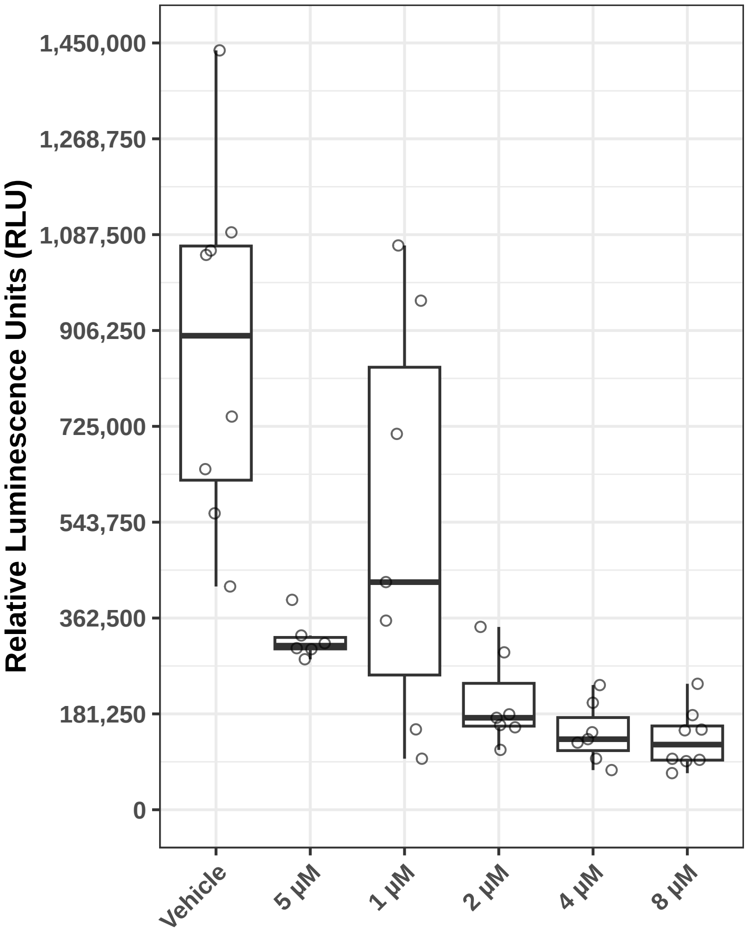

### Maleic acid_JHDL_17_019_004_2019_09_20_09_52.pdf

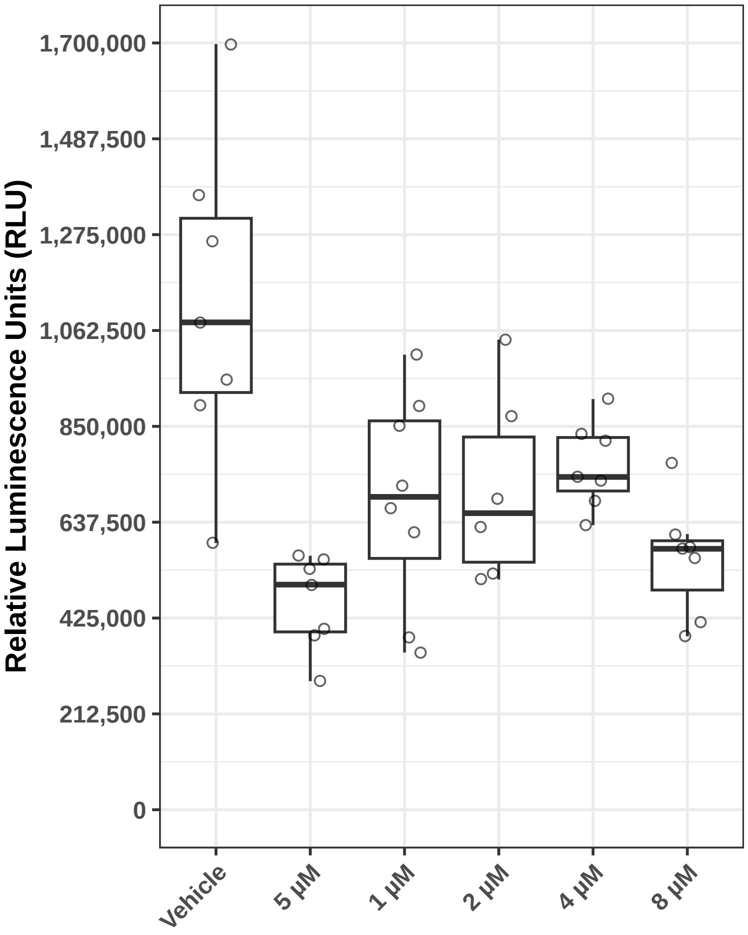

### MEDROXYPROGESTERONE ACETATE_JHDL_4_008_008_2019_05_10_11_14.pdf

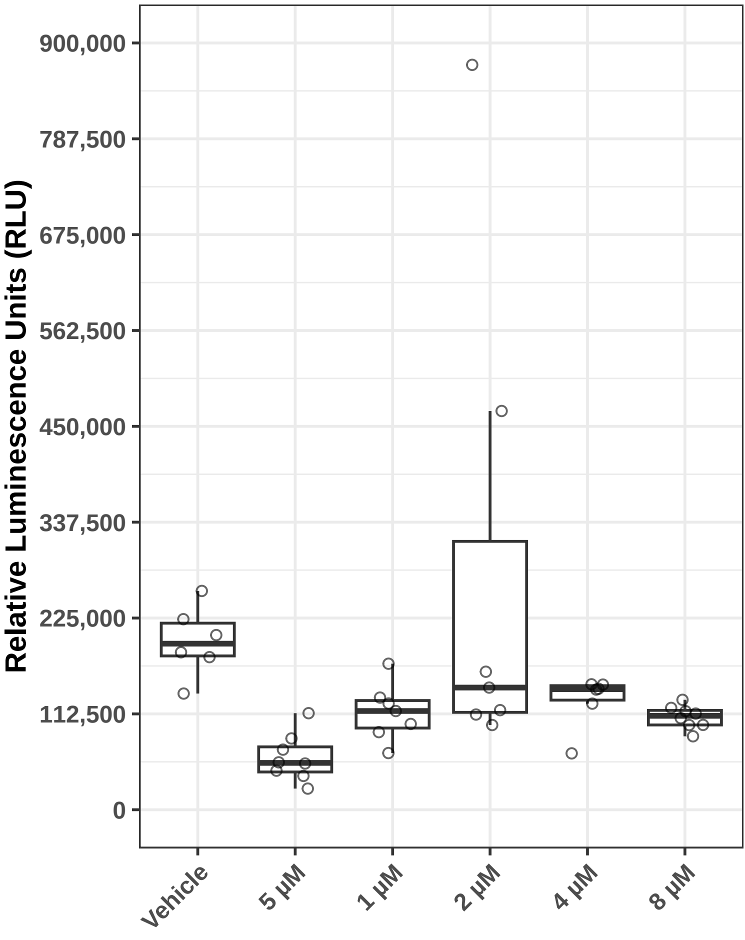

### Medroxyprogesterone acetate_JHDL_14_027_002_2019_08_30_10_37.pdf

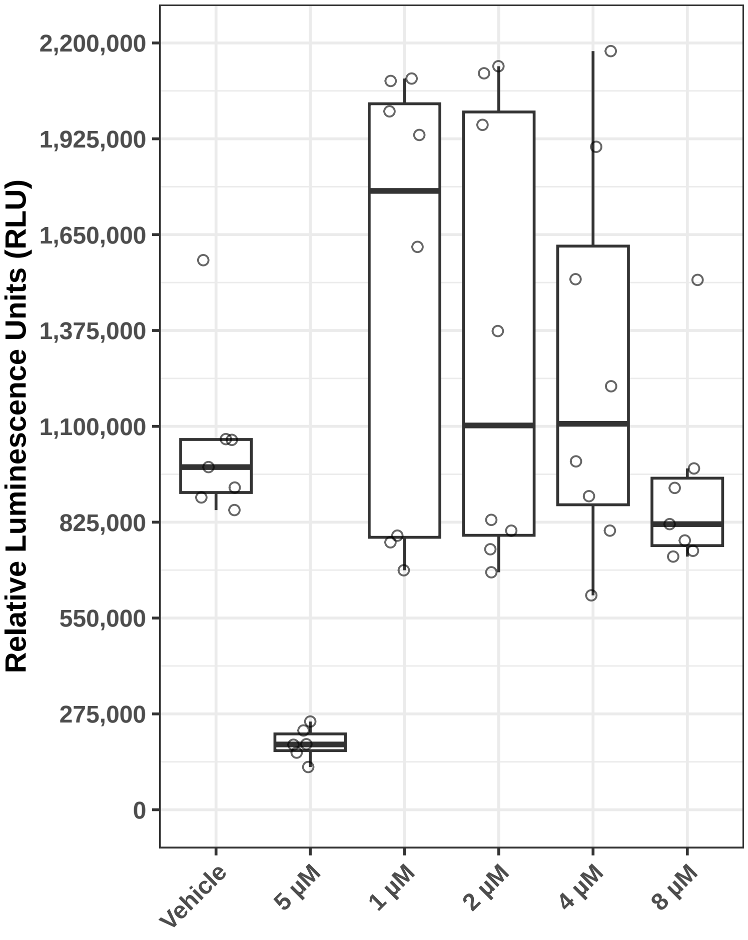

### PAGE Organization Document.docx

12/12/2025

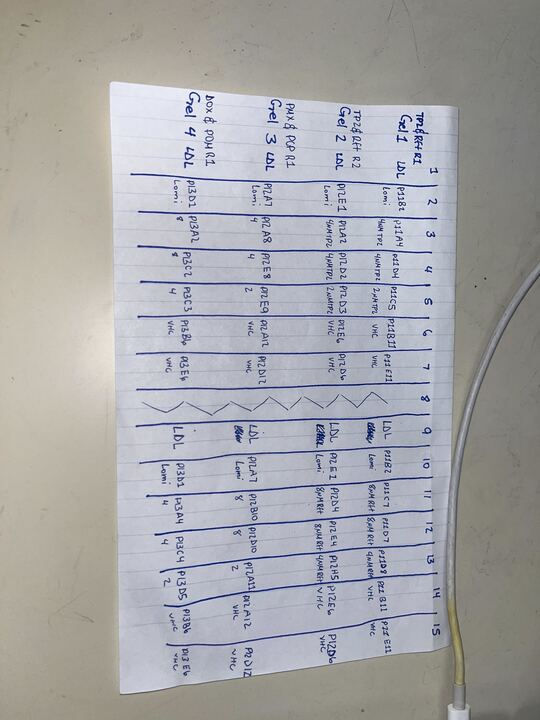

12/16/2025

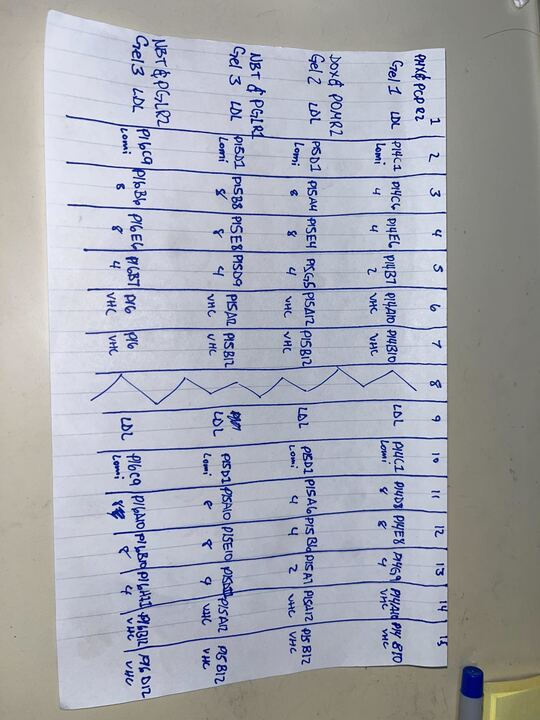

12/17/2025
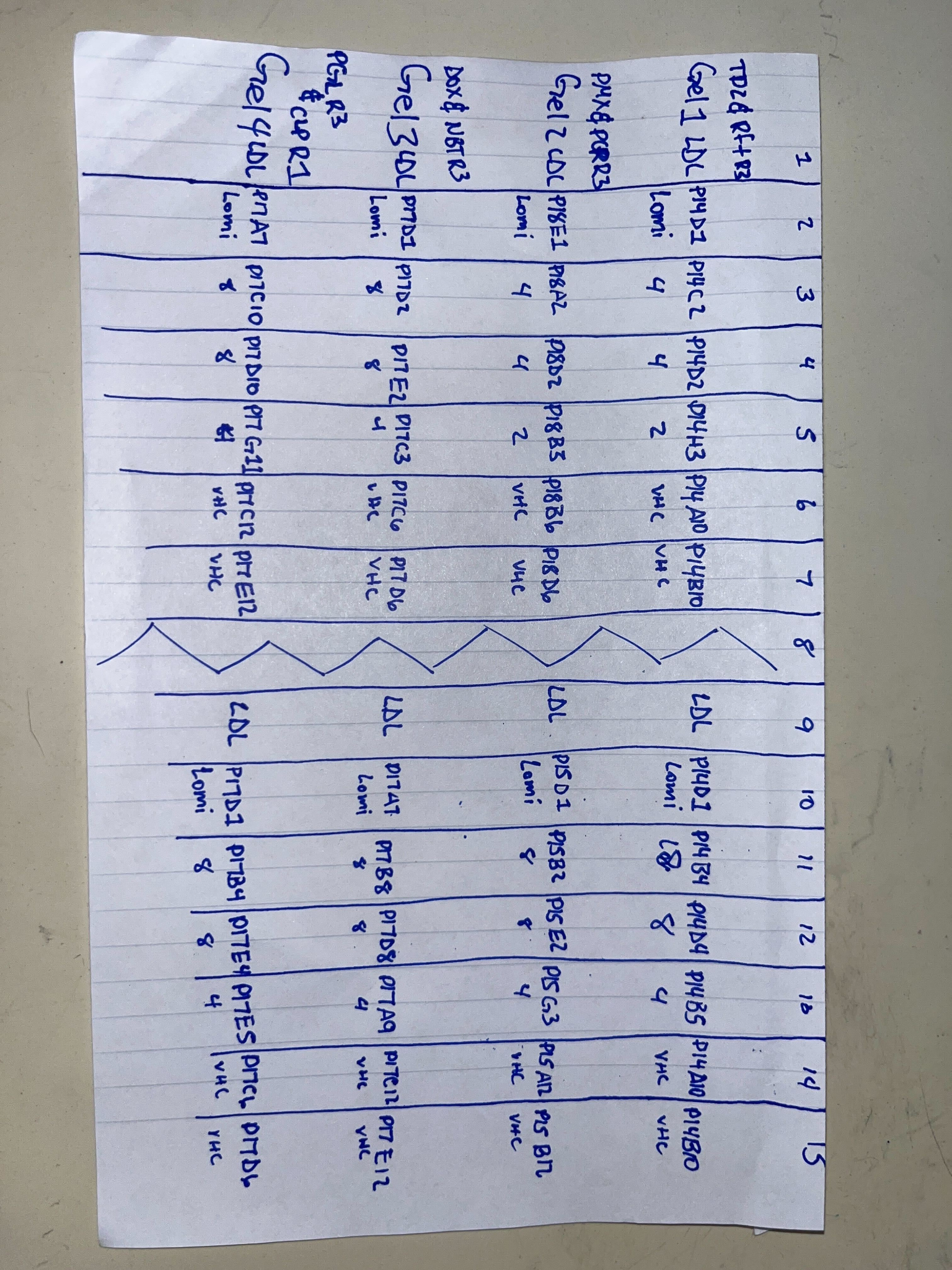

01/21/2026

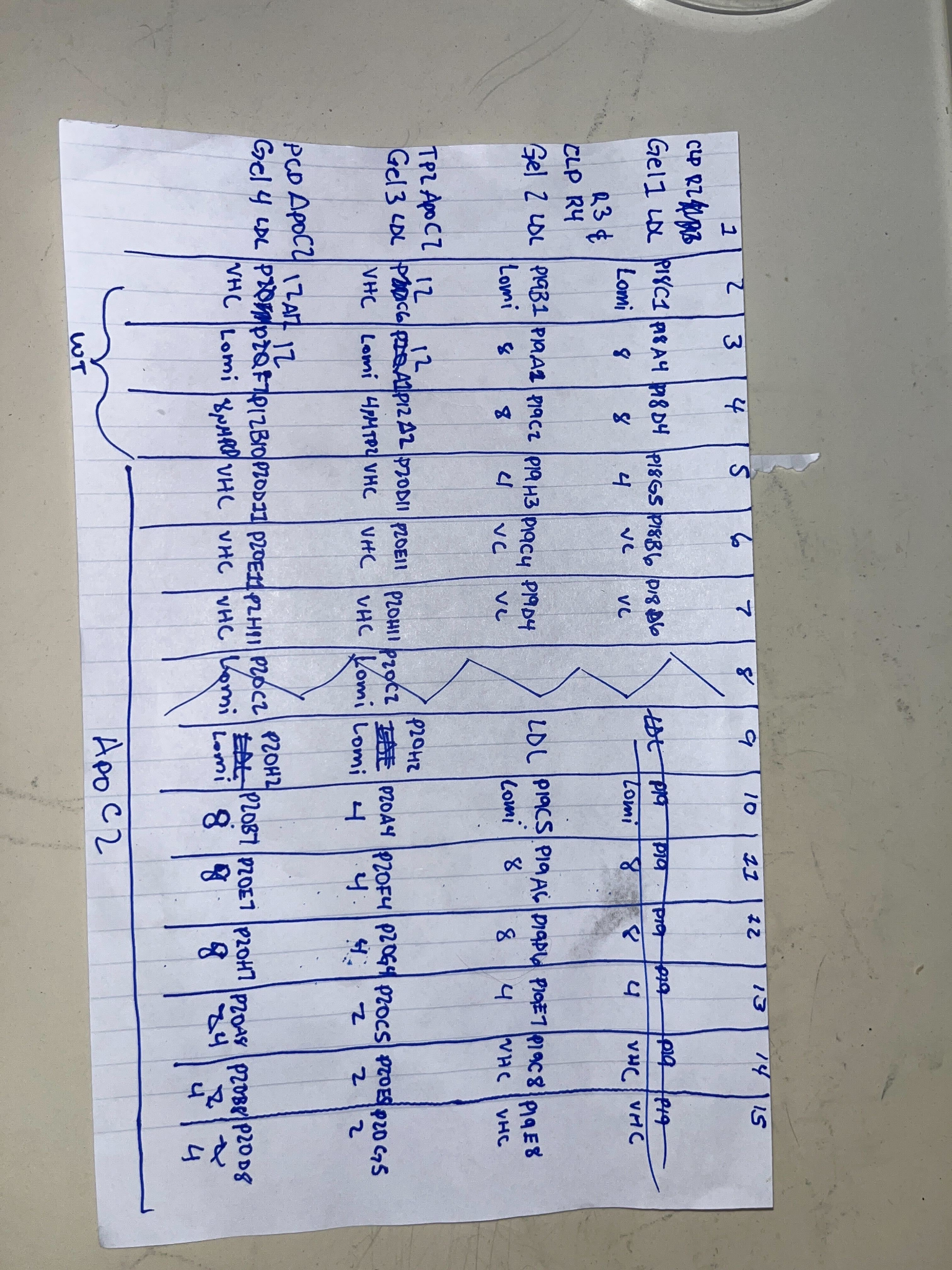

### PCP_0000026_01_600.png

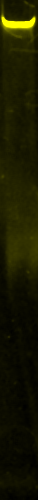

### PCP_0000026_01_Chemi.png

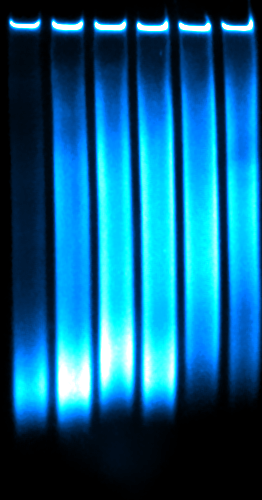

### Peanut oil_JHDL_30_019_019_2019_12_13_11_22.pdf

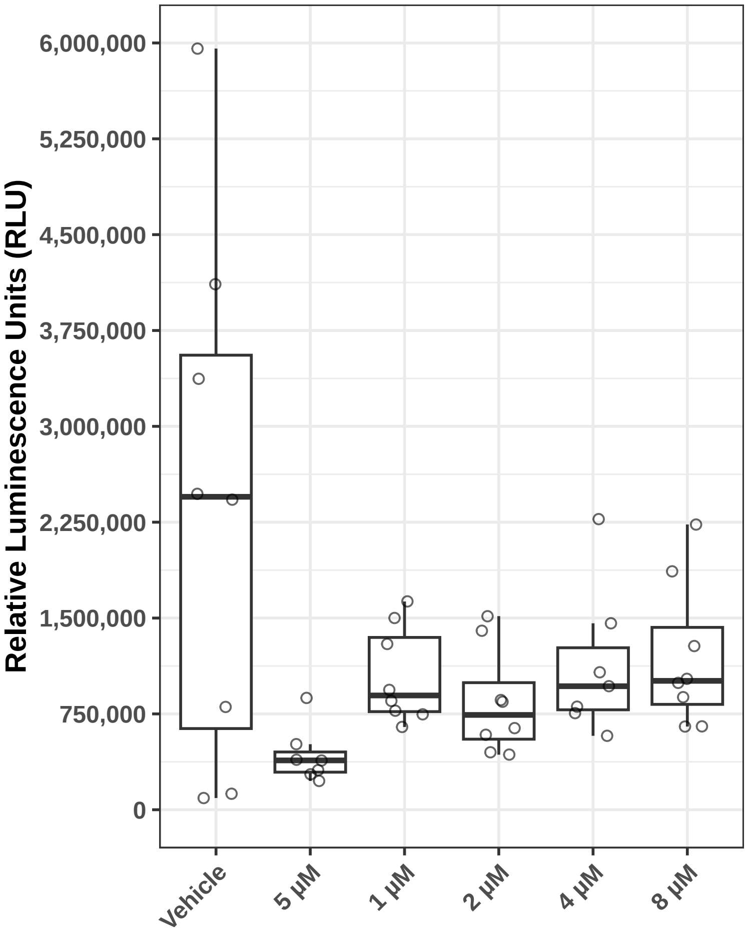

### Polysorbate 65 (Tween 65)_JHDL_32_035_020_2020_01_24_10_41.pdf

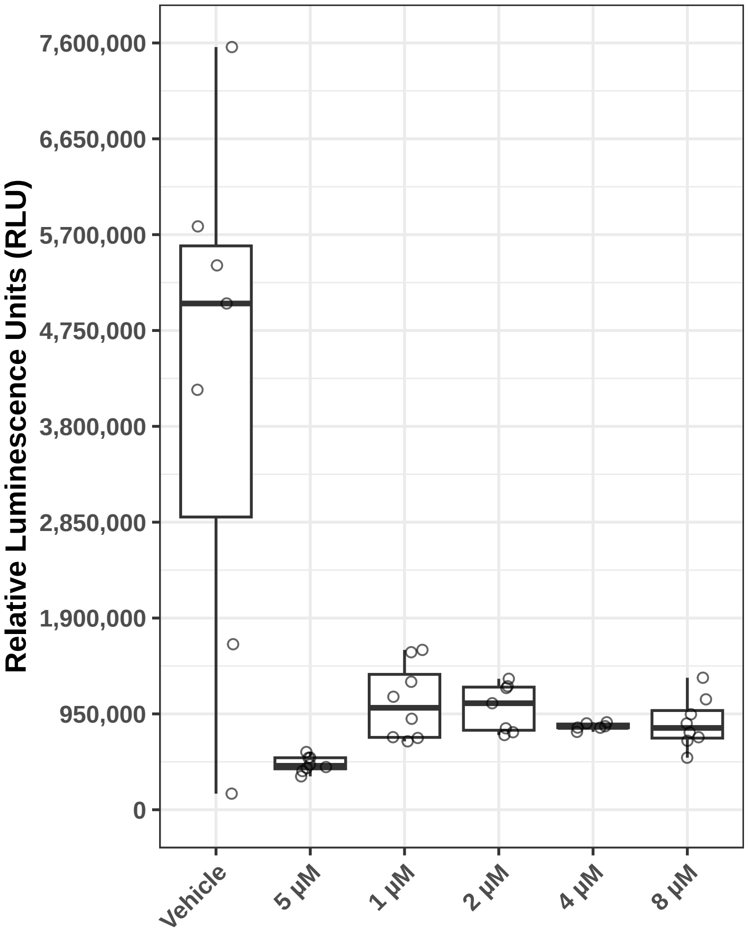

### Riboflavin tetrabutyrate_JHDL_35_032_077_2020_01_31_12_55.pdf

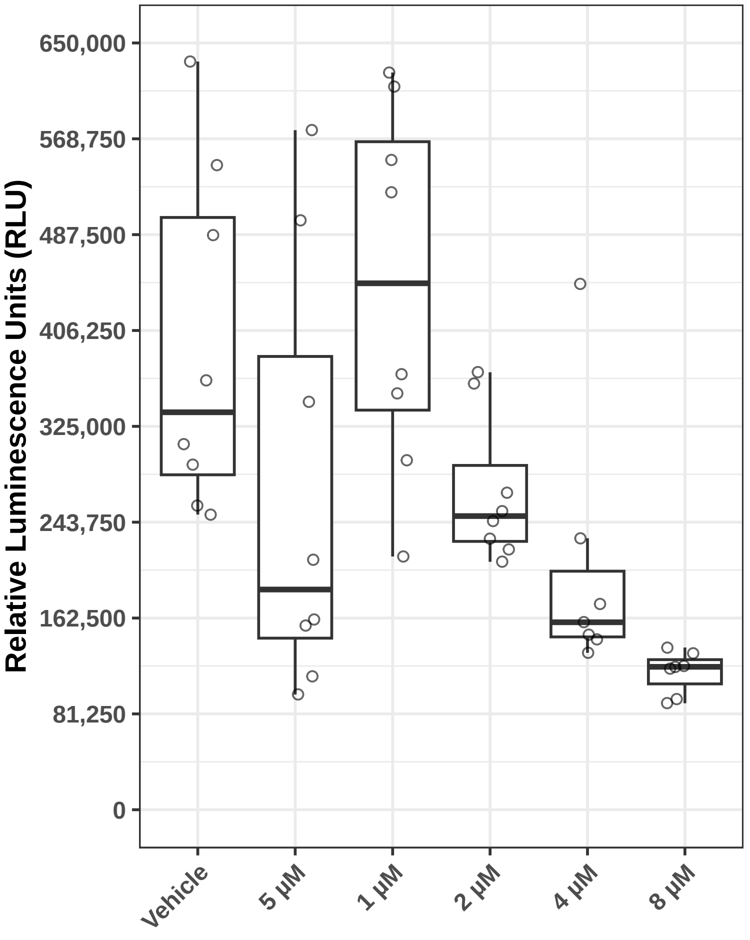

### Ricobendazole (Albendazole oxide)_JHDL_32_035_020_2020_01_24_10_41.pdf

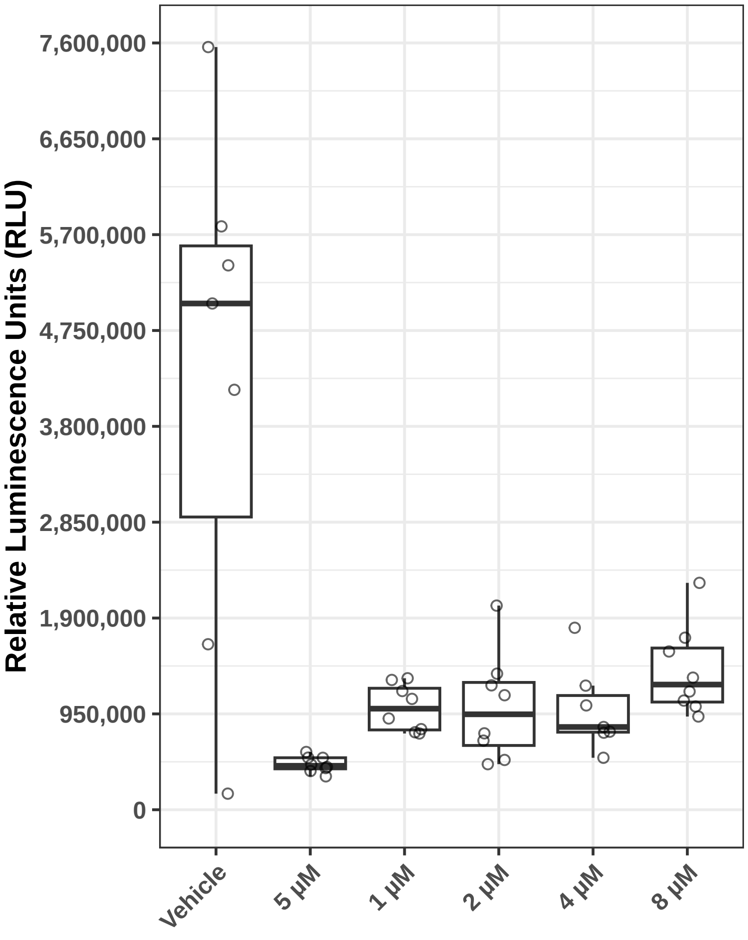

### Ricobendazole_JHDL_32_035_020_2020_01_24_10_41.pdf

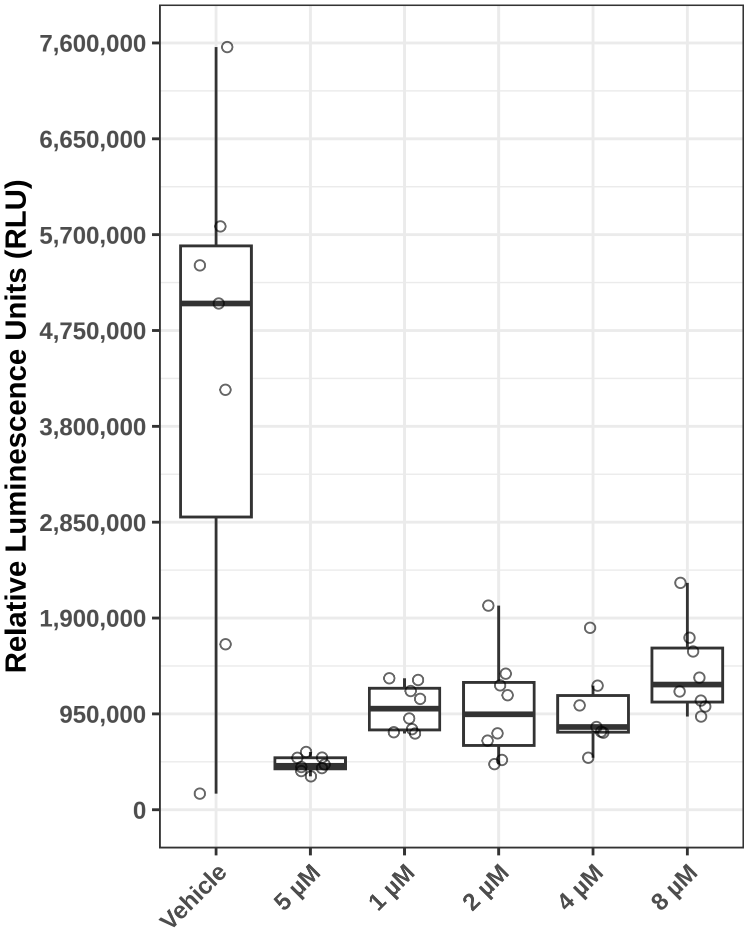

### Supplemental Figure 1

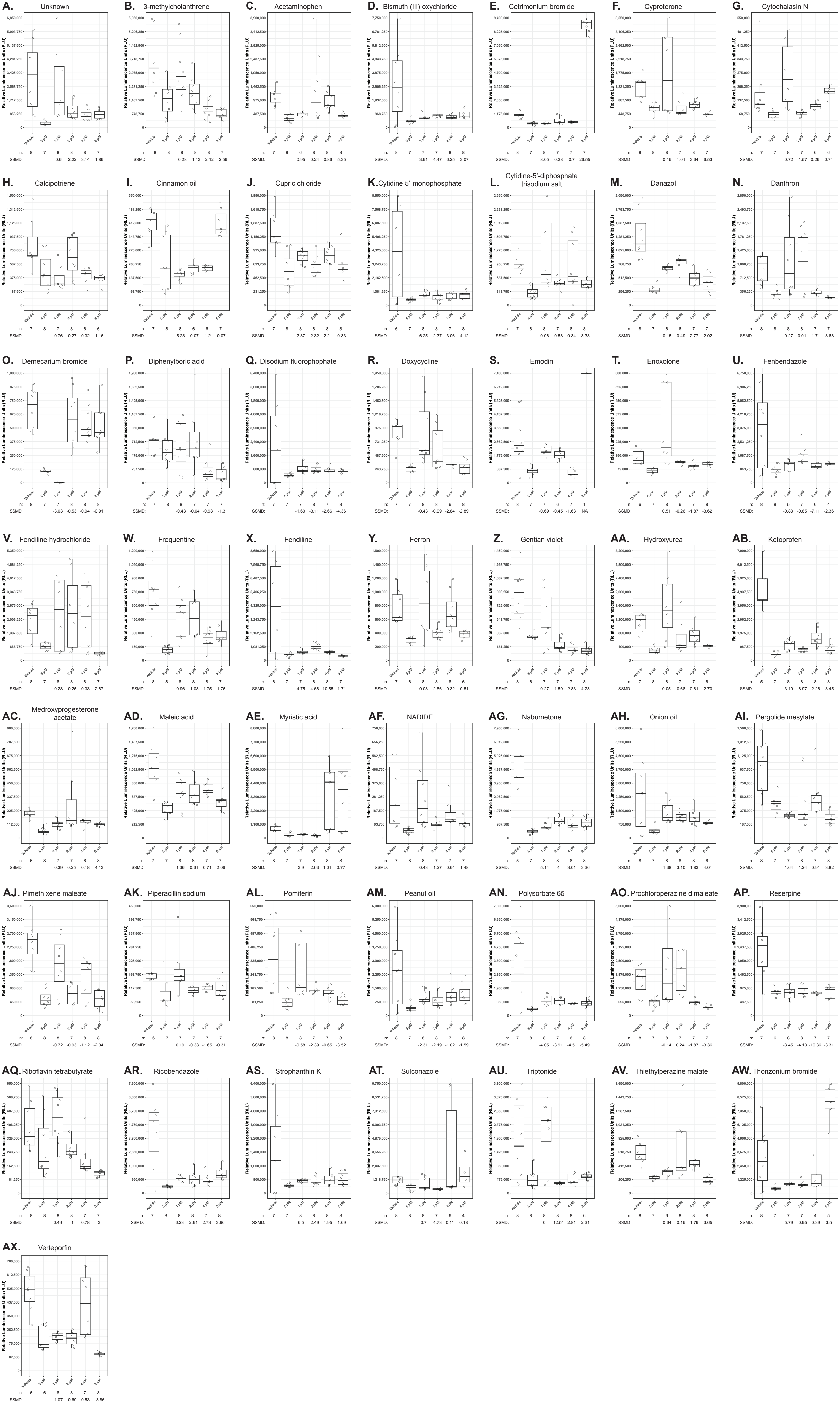

### Supplemental Figure 2

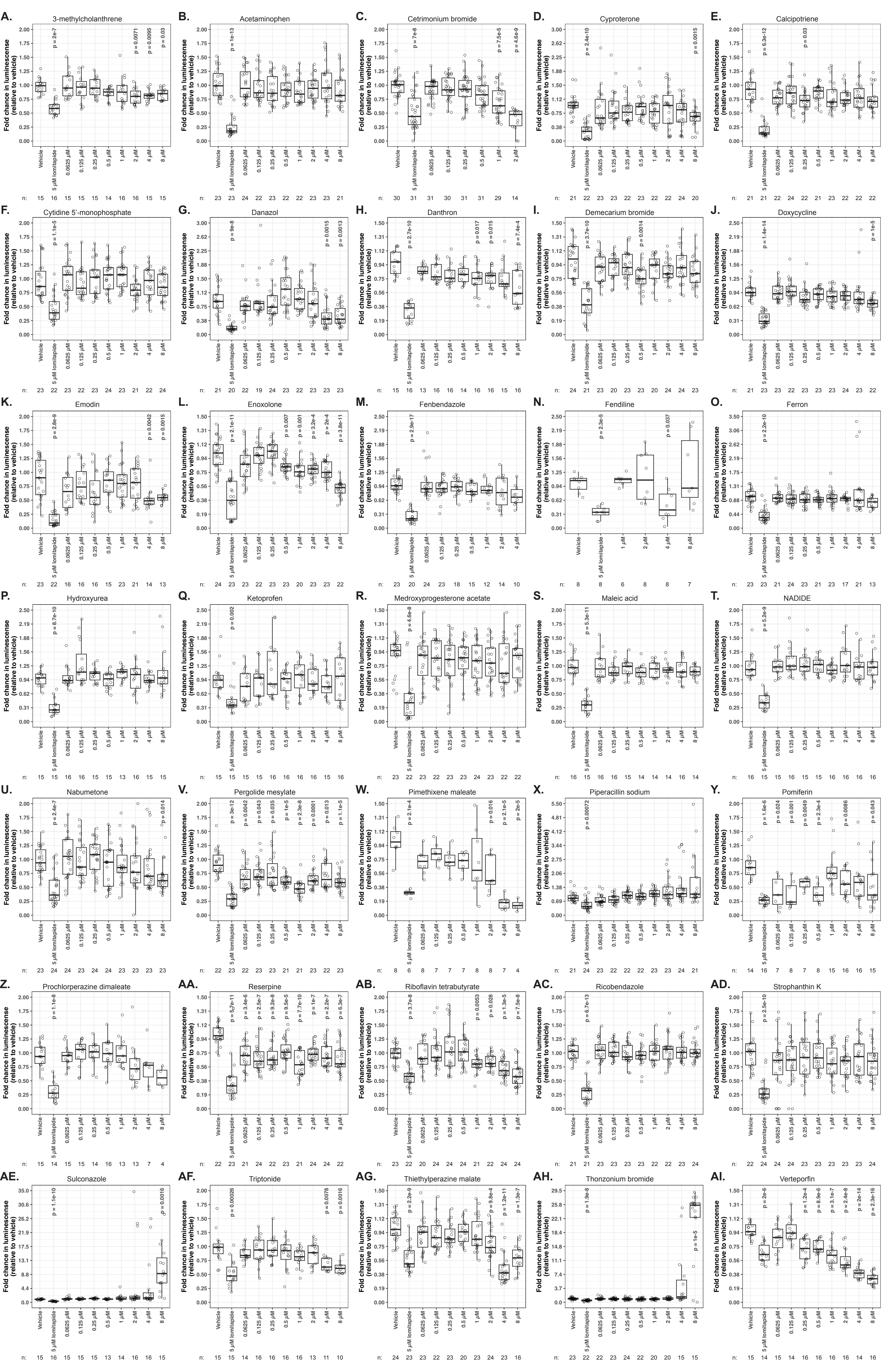

### Supplemental Figure 3

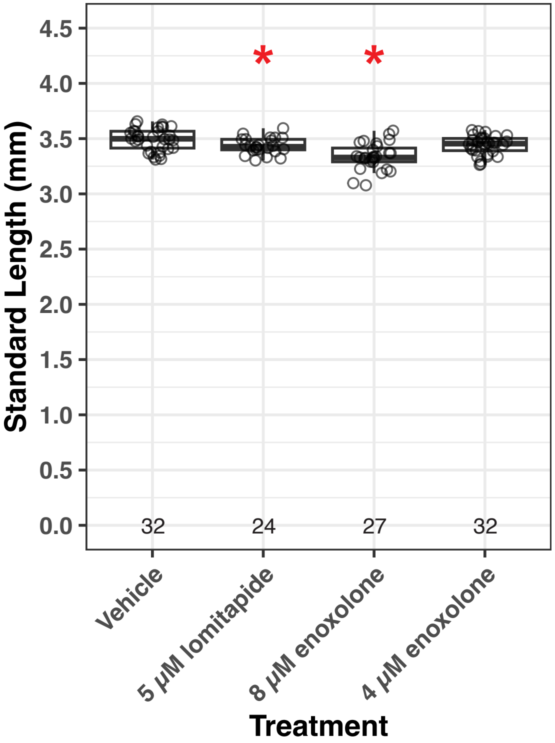

### Supplemental Figure 4

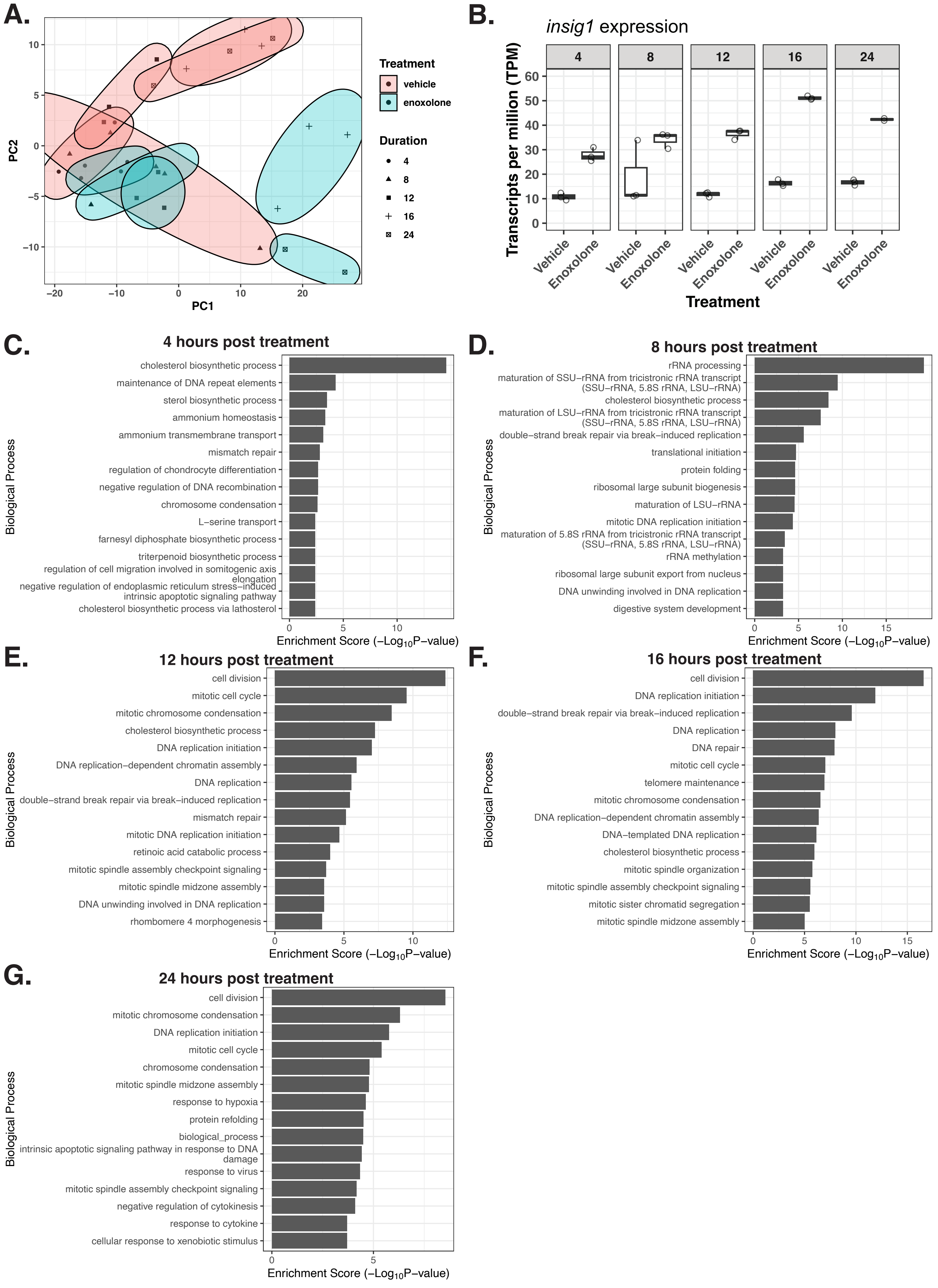

### Thiethylperazine malate_JHDL_34_025_030_2020_01_31_11_18.pdf

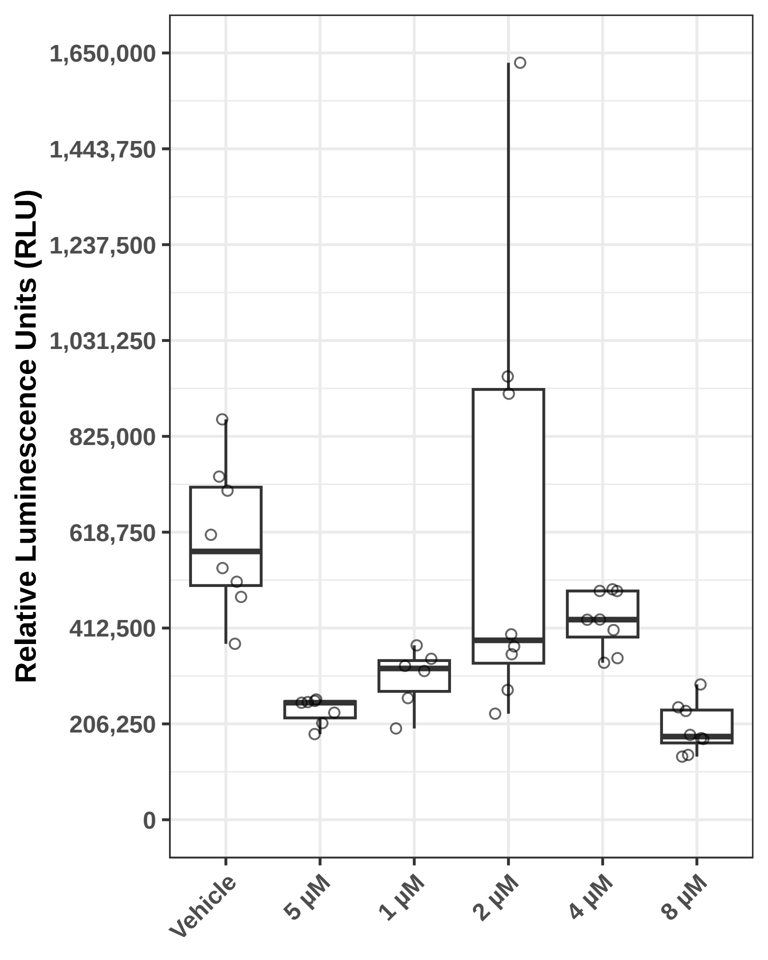

### TPZ_0000025_01_600.png

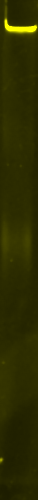

### TPZ_0000025_01_Chemi.png

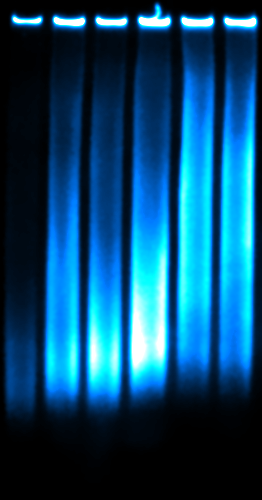
