## Supplementary material for "A whole-animal phenotypic drug screen identifies suppressors of atherogenic lipoproteins": Figure_making.qmd

````
---
title: "Figure making for the primary JHDL screen results, hits, and enoxolone studies"
subtitle: ""
author: "Dan Kelpsch"
format:
  html:
    self-contained: true
editor: visual
---

```{r}
#| include: false
list.of.packages <- c("tidyverse", "ggpubr", "ggbeeswarm", "scales", "readxl", "plater", "car", "inflection", "gt", "ggrepel")
new.packages <- list.of.packages[!(list.of.packages %in% installed.packages()[,"Package"])]
if(length(new.packages)) install.packages(new.packages)
lapply(list.of.packages, library, character.only = TRUE)

##round_any() is a function in the package plyr, but plyr has some 
##compatibility issues with the tidyverse, thus just writing in the 
##function manually
round_any = function(x, accuracy, f=round){f(x/ accuracy) * accuracy}
```

# Figure 1

## JHDL Primary Screen

## Data import and cleaning

```{r}
JHDL_all_data <- as_tibble(read.csv("JHDL_all_data.csv",
                                    na.strings = c("#N/A")))
JHDL_summary_data <- as_tibble(read.csv("JHDL_summary_data.csv",
                                        na.strings = c("#N/A", "#NUM!")))

#Clean up data
JHDL_all_data <- JHDL_all_data %>% 
  filter(Column != c(1,12))
jhdl_all_data_compounds <- JHDL_all_data %>% pull(Compound_name) %>% unique()

JHDL_summary_data <- JHDL_summary_data %>% filter(Column != c(1,12)) 

jhdl_summary_data_compounds <- JHDL_summary_data %>% pull(Compound_name) %>% unique()

JHDL_summary_nocontrol <- JHDL_summary_data %>% 
  filter(Drug != 'Lomitapide' & Drug != 'Vehicle')
JHDL_summary_controls <- JHDL_summary_data %>% 
  filter(Drug == 'Lomitapide' | Drug == 'Vehicle')

dir.create("./Output_data", showWarnings = FALSE)
```

## Manhattan plot

```{r}
ggplot(data = JHDL_summary_nocontrol, 
       aes(x = X, 
           y = hsSSMD)) +
  theme_classic() + 
  geom_point(shape = 1) +
  geom_hline(yintercept=c(-5,-3,-2,-1.645,-1.25,-1,1,1.25,1.645,2,3,5), 
             linetype="dashed") +
  scale_x_discrete(expand = c(0,0)) +
  scale_y_continuous(
    name = "SSMD",
    labels = comma, limits = c(-14,30), #c(lower limit, upper limit)
    breaks = seq(-15,30,5), #seq(lower limit, upper limit, tick frequency)
    expand = c(0,0)) +
  labs(y = "SSMD",
       x = "Drug-dose") +
  theme(
    axis.text.x = element_text(face = 'bold'), 
    axis.text.y = element_text(face = 'bold'), 
    axis.title.x = element_text(face = 'bold'),
    axis.title.y = element_text(face = 'bold'),
    legend.title = element_text(face = 'bold'),
    legend.text = element_text(face = 'bold'),
    strip.text.x = element_text(face = 'bold')) 
      
ggsave("SSMD_manhattan.pdf", width = 4, height = 3, path = "./output_data/")
```

## Dual flashlight plot

```{r}
ggplot(data = JHDL_summary_nocontrol,
       aes(x = log2_Average_Fold_Change,
           y = hsSSMD,
           color = ifelse(((log2_Average_Fold_Change<=-1.5 & hsSSMD<=-1)),"A", "B"),
           shape = ifelse(((log2_Average_Fold_Change<=-1.5 & hsSSMD<=-1)),"A", "B"))) + 
  geom_point() +
  theme_classic() +
  scale_shape_manual(guide = "none", 
                     values = c(19,1)) +
  scale_color_manual(guide = "none", 
                     values = c("red", "black")) +
  geom_hline(yintercept = c(-1,1), 
             linetype="dashed") +
  geom_hline(yintercept = 0) +
  geom_vline(xintercept = c(-1.5,1.5), 
             linetype="dashed") +
  geom_vline(xintercept = 0) +
  geom_label_repel(
    data = JHDL_summary_nocontrol %>% filter(Compound_name == "Enoxolone"),
    aes(label = paste(Dose, Compound_name)),
    color = "black",
    nudge_x = 3, nudge_y = -6) + 
  scale_y_continuous(
    name = "SSMD",
    labels = comma, limits = c(-15,30),
    breaks = c(-15,-10,-5,-1,0,1,5,10,15,20,25,30),
    expand = c(0,0)) + 
  scale_x_continuous(
    name = "Fold Change (log2)",
    limits = c(-6,6),
    breaks = c(-5,-3,-1.5,0,1.5,5,3),
    expand = c(0,0)) +
  theme(
    axis.text.x = element_text(face = 'bold'), 
    axis.text.y = element_text(face = 'bold'), 
    axis.title.x = element_text(face = 'bold'),
    axis.title.y = element_text(face = 'bold'),
    legend.title = element_text(face = 'bold'),
    legend.text = element_text(face = 'bold'),
    strip.text.x = element_text(face = 'bold')) 

ggsave("dual_flashlight_labeledEnoxo.svg", width = 6, height = 6, path = "./output_data/")
```

## JHDL positive control results

```{r}
#not sure what to do here... boxplot? violin plot? histogram/density?
#relative results - what does this look like for fold change and/or SSMD
#all control dual flashlight

y_lim <- JHDL_summary_controls %>% 
  pull(Average_RLU) %>% 
  max(.) %>% 
  round_any(., 50000, ceiling)

#boxplot
ggplot(data = JHDL_summary_controls,
       aes(x = factor(Drug, 
                      levels = c("Vehicle", 
                                 "Lomitapide")),
           y = Average_RLU)) +
  theme_classic() +
  geom_boxplot(outlier.shape = NA) +
  geom_quasirandom(alpha = 0.03) +
  scale_y_continuous(
    name = "Relative Luminescence Units (RLU)",
    labels = comma, limits = c(0,y_lim), #c(lower limit, upper limit)
    breaks = seq(0,y_lim,y_lim/8)) +
  scale_x_discrete(labels=c("Lomitapide" = "5 µM lomitapide")) +
  theme(
    axis.text.x = element_text(face = 'bold', angle = 45, hjust = 1), 
    axis.text.y = element_text(face = 'bold'), 
    axis.title.x = element_blank(),
    axis.title.y = element_text(face = 'bold'),
    legend.title = element_text(face = 'bold'),
    legend.text = element_text(face = 'bold'),
    strip.text.x = element_text(face = 'bold')) 

ggsave("JHDL_controls.pdf", width = 2, height = 4, path = "./output_data/")
```

```{r}
JHDL_summary_controls %>% 
  select(Drug, Average_RLU) %>%
  group_by(Drug) %>%
  summarize(mean_of_RLU = mean(Average_RLU))

JHDL_summary_controls %>% 
  select(Drug, Average_RLU) %>%
  mutate(fold_change = Average_RLU/1013062.4) %>%
  group_by(Drug) %>%
  summarize(mean = mean(fold_change),
            sd = sd(fold_change))
```

```{r}
ordered_JHDL_summary_controls <- JHDL_summary_controls %>% 
  filter(Drug != "Vehicle") %>% 
  arrange(-hsSSMD) %>% 
  mutate(compound_number = seq(1:nrow(.))) %>%
  select(compound_number, hsSSMD)

fit <- lm(hsSSMD ~ poly(compound_number, 3), 
   data = ordered_JHDL_summary_controls)

cc <- check_curve(ordered_JHDL_summary_controls$hsSSMD,
                  ordered_JHDL_summary_controls$compound_number) 

ip <- bese(ordered_JHDL_summary_controls$hsSSMD, 
           ordered_JHDL_summary_controls$compound_number, 
           cc$index)
  
ggplot(data = ordered_JHDL_summary_controls,
       aes(x = compound_number, 
           y = hsSSMD)) +
  geom_point() +
  theme_classic() +
  geom_hline(yintercept = 0) +
  geom_hline(yintercept = -1, linetype = "dashed") +
  geom_hline(yintercept = ip$iplast,color = "blue") +
#  stat_smooth(method = "lm",
#              formula = y ~ poly(x, 3),
#              color = "red") +
  scale_x_continuous(limits = c(0,1400),
                     breaks = seq(0,1500,250)) +
  labs(x = str_wrap("Position in the ranked list of lomitapide SSMD measurements", 
                    width = 40),
       y = "SSMD") +
  theme(
    axis.text.x = element_text(face = 'bold'), 
    axis.text.y = element_text(face = 'bold'), 
    axis.title.x = element_text(face = 'bold'),
    axis.title.y = element_text(face = 'bold')) 

ggsave("ranked_lomi_SSMD.pdf", width = 4, height = 4, path = "./output_data/")
```

```{r}
ordered_JHDL_summary_controls <- JHDL_summary_controls %>% 
  filter(Drug != "Vehicle") %>% 
  arrange(-log2_Average_Fold_Change) %>% 
  mutate(compound_number = seq(1:nrow(.))) %>%
  select(compound_number, log2_Average_Fold_Change)

fit <- lm(log2_Average_Fold_Change ~ poly(compound_number, 3), 
   data = ordered_JHDL_summary_controls)

cc <- check_curve(ordered_JHDL_summary_controls$log2_Average_Fold_Change,
                  ordered_JHDL_summary_controls$compound_number) 

ip <- bese(ordered_JHDL_summary_controls$log2_Average_Fold_Change, 
           ordered_JHDL_summary_controls$compound_number, 
           cc$index)
  
ggplot(data = ordered_JHDL_summary_controls,
       aes(x = compound_number, 
           y = log2_Average_Fold_Change)) +
  geom_point() +
  theme_classic() +
  geom_hline(yintercept = 0) +
  geom_hline(yintercept = -1.5, linetype = "dashed") +
  geom_hline(yintercept = ip$iplast,color = "blue") +
#  stat_smooth(method = "lm",
#              formula = y ~ poly(x, 3),
#              color = "red") +
  scale_x_continuous(limits = c(0,1400),
                     breaks = seq(0,1500,250)) +
  labs(x = str_wrap("Position in the ranked list of lomitapide fold change measurements", 
                    width = 40),
       y = "Fold change (log2)") +
  theme(
    axis.text.x = element_text(face = 'bold'), 
    axis.text.y = element_text(face = 'bold'), 
    axis.title.x = element_text(face = 'bold'),
    axis.title.y = element_text(face = 'bold')) 

ggsave("ranked_lomi_foldchange.pdf", width = 4, height = 4, path = "./output_data/")
```

## Table of hits from JHDL

```{r}
Hit_list <- JHDL_summary_nocontrol %>% 
  filter(hsSSMD < -1 & log2_Average_Fold_Change < -1.5) %>%
  select(Compound_name) %>%
  unique() %>%
  arrange(Compound_name) 

Hit_list <- Hit_list %>%
  add_column(Compound_ID = seq(1:nrow(Hit_list)))

Hit_list_dose <- JHDL_summary_nocontrol %>% 
  filter(hsSSMD < -1 & log2_Average_Fold_Change < -1.5) %>%
  select(Compound_name, Dose, Filename) %>%
  arrange(Compound_name) %>% 
  right_join(Hit_list)
```

# Supplemental Figure 1

## Raw results of hits from JHDL

```{r}
dir.create("./JHDL_primary_result", showWarnings = FALSE)

JHDL_hit_data <- JHDL_all_data[0,]

Define_y_limit <- function(x) {
  y_lim <<- x %>%
    filter(Outlier != "Outlier") %>% 
    pull(RLU) %>% 
    max(.) %>%
    round_any(.,50000, ceiling)}

Define_outliers <- function(x) {
  Outliers <<- x %>% 
    group_by(Dose) %>% 
    mutate(RLU_Q1 = (quantile(RLU, na.rm = TRUE)[[2]]), 
           RLU_Q3 = (quantile(RLU, na.rm = TRUE)[[4]]), 
           avg = mean(RLU, na.rm = TRUE),
           RLU_IQR = IQR(RLU, na.rm = TRUE),
           Outlier = if_else(RLU > RLU_Q3+1.5*(RLU_IQR) | RLU < RLU_Q1-1.5*(RLU_IQR), 
                             "Outlier",
                             "Normal"))}

plot_data <- function(x) {
  m <<- ggplot(data = x, 
               mapping = aes(x = factor(Dose, 
                                        levels = c(
                                          "Vehicle",
                                          "5 µM",
                                          "1 µM",
                                          "2 µM",
                                          "4 µM",
                                          "8 µM")), 
                 y = RLU)) +
    geom_boxplot(outlier.shape = NA) +
    geom_jitter(width = 0.2, shape = 1, alpha = 0.6) +
    theme_bw() +
    scale_y_continuous(
      name = "Relative Luminescence Units (RLU)",
      labels = comma, limits = c(0,y_lim), #c(lower limit, upper limit)
      breaks = seq(0,y_lim,y_lim/8)) +  #seq(lower limit, upper limit, tick frequency)
    theme(
      axis.text.x = element_text(face = 'bold', angle = 45, hjust = 1), 
      axis.title.x = element_blank(),
      axis.text.y = element_text(face = 'bold'), 
      axis.title.y = element_text(face = 'bold'),
      legend.title = element_text(face = 'bold'),
      legend.text = element_text(face = 'bold'),
      strip.text.x = element_text(face = 'bold')) 
}

for (i in Hit_list$Compound_name) {
  if (nrow(JHDL_all_data %>% filter(Compound_name == i)) == 0) {next}
  
  dir.create(path = paste("./JHDL_primary_result/",i, sep = ""), showWarnings = FALSE)
  
  Plate_name <- JHDL_all_data %>% 
    filter(Compound_name == i) %>% 
    pull(Filename) %>%
    unique()
  
  JHDL_all_data.i <- JHDL_all_data %>% 
    filter(Filename %in% Plate_name) %>% 
    filter(Drug == "Vehicle" | Drug == "Lomitapide" | Compound_name == i)
  
  for (a in JHDL_all_data.i %>% distinct(Filename) %>% pull(Filename)) {
    JHDL_all_data.i.a <- JHDL_all_data.i %>% filter(Filename == a)
    Define_outliers(JHDL_all_data.i.a)
    Outlier.count <- Outliers %>% filter(Outlier == "Outlier") %>% nrow(.)
    Define_y_limit(Outliers)
    
    #write to new data frame
    JHDL_hit_data <- JHDL_hit_data %>% 
      rbind(Outliers)
    
    sink(file = paste("./JHDL_primary_result/",i,"/",i,"_",a,".txt", sep = ""))
    cat("Sample sizes\n")
    Outliers %>% 
      filter(Outlier == "Normal") %>%
      group_by(Drug, Dose) %>%
      summarize(n = n()) %>%
      print.data.frame()      
    cat("\nOutliers removed\n")
    Outliers %>% 
      filter(Outlier != "Normal") %>% 
      select(c(Row, Column, Dose, Drug)) %>% 
      print.data.frame()
    sink(file = NULL)
    
    plot_data(Outliers %>% filter(Outlier == "Normal"))
    ggsave(m, 
           filename = paste(i,"_", a, ".pdf", sep = ""), 
           width = 4, 
           height = 5, 
           path = paste("./JHDL_primary_result/",i, sep = ""))
  }
}
```

# Figure 2

## Summary results of hits from JHDL

```{r}
#change drugs names to letters and control
JHDL_hit_only_data <- JHDL_hit_data %>%
  filter(Drug != "Lomitapide") %>%
  filter(Drug != "Vehicle") %>%
  right_join(Hit_list_dose, by = c("Compound_name", "Filename", "Dose"))

JHDL_hit_control_data <- JHDL_hit_data %>%
  filter(Drug == "Lomitapide" | Drug == "Vehicle") %>%
  mutate(Compound_name = "Control")

JHDL_hit_data_clean <- JHDL_hit_only_data %>%
  rbind(JHDL_hit_control_data) %>% 
  mutate(Dose = case_match(Dose, 
                           "5 µM" ~ "5 µM lomitapide",
                           .default = Dose))

JHDL_hit_data_clean$Compound_ID <- JHDL_hit_data_clean$Compound_ID %>% 
  as.character() %>% 
  replace_na("Control")

JHDL_hit_data_clean <- JHDL_hit_data_clean %>%
  left_join(Hit_list %>% mutate(Compound_ID = as.character(Compound_ID)), 
            by = "Compound_ID") %>%
  mutate(Compound_label = ifelse(Compound_ID == "Control", "Control", Compound_name.y),
         Compound_label = case_match(Compound_label,
           "Cytidine 5'-diphosphate trisodium salt" ~ "CDP",
           "Cytidine 5'-monophosphate" ~ "CMP",
           .default = Compound_label))

JHDL_hit_data_clean_1 <- JHDL_hit_data_clean %>%
  filter(Compound_ID %in% c("Control", "1", "2", "3", "4", "5", "6", "7", "8", "9", "10", "11", "12", "13", "14", "15", "16", "17", "18", "19", "20", "21", "22", "23", "24", "25"))
         
JHDL_hit_data_clean_2 <- JHDL_hit_data_clean %>%
  filter(Compound_ID %in% c("Control", "26", "27", "28", "29", "30", "31", "32", "33", "34", "35", "36", "37", "38", "39", "40", "41", "42", "43", "44", "45", "46", "47", "48", "49", "50"))

label_order <- c("Control", Hit_list %>% arrange(Compound_ID) %>% pull(Compound_name)) %>%
  str_replace("Cytidine 5'-diphosphate trisodium salt", "CDP") %>%
  str_replace("Cytidine 5'-monophosphate", "CMP")


ggplot(data = JHDL_hit_data_clean_1 %>% filter(Outlier == "Normal"), 
       mapping = aes(x = factor(Dose,
                                levels = c("Vehicle",
                                           "5 µM lomitapide",
                                           "1 µM",
                                           "2 µM",
                                           "4 µM",
                                           "8 µM")), 
                     y = Fold_Change)) +
  geom_jitter(width = 0.2, alpha = 0.1) +
  geom_boxplot(outlier.shape = NA, alpha = 0.5) +
  theme_bw() +
  theme(panel.grid.major = element_blank(),
        panel.grid.minor = element_blank()) +
  geom_hline(yintercept = 1, linetype = "dashed") +
  scale_y_continuous(
    name = "Fold change (relative to respective vehicle control)",
    labels = comma, limits = c(0,2.2), #c(lower limit, upper limit)
    breaks = seq(0,2.2,0.2)) +  #seq(lower limit, upper limit, tick frequency)
  theme(axis.text.x = element_text(face = 'bold', 
                                   angle = 45, 
                                   hjust = 1), 
        axis.title.x = element_blank(),
        axis.text.y = element_text(face = 'bold'), 
        axis.title.y = element_text(face = 'bold'),
        legend.title = element_text(face = 'bold'),
        legend.text = element_text(face = 'bold'),
        strip.text.x = element_text(face = 'bold', angle = 90, hjust = 0),
        panel.spacing = unit(0, "line")) +
  facet_grid(~factor(Compound_label, levels = label_order),
           space = "free",
           scales = "free")

ggsave("JHDL_hit_summary_boxplot_1.svg", 
       width = 12, height = 6, path = "./Output_data/")

ggplot(data = JHDL_hit_data_clean_2 %>% filter(Outlier == "Normal"), 
       mapping = aes(x = factor(Dose,
                                levels = c("Vehicle",
                                           "5 µM lomitapide",
                                           "1 µM",
                                           "2 µM",
                                           "4 µM",
                                           "8 µM")), 
                     y = Fold_Change)) +
  geom_jitter(width = 0.2, alpha = 0.1) +
  geom_boxplot(outlier.shape = NA, alpha = 0.5) +
  theme_bw() +
  theme(panel.grid.major = element_blank(),
        panel.grid.minor = element_blank()) +
  geom_hline(yintercept = 1, linetype = "dashed") +
  scale_y_continuous(
    name = "Fold change (relative to respective vehicle control)",
    labels = comma, limits = c(0,2.2), #c(lower limit, upper limit)
    breaks = seq(0,2.2,0.2)) +  #seq(lower limit, upper limit, tick frequency)
  theme(axis.text.x = element_text(face = 'bold', 
                                   angle = 45, 
                                   hjust = 1), 
        axis.title.x = element_blank(),
        axis.text.y = element_text(face = 'bold'), 
        axis.title.y = element_text(face = 'bold'),
        legend.title = element_text(face = 'bold'),
        legend.text = element_text(face = 'bold'),
        strip.text.x = element_text(face = 'bold', angle = 90, hjust = 0),
        panel.spacing = unit(0, "line")) +
  facet_grid(~factor(Compound_label, levels = label_order),
           space = "free",
           scales = "free")

ggsave("JHDL_hit_summary_boxplot_2.svg", 
       width = 12, height = 6, path = "./Output_data/")
```

# Supplemental Figure 2

## Raw validation of hits from JHDL

```{r}
dir.create("./JHDL_validation_result", showWarnings = FALSE)

Validation <- read_plate("JHDL_validation_raw/Validation_summary.csv", 
                         well_ids_column = "Wells")

Validation <- Validation %>% 
  mutate(across(c(3:ncol(.)), as.integer)) %>%
  pivot_longer(cols = c(3:ncol(.)), names_to = "Drug", values_to = "RLU") %>%
  separate(col = "Drug", sep = "-", into = c("Experiment_name", "Drug_name"))  %>%
  filter(Treatment != "Empty")

Drug_list <- Validation %>% 
  pull(Drug_name) %>% 
  unique()

##round_any() is a function in the package plyr, but plyr has some 
##compatibility issues with the tidyverse, thus just writing in the 
##function manually
round_any = function(x, accuracy, f=round){f(x/ accuracy) * accuracy}

Define_outliers <- function(x) {
    Validation.outliers <<- x %>% 
    filter(Drug_name == i) %>% 
    group_by(Treatment) %>% 
    mutate(RLU_Q1 = (quantile(RLU, na.rm = TRUE)[[2]]), 
           RLU_Q3 = (quantile(RLU, na.rm = TRUE)[[4]]), 
           avg = mean(RLU, na.rm = TRUE),
           RLU_IQR = IQR(RLU, na.rm = TRUE),
           Outlier = if_else(RLU > RLU_Q3+1.5*(RLU_IQR) | RLU < RLU_Q1-1.5*(RLU_IQR), "Outlier", "Normal"))}

Define_y_limit <- function(x) {
  y_lim <<- x %>%
    filter(Outlier != "Outlier") %>% 
    pull(RLU) %>% 
    max(.) %>%
    round_any(.,50000, ceiling)}

plot_validation_data <- function(x) {
  m <<- ggplot(data = x, 
                 mapping = aes(x = factor(Treatment, levels = c(
                   "Vehicle",
                   "5 µM lomitapide",
                   "0.0625 µM",
                   "0.125 µM",
                   "0.25 µM",
                   "0.5 µM",
                   "1 µM",
                   "2 µM",
                   "4 µM",
                   "8 µM")), 
                   y = RLU)) +
    geom_boxplot() +
    geom_point() +
    scale_y_continuous(
      name = "Relative Luminescence Units (RLU)",
      labels = comma, limits = c(0,y_lim), #c(lower limit, upper limit)
      breaks = seq(0,y_lim,y_lim/10)) +  #seq(lower limit, upper limit, tick frequency)
    theme(
      axis.text.x = element_text(face = 'bold', angle = 45, hjust = 1), 
      axis.title.x = element_blank(),
      axis.text.y = element_text(face = 'bold'), 
      axis.title.y = element_text(face = 'bold'),
      legend.title = element_text(face = 'bold'),
      legend.text = element_text(face = 'bold'),
      strip.text.x = element_text(face = 'bold')) +
    labs(title = paste("Validation of ",i, sep=""), 
         subtitle = paste(Outlier.count, " total outliers removed", sep = ""),
         caption = paste("Experiment: ", a, sep = ""))
}

#generate relative data to combine into a single graph
for (i in Drug_list) {
  if (nrow(Validation %>% filter(Drug_name == i)) == 0) {next}
  
  Validation.i <- Validation %>% filter(Drug_name == i)
  
  for (a in Validation.i %>% distinct(Experiment_name) %>% pull(Experiment_name)) {
    Validation.i.a <- Validation.i %>% filter(Experiment_name == a)
    Define_outliers(Validation.i.a)
    Outlier.count <- Validation.outliers %>% filter(Outlier == "Outlier") %>% nrow(.)
    Define_y_limit(Validation.outliers)
    
    plot_validation_data(Validation.outliers)
    ggsave(m, filename = paste(i,"_", a, ".pdf", sep = ""), width = 4, height = 5, 
           path = "JHDL_validation_result/")
  }
}
```

## Relative validation of hits from JHDL

```{r}
dir.create("./JHDL_validation_result/Relative_data", showWarnings = FALSE) 

Relative_validation <- Validation %>% 
  group_by(Experiment_name, Drug_name, Treatment) %>%
  summarise(vehicle_mean = mean(RLU)) %>%
  filter(Treatment == "Vehicle") %>%
  select(-Treatment) %>%
  right_join(Validation, by = join_by(Experiment_name, Drug_name)) %>%
  mutate(fold_change = RLU/vehicle_mean) %>%
  group_by(Experiment_name, Drug_name, Treatment) %>%
  mutate(fold_change_Q1 = (quantile(fold_change, na.rm = TRUE)[[2]]), 
         fold_change_Q3 = (quantile(fold_change, na.rm = TRUE)[[4]]), 
         fold_change_IQR = IQR(fold_change, na.rm = TRUE),
         Outlier = if_else(fold_change > fold_change_Q3+1.5*(fold_change_IQR) |
                             fold_change < fold_change_Q1-1.5*(fold_change_IQR),
                           "Outlier", 
                           "Normal"))

for (i in Drug_list) {
  y_lim <- Relative_validation %>% 
    filter(Drug_name == i,
           Outlier == "Normal") %>%
    pull(fold_change) %>% 
    max(.) %>%
    round_any(.,0.5, ceiling)
  
  p <- ggplot(data = Relative_validation %>% 
           filter(Drug_name == i,
                  Outlier == "Normal"),
         aes(x = factor(Treatment, levels = c("Vehicle",
                                            "5 µM lomitapide",
                                            "0.0625 µM",
                                            "0.125 µM",
                                            "0.25 µM",
                                            "0.5 µM",
                                            "1 µM",
                                            "2 µM",
                                            "4 µM",
                                            "8 µM")), 
             y = fold_change)) +
    geom_boxplot(outlier.shape = NA) +
    geom_jitter(width = 0.2, shape = 1, alpha = 0.6) +
    theme_bw() +
    scale_y_continuous(
      labels = comma, limits = c(0,y_lim), #c(lower limit, upper limit)
      breaks = seq(0,y_lim,y_lim/8)) +  #seq(lower limit, upper limit, tick frequency)
    labs(x = NULL,
         y = str_wrap("Fold chance in luminescense (relative to vehicle)", 
                      width = 30)) +
    theme(
      axis.text.x = element_text(face = 'bold', angle = 90, hjust = 1, vjust = 0.5), 
      axis.title.x = element_text(face = 'bold'),
      axis.text.y = element_text(face = 'bold'), 
      axis.title.y = element_text(face = 'bold'),
      legend.title = element_text(face = 'bold'),
      legend.text = element_text(face = 'bold'),
      strip.text.x = element_text(face = 'bold'))
  
  ggsave(filename = paste0(i, ".pdf"), 
         plot = p,
         width = 4, 
         height = 4, 
         path = "./JHDL_validation_result/Relative_data")
  
  sink(file = paste0("./JHDL_validation_result/Relative_data/", i, ".txt"))
  cat("Sample sizes\n")
  Relative_validation %>% 
    filter(Drug_name == i,
           Outlier == "Normal") %>%
    group_by(Treatment) %>%
    summarize(n = n()) %>%
    print.data.frame() 
  cat("\nOutliers removed\n")
  Relative_validation %>% 
    filter(Drug_name == i,
           Outlier != "Normal") %>%
    group_by(Treatment) %>%
    summarize(n = n()) %>%
    print.data.frame() 
  cat("\nANOVA results\n")
  Relative_validation %>% 
      filter(Drug_name == i,
             Outlier == "Normal") %>%
      ungroup() %>%
      select(Treatment, fold_change) %>%
      compare_means(formula = fold_change~Treatment, 
                    method = "anova") %>%
  print.data.frame()
  cat("\nPairwise t test results\n")
  Relative_validation %>% 
          filter(Drug_name == i,
                 Outlier == "Normal") %>% 
          ungroup() %>%
          select(Treatment, fold_change) %>%
          compare_means(formula = fold_change~Treatment,
                        ref.group = "Vehicle", 
                        p.adjust.method = "bonf", 
                        method = "t.test") %>% 
          arrange(factor(group2, levels = c(
                          "5 µM lomitapide",
                          "0.0625 µM",
                          "0.125 µM",
                          "0.25 µM",
                          "0.5 µM",
                          "1 µM",
                          "2 µM",
                          "4 µM",
                          "8 µM"))) %>%
    print.data.frame()
  cat("\nns: p > 0.05\n*: p <= 0.05\n**: p <= 0.01\n***: p <= 0.001\n****: p <= 0.0001\n")
  sink(file = NULL)
}
```

# Figure 3 - Drug validation

## Calcipotriene

```{r}
#| label: LipoGlo assay with homogenization endpoint of Calcipotriene treated samples
drug_name = "Calcipotriene"

drug_hom <- read_plate("Figure_3_data/homogenate/Clp_lpCount.csv", well_ids_column = "Wells")
drug_hom <- drug_hom %>%
  na.omit() %>%
  filter(Treatment != "Empty") %>%
  pivot_longer(cols = c("P1", "P2", "P3", "P4"), names_to = "Replicate", values_to = "RLU")

drug_hom_veh_summary <- drug_hom %>% 
  group_by(Replicate, Treatment) %>% 
  summarise(vehicle_mean = mean(RLU, na.rm = TRUE)) %>% 
  filter(Treatment == "Vehicle") %>%
  select(-Treatment)

drug_hom <- drug_hom %>% 
  full_join(drug_hom_veh_summary) %>%
  mutate(fold_change = RLU/vehicle_mean)

drug_hom <- drug_hom %>% 
  group_by(Treatment) %>%
  mutate(fold_change_Q1 = (quantile(fold_change, na.rm = TRUE)[[2]]), 
         fold_change_Q3 = (quantile(fold_change, na.rm = TRUE)[[4]]), 
         fold_change_IQR = IQR(fold_change, na.rm = TRUE),
         Outlier = if_else(fold_change > fold_change_Q3+1.5*(fold_change_IQR) |
                             fold_change < fold_change_Q1-1.5*(fold_change_IQR),
                           "Outlier", 
                           "Normal"))

y_lim <- drug_hom %>% 
  filter(Outlier == "Normal") %>%
  pull(fold_change) %>% 
  max(.) %>% 
  round_any(., 1, ceiling)

ggplot(data = drug_hom %>% 
  filter(Outlier == "Normal"),
       aes(x = factor(Treatment, levels = c("Vehicle",
                                            "5 µM lomitapide",
                                            "4 µM",
                                            "8 µM")), 
           y = fold_change)) +
  geom_boxplot(outlier.shape = NA) +
  geom_jitter(width = 0.2, shape = 1, alpha = 0.6) +
  theme_bw() +
  scale_y_continuous(
    labels = comma, limits = c(0,1.75), #c(lower limit, upper limit)
    breaks = seq(0,1.75,0.25)) +  #seq(lower limit, upper limit, tick frequency)
  labs(x = "Treatment",
       y = str_wrap("Fold chance in luminescense (relative to vehicle)", 
                    width = 30)) +
  theme(
    axis.text.x = element_text(face = 'bold', angle = 45, hjust = 1), 
    axis.title.x = element_text(face = 'bold'),
    axis.text.y = element_text(face = 'bold'), 
    axis.title.y = element_text(face = 'bold'),
    legend.title = element_text(face = 'bold'),
    legend.text = element_text(face = 'bold'),
    strip.text.x = element_text(face = 'bold'))
ggsave(paste0(drug_name, "_hom.svg"), width = 3, height = 4, path = "./output_data/")

drug_hom %>%
  filter(Outlier == "Normal") %>%
  group_by(Treatment) %>%
  summarise(n = n())

drug_hom %>% 
  filter(Outlier == "Normal") %>%
  compare_means(formula = fold_change~Treatment,
                ref.group = "Vehicle", 
                p.adjust.method = "bonf", 
                method = "t.test")

aov_drug_hom <- aov(data = drug_hom %>% filter(Outlier == "Normal"), 
                   formula = fold_change ~ Treatment)
summary(aov_drug_hom)

# nativePAGE
drug_PAGE <- read.csv(file = "Figure_3_data/nativePAGE/Clp_PAGEquant.csv")

drug_PAGE <- drug_PAGE %>% 
  pivot_longer(cols = c("ZM", "VLDL", "IDL", "LDL"),
               names_to = "LP_class", 
               values_to = "Bin_value")

ggplot(data = drug_PAGE,
       aes(x = factor(Treatment, c("Vehicle",
                                   "5 µM lomitapide",
                                   "4 µM",
                                   "8 µM")),
           y = Bin_value*100)) +
  geom_boxplot(outlier.shape = NA) +
  geom_jitter(width = 0.2, shape = 1, alpha = 0.6) +
  theme_bw() +
  scale_y_continuous(
    labels = comma, limits = c(0,80), #c(lower limit, upper limit)
    breaks = seq(0,80,10)) +  #seq(lower limit, upper limit, tick frequency)
  labs(x = "Treatment",
       y = "Proportion of total lipoproteins (%)") +
  theme(
    axis.text.x = element_text(face = 'bold', angle = 45, hjust = 1), 
    axis.title.x = element_text(face = 'bold'),
    axis.text.y = element_text(face = 'bold'), 
    axis.title.y = element_text(face = 'bold'),
    legend.title = element_text(face = 'bold'),
    legend.text = element_text(face = 'bold'),
    strip.text.x = element_text(face = 'bold')) +
  facet_grid(~factor(LP_class, c("ZM", "VLDL", "IDL", "LDL")))
ggsave(paste0(drug_name, "_nPAGE.svg"), width = 5, height = 4, path = "./output_data/")

drug_PAGE %>%
  group_by(Treatment, LP_class) %>%
  summarise(n = n())

means_comp <- aov(data = drug_PAGE %>% filter(LP_class == "ZM"), formula = Bin_value ~ Treatment)
summary(means_comp)
Anova(means_comp, type = "3")

drug_PAGE %>% 
  filter(LP_class == "ZM") %>%
  compare_means(formula = Bin_value~Treatment,
                ref.group = "Vehicle", 
                p.adjust.method = "bonf", 
                method = "t.test")

means_comp <- aov(data = drug_PAGE %>% filter(LP_class == "VLDL"), formula = Bin_value ~ Treatment)
summary(means_comp)
Anova(means_comp, type = "3")

drug_PAGE %>% 
  filter(LP_class == "VLDL") %>%
  compare_means(formula = Bin_value~Treatment,
                ref.group = "Vehicle", 
                p.adjust.method = "bonf", 
                method = "t.test")

means_comp <- aov(data = drug_PAGE %>% filter(LP_class == "IDL"), formula = Bin_value ~ Treatment)
summary(means_comp)
Anova(means_comp, type = "3")

drug_PAGE %>% 
  filter(LP_class == "IDL") %>%
  compare_means(formula = Bin_value~Treatment,
                ref.group = "Vehicle", 
                p.adjust.method = "bonf", 
                method = "t.test")

means_comp <- aov(data = drug_PAGE %>% filter(LP_class == "LDL"), formula = Bin_value ~ Treatment)
summary(means_comp)
Anova(means_comp, type = "3")

drug_PAGE %>% 
  filter(LP_class == "LDL") %>%
  compare_means(formula = Bin_value~Treatment,
                ref.group = "Vehicle", 
                p.adjust.method = "bonf", 
                method = "t.test")
```

## Doxycycline

```{r}
#| label: LipoGlo assay with homogenization endpoint of Doxycycline treated samples
drug_name = "Doxycycline"

drug_hom <- read_plate("Figure_3_data/homogenate/Dox_lpCount.csv", well_ids_column = "Wells")
drug_hom <- drug_hom %>%
  na.omit() %>%
  filter(Treatment != "Empty") %>%
  pivot_longer(cols = c("P1", "P2", "P3"), names_to = "Replicate", values_to = "RLU")

drug_hom_veh_summary <- drug_hom %>% 
  group_by(Replicate, Treatment) %>% 
  summarise(vehicle_mean = mean(RLU, na.rm = TRUE)) %>% 
  filter(Treatment == "Vehicle") %>%
  select(-Treatment)

drug_hom <- drug_hom %>% 
  full_join(drug_hom_veh_summary) %>%
  mutate(fold_change = RLU/vehicle_mean)

drug_hom <- drug_hom %>% 
  group_by(Treatment) %>%
  mutate(fold_change_Q1 = (quantile(fold_change, na.rm = TRUE)[[2]]), 
         fold_change_Q3 = (quantile(fold_change, na.rm = TRUE)[[4]]), 
         fold_change_IQR = IQR(fold_change, na.rm = TRUE),
         Outlier = if_else(fold_change > fold_change_Q3+1.5*(fold_change_IQR) |
                             fold_change < fold_change_Q1-1.5*(fold_change_IQR),
                           "Outlier", 
                           "Normal"))

y_lim <- drug_hom %>% 
  filter(Outlier == "Normal") %>%
  pull(fold_change) %>% 
  max(.) %>% 
  round_any(., 1, ceiling)

ggplot(data = drug_hom %>% 
  filter(Outlier == "Normal"),
       aes(x = factor(Treatment, levels = c("Vehicle",
                                            "5 µM lomitapide",
                                            "4 µM",
                                            "8 µM")), 
           y = fold_change)) +
  geom_boxplot(outlier.shape = NA) +
  geom_jitter(width = 0.2, shape = 1, alpha = 0.6) +
  theme_bw() +
  scale_y_continuous(
    labels = comma, limits = c(0,1.75), #c(lower limit, upper limit)
    breaks = seq(0,1.75,0.25)) +  #seq(lower limit, upper limit, tick frequency)
  labs(x = "Treatment",
       y = str_wrap("Fold chance in luminescense (relative to vehicle)", 
                    width = 30)) +
  theme(
    axis.text.x = element_text(face = 'bold', angle = 45, hjust = 1), 
    axis.title.x = element_text(face = 'bold'),
    axis.text.y = element_text(face = 'bold'), 
    axis.title.y = element_text(face = 'bold'),
    legend.title = element_text(face = 'bold'),
    legend.text = element_text(face = 'bold'),
    strip.text.x = element_text(face = 'bold'))
ggsave(paste0(drug_name, "_hom.svg"), width = 3, height = 4, path = "./output_data/")

drug_hom %>%
  filter(Outlier == "Normal") %>%
  group_by(Treatment) %>%
  summarise(n = n())

drug_hom %>% 
  filter(Outlier == "Normal") %>%
  compare_means(formula = fold_change~Treatment,
                ref.group = "Vehicle", 
                p.adjust.method = "bonf", 
                method = "t.test")

aov_drug_hom <- aov(data = drug_hom %>% filter(Outlier == "Normal"), 
                   formula = fold_change ~ Treatment)
summary(aov_drug_hom)

# nativePAGE
drug_PAGE <- read.csv(file = "Figure_3_data/nativePAGE/Dox_PAGEquant_cleanLadder.csv")

drug_PAGE <- drug_PAGE %>% 
  pivot_longer(cols = c("ZM", "VLDL", "IDL", "LDL"),
               names_to = "LP_class", 
               values_to = "Bin_value")

ggplot(data = drug_PAGE,
       aes(x = factor(Treatment, c("Vehicle",
                                   "5 µM lomitapide",
                                   "4 µM",
                                   "8 µM")),
           y = Bin_value*100)) +
  geom_boxplot(outlier.shape = NA) +
  geom_jitter(width = 0.2, shape = 1, alpha = 0.6) +
  theme_bw() +
  scale_y_continuous(
    labels = comma, limits = c(0,80), #c(lower limit, upper limit)
    breaks = seq(0,80,10)) +  #seq(lower limit, upper limit, tick frequency)
  labs(x = "Treatment",
       y = "Proportion of total lipoproteins (%)") +
  theme(
    axis.text.x = element_text(face = 'bold', angle = 45, hjust = 1), 
    axis.title.x = element_text(face = 'bold'),
    axis.text.y = element_text(face = 'bold'), 
    axis.title.y = element_text(face = 'bold'),
    legend.title = element_text(face = 'bold'),
    legend.text = element_text(face = 'bold'),
    strip.text.x = element_text(face = 'bold')) +
  facet_grid(~factor(LP_class, c("ZM", "VLDL", "IDL", "LDL")))
ggsave(paste0(drug_name, "_nPAGE.svg"), width = 5, height = 4, path = "./output_data/")

drug_PAGE %>%
  group_by(Treatment, LP_class) %>%
  summarise(n = n())

means_comp <- aov(data = drug_PAGE %>% filter(LP_class == "ZM"), formula = Bin_value ~ Treatment)
summary(means_comp)
Anova(means_comp, type = "3")

drug_PAGE %>% 
  filter(LP_class == "ZM") %>%
  compare_means(formula = Bin_value~Treatment,
                ref.group = "Vehicle", 
                p.adjust.method = "bonf", 
                method = "t.test")

means_comp <- aov(data = drug_PAGE %>% filter(LP_class == "VLDL"), formula = Bin_value ~ Treatment)
summary(means_comp)
Anova(means_comp, type = "3")

drug_PAGE %>% 
  filter(LP_class == "VLDL") %>%
  compare_means(formula = Bin_value~Treatment,
                ref.group = "Vehicle", 
                p.adjust.method = "bonf", 
                method = "t.test")

means_comp <- aov(data = drug_PAGE %>% filter(LP_class == "IDL"), formula = Bin_value ~ Treatment)
summary(means_comp)
Anova(means_comp, type = "3")

drug_PAGE %>% 
  filter(LP_class == "IDL") %>%
  compare_means(formula = Bin_value~Treatment,
                ref.group = "Vehicle", 
                p.adjust.method = "bonf", 
                method = "t.test")

means_comp <- aov(data = drug_PAGE %>% filter(LP_class == "LDL"), formula = Bin_value ~ Treatment)
summary(means_comp)
Anova(means_comp, type = "3")

drug_PAGE %>% 
  filter(LP_class == "LDL") %>%
  compare_means(formula = Bin_value~Treatment,
                ref.group = "Vehicle", 
                p.adjust.method = "bonf", 
                method = "t.test")
```

## Prochlorperazine

```{r}
#| label: LipoGlo assay with homogenization endpoint of Prochlorperazine treated samples
drug_name = "Prochlorperazine"

drug_hom <- read_plate("Figure_3_data/homogenate/Pcp_lpCount.csv", well_ids_column = "Wells")
drug_hom <- drug_hom %>%
  na.omit() %>%
  filter(Treatment != "Empty") %>%
  pivot_longer(cols = c("P1", "P2", "P3"), names_to = "Replicate", values_to = "RLU")

drug_hom_veh_summary <- drug_hom %>% 
  group_by(Replicate, Treatment) %>% 
  summarise(vehicle_mean = mean(RLU, na.rm = TRUE)) %>% 
  filter(Treatment == "Vehicle") %>%
  select(-Treatment)

drug_hom <- drug_hom %>% 
  full_join(drug_hom_veh_summary) %>%
  mutate(fold_change = RLU/vehicle_mean)

drug_hom <- drug_hom %>% 
  group_by(Treatment) %>%
  mutate(fold_change_Q1 = (quantile(fold_change, na.rm = TRUE)[[2]]), 
         fold_change_Q3 = (quantile(fold_change, na.rm = TRUE)[[4]]), 
         fold_change_IQR = IQR(fold_change, na.rm = TRUE),
         Outlier = if_else(fold_change > fold_change_Q3+1.5*(fold_change_IQR) |
                             fold_change < fold_change_Q1-1.5*(fold_change_IQR),
                           "Outlier", 
                           "Normal"))

y_lim <- drug_hom %>% 
  filter(Outlier == "Normal") %>%
  pull(fold_change) %>% 
  max(.) %>% 
  round_any(., 1, ceiling)

ggplot(data = drug_hom %>% 
  filter(Outlier == "Normal"),
       aes(x = factor(Treatment, levels = c("Vehicle",
                                            "5 µM lomitapide",
                                            "4 µM",
                                            "8 µM")), 
           y = fold_change)) +
  geom_boxplot(outlier.shape = NA) +
  geom_jitter(width = 0.2, shape = 1, alpha = 0.6) +
  theme_bw() +
  scale_y_continuous(
    labels = comma, limits = c(0,1.75), #c(lower limit, upper limit)
    breaks = seq(0,1.75,0.25)) +  #seq(lower limit, upper limit, tick frequency)
  labs(x = "Treatment",
       y = str_wrap("Fold chance in luminescense (relative to vehicle)", 
                    width = 30)) +
  theme(
    axis.text.x = element_text(face = 'bold', angle = 45, hjust = 1), 
    axis.title.x = element_text(face = 'bold'),
    axis.text.y = element_text(face = 'bold'), 
    axis.title.y = element_text(face = 'bold'),
    legend.title = element_text(face = 'bold'),
    legend.text = element_text(face = 'bold'),
    strip.text.x = element_text(face = 'bold'))
ggsave(paste0(drug_name, "_hom.svg"), width = 3, height = 4, path = "./output_data/")

drug_hom %>%
  filter(Outlier == "Normal") %>%
  group_by(Treatment) %>%
  summarise(n = n())

drug_hom %>% 
  filter(Outlier == "Normal") %>%
  compare_means(formula = fold_change~Treatment,
                ref.group = "Vehicle", 
                p.adjust.method = "bonf", 
                method = "t.test")

aov_drug_hom <- aov(data = drug_hom %>% filter(Outlier == "Normal"), 
                   formula = fold_change ~ Treatment)
summary(aov_drug_hom)

# nativePAGE
drug_PAGE <- read.csv(file = "Figure_3_data/nativePAGE/Pcp_PAGEquant.csv")

drug_PAGE <- drug_PAGE %>% 
  pivot_longer(cols = c("ZM", "VLDL", "IDL", "LDL"),
               names_to = "LP_class", 
               values_to = "Bin_value")

ggplot(data = drug_PAGE,
       aes(x = factor(Treatment, c("Vehicle",
                                   "5 µM lomitapide",
                                   "4 µM",
                                   "8 µM")),
           y = Bin_value*100)) +
  geom_boxplot(outlier.shape = NA) +
  geom_jitter(width = 0.2, shape = 1, alpha = 0.6) +
  theme_bw() +
  scale_y_continuous(
    labels = comma, limits = c(0,80), #c(lower limit, upper limit)
    breaks = seq(0,80,10)) +  #seq(lower limit, upper limit, tick frequency)
  labs(x = "Treatment",
       y = "Proportion of total lipoproteins (%)") +
  theme(
    axis.text.x = element_text(face = 'bold', angle = 45, hjust = 1), 
    axis.title.x = element_text(face = 'bold'),
    axis.text.y = element_text(face = 'bold'), 
    axis.title.y = element_text(face = 'bold'),
    legend.title = element_text(face = 'bold'),
    legend.text = element_text(face = 'bold'),
    strip.text.x = element_text(face = 'bold')) +
  facet_grid(~factor(LP_class, c("ZM", "VLDL", "IDL", "LDL")))
ggsave(paste0(drug_name, "_nPAGE.svg"), width = 5, height = 4, path = "./output_data/")

drug_PAGE %>%
  group_by(Treatment, LP_class) %>%
  summarise(n = n())

means_comp <- aov(data = drug_PAGE %>% filter(LP_class == "ZM"), formula = Bin_value ~ Treatment)
summary(means_comp)
Anova(means_comp, type = "3")

drug_PAGE %>% 
  filter(LP_class == "ZM") %>%
  compare_means(formula = Bin_value~Treatment,
                ref.group = "Vehicle", 
                p.adjust.method = "bonf", 
                method = "t.test")

means_comp <- aov(data = drug_PAGE %>% filter(LP_class == "VLDL"), formula = Bin_value ~ Treatment)
summary(means_comp)
Anova(means_comp, type = "3")

drug_PAGE %>% 
  filter(LP_class == "VLDL") %>%
  compare_means(formula = Bin_value~Treatment,
                ref.group = "Vehicle", 
                p.adjust.method = "bonf", 
                method = "t.test")

means_comp <- aov(data = drug_PAGE %>% filter(LP_class == "IDL"), formula = Bin_value ~ Treatment)
summary(means_comp)
Anova(means_comp, type = "3")

drug_PAGE %>% 
  filter(LP_class == "IDL") %>%
  compare_means(formula = Bin_value~Treatment,
                ref.group = "Vehicle", 
                p.adjust.method = "bonf", 
                method = "t.test")

means_comp <- aov(data = drug_PAGE %>% filter(LP_class == "LDL"), formula = Bin_value ~ Treatment)
summary(means_comp)
Anova(means_comp, type = "3")

drug_PAGE %>% 
  filter(LP_class == "LDL") %>%
  compare_means(formula = Bin_value~Treatment,
                ref.group = "Vehicle", 
                p.adjust.method = "bonf", 
                method = "t.test")
```

## Pimethixene

```{r}
#| label: LipoGlo assay with homogenization endpoint of Pimethixene treated samples
drug_name = "Pimethixene"

drug_hom <- read_plate("Figure_3_data/homogenate/Pmx_lpCount.csv", well_ids_column = "Wells")
drug_hom <- drug_hom %>%
  na.omit() %>%
  filter(Treatment != "Empty") %>%
  pivot_longer(cols = c("P1", "P2", "P3"), names_to = "Replicate", values_to = "RLU")

drug_hom_veh_summary <- drug_hom %>% 
  group_by(Replicate, Treatment) %>% 
  summarise(vehicle_mean = mean(RLU, na.rm = TRUE)) %>% 
  filter(Treatment == "Vehicle") %>%
  select(-Treatment)

drug_hom <- drug_hom %>% 
  full_join(drug_hom_veh_summary) %>%
  mutate(fold_change = RLU/vehicle_mean)

drug_hom <- drug_hom %>% 
  group_by(Treatment) %>%
  mutate(fold_change_Q1 = (quantile(fold_change, na.rm = TRUE)[[2]]), 
         fold_change_Q3 = (quantile(fold_change, na.rm = TRUE)[[4]]), 
         fold_change_IQR = IQR(fold_change, na.rm = TRUE),
         Outlier = if_else(fold_change > fold_change_Q3+1.5*(fold_change_IQR) |
                             fold_change < fold_change_Q1-1.5*(fold_change_IQR),
                           "Outlier", 
                           "Normal"))

y_lim <- drug_hom %>% 
  filter(Outlier == "Normal") %>%
  pull(fold_change) %>% 
  max(.) %>% 
  round_any(., 1, ceiling)

ggplot(data = drug_hom %>% 
  filter(Outlier == "Normal"),
       aes(x = factor(Treatment, levels = c("Vehicle",
                                            "5 µM lomitapide",
                                            "2 µM",
                                            "4 µM")), 
           y = fold_change)) +
  geom_boxplot(outlier.shape = NA) +
  geom_jitter(width = 0.2, shape = 1, alpha = 0.6) +
  theme_bw() +
  scale_y_continuous(
    labels = comma, limits = c(0,1.75), #c(lower limit, upper limit)
    breaks = seq(0,1.75,0.25)) +  #seq(lower limit, upper limit, tick frequency)
  labs(x = "Treatment",
       y = str_wrap("Fold chance in luminescense (relative to vehicle)", 
                    width = 30)) +
  theme(
    axis.text.x = element_text(face = 'bold', angle = 45, hjust = 1), 
    axis.title.x = element_text(face = 'bold'),
    axis.text.y = element_text(face = 'bold'), 
    axis.title.y = element_text(face = 'bold'),
    legend.title = element_text(face = 'bold'),
    legend.text = element_text(face = 'bold'),
    strip.text.x = element_text(face = 'bold'))
ggsave(paste0(drug_name, "_hom.svg"), width = 3, height = 4, path = "./output_data/")

drug_hom %>%
  filter(Outlier == "Normal") %>%
  group_by(Treatment) %>%
  summarise(n = n())

drug_hom %>% 
  filter(Outlier == "Normal") %>%
  compare_means(formula = fold_change~Treatment,
                ref.group = "Vehicle", 
                p.adjust.method = "bonf", 
                method = "t.test")

aov_drug_hom <- aov(data = drug_hom %>% filter(Outlier == "Normal"), 
                   formula = fold_change ~ Treatment)
summary(aov_drug_hom)

# nativePAGE
drug_PAGE <- read.csv(file = "Figure_3_data/nativePAGE/Pmx_PAGEquant.csv")

drug_PAGE <- drug_PAGE %>% 
  pivot_longer(cols = c("ZM", "VLDL", "IDL", "LDL"),
               names_to = "LP_class", 
               values_to = "Bin_value")

ggplot(data = drug_PAGE,
       aes(x = factor(Treatment, c("Vehicle",
                                   "5 µM lomitapide",
                                   "4 µM",
                                   "8 µM")),
           y = Bin_value*100)) +
  geom_boxplot(outlier.shape = NA) +
  geom_jitter(width = 0.2, shape = 1, alpha = 0.6) +
  theme_bw() +
  scale_y_continuous(
    labels = comma, limits = c(0,80), #c(lower limit, upper limit)
    breaks = seq(0,80,10)) +  #seq(lower limit, upper limit, tick frequency)
  labs(x = "Treatment",
       y = "Proportion of total lipoproteins (%)") +
  theme(
    axis.text.x = element_text(face = 'bold', angle = 45, hjust = 1), 
    axis.title.x = element_text(face = 'bold'),
    axis.text.y = element_text(face = 'bold'), 
    axis.title.y = element_text(face = 'bold'),
    legend.title = element_text(face = 'bold'),
    legend.text = element_text(face = 'bold'),
    strip.text.x = element_text(face = 'bold')) +
  facet_grid(~factor(LP_class, c("ZM", "VLDL", "IDL", "LDL")))
ggsave(paste0(drug_name, "_nPAGE.svg"), width = 5, height = 4, path = "./output_data/")

drug_PAGE %>%
  group_by(Treatment, LP_class) %>%
  summarise(n = n())

means_comp <- aov(data = drug_PAGE %>% filter(LP_class == "ZM"), formula = Bin_value ~ Treatment)
summary(means_comp)
Anova(means_comp, type = "3")

drug_PAGE %>% 
  filter(LP_class == "ZM") %>%
  compare_means(formula = Bin_value~Treatment,
                ref.group = "Vehicle", 
                p.adjust.method = "bonf", 
                method = "t.test")

means_comp <- aov(data = drug_PAGE %>% filter(LP_class == "VLDL"), formula = Bin_value ~ Treatment)
summary(means_comp)
Anova(means_comp, type = "3")

drug_PAGE %>% 
  filter(LP_class == "VLDL") %>%
  compare_means(formula = Bin_value~Treatment,
                ref.group = "Vehicle", 
                p.adjust.method = "bonf", 
                method = "t.test")

means_comp <- aov(data = drug_PAGE %>% filter(LP_class == "IDL"), formula = Bin_value ~ Treatment)
summary(means_comp)
Anova(means_comp, type = "3")

drug_PAGE %>% 
  filter(LP_class == "IDL") %>%
  compare_means(formula = Bin_value~Treatment,
                ref.group = "Vehicle", 
                p.adjust.method = "bonf", 
                method = "t.test")

means_comp <- aov(data = drug_PAGE %>% filter(LP_class == "LDL"), formula = Bin_value ~ Treatment)
summary(means_comp)
Anova(means_comp, type = "3")

drug_PAGE %>% 
  filter(LP_class == "LDL") %>%
  compare_means(formula = Bin_value~Treatment,
                ref.group = "Vehicle", 
                p.adjust.method = "bonf", 
                method = "t.test")
```

## Pomiferin

```{r}
#| label: LipoGlo assay with homogenization endpoint of Pomiferin treated samples
drug_name = "Pomiferin"

drug_hom <- read_plate("Figure_3_data/homogenate/Pom_lpCount.csv", well_ids_column = "Wells")
drug_hom <- drug_hom %>%
  na.omit() %>%
  filter(Treatment != "Empty") %>%
  pivot_longer(cols = c("P1", "P2"), names_to = "Replicate", values_to = "RLU")

drug_hom_veh_summary <- drug_hom %>% 
  group_by(Replicate, Treatment) %>% 
  summarise(vehicle_mean = mean(RLU, na.rm = TRUE)) %>% 
  filter(Treatment == "Vehicle") %>%
  select(-Treatment)

drug_hom <- drug_hom %>% 
  full_join(drug_hom_veh_summary) %>%
  mutate(fold_change = RLU/vehicle_mean)

drug_hom <- drug_hom %>% 
  group_by(Treatment) %>%
  mutate(fold_change_Q1 = (quantile(fold_change, na.rm = TRUE)[[2]]), 
         fold_change_Q3 = (quantile(fold_change, na.rm = TRUE)[[4]]), 
         fold_change_IQR = IQR(fold_change, na.rm = TRUE),
         Outlier = if_else(fold_change > fold_change_Q3+1.5*(fold_change_IQR) |
                             fold_change < fold_change_Q1-1.5*(fold_change_IQR),
                           "Outlier", 
                           "Normal"))

y_lim <- drug_hom %>% 
  filter(Outlier == "Normal") %>%
  pull(fold_change) %>% 
  max(.) %>% 
  round_any(., 1, ceiling)

ggplot(data = drug_hom %>% 
  filter(Outlier == "Normal"),
       aes(x = factor(Treatment, levels = c("Vehicle",
                                            "5 µM lomitapide",
                                            "2 µM",
                                            "4 µM")), 
           y = fold_change)) +
  geom_boxplot(outlier.shape = NA) +
  geom_jitter(width = 0.2, shape = 1, alpha = 0.6) +
  theme_bw() +
  scale_y_continuous(
    labels = comma, limits = c(0,1.75), #c(lower limit, upper limit)
    breaks = seq(0,1.75,0.25)) +  #seq(lower limit, upper limit, tick frequency)
  labs(x = "Treatment",
       y = str_wrap("Fold chance in luminescense (relative to vehicle)", 
                    width = 30)) +
  theme(
    axis.text.x = element_text(face = 'bold', angle = 45, hjust = 1), 
    axis.title.x = element_text(face = 'bold'),
    axis.text.y = element_text(face = 'bold'), 
    axis.title.y = element_text(face = 'bold'),
    legend.title = element_text(face = 'bold'),
    legend.text = element_text(face = 'bold'),
    strip.text.x = element_text(face = 'bold'))
ggsave(paste0(drug_name, "_hom.svg"), width = 3, height = 4, path = "./output_data/")

drug_hom %>%
  filter(Outlier == "Normal") %>%
  group_by(Treatment) %>%
  summarise(n = n())

drug_hom %>% 
  filter(Outlier == "Normal") %>%
  compare_means(formula = fold_change~Treatment,
                ref.group = "Vehicle", 
                p.adjust.method = "bonf", 
                method = "t.test")

aov_drug_hom <- aov(data = drug_hom %>% filter(Outlier == "Normal"), 
                   formula = fold_change ~ Treatment)
summary(aov_drug_hom)

# nativePAGE
drug_PAGE <- read.csv(file = "Figure_3_data/nativePAGE/Pom_PAGEquant.csv")

drug_PAGE <- drug_PAGE %>% 
  pivot_longer(cols = c("ZM", "VLDL", "IDL", "LDL"),
               names_to = "LP_class", 
               values_to = "Bin_value")

ggplot(data = drug_PAGE,
       aes(x = factor(Treatment, c("Vehicle",
                                   "5 µM lomitapide",
                                   "4 µM",
                                   "8 µM")),
           y = Bin_value*100)) +
  geom_boxplot(outlier.shape = NA) +
  geom_jitter(width = 0.2, shape = 1, alpha = 0.6) +
  theme_bw() +
  scale_y_continuous(
    labels = comma, limits = c(0,80), #c(lower limit, upper limit)
    breaks = seq(0,80,10)) +  #seq(lower limit, upper limit, tick frequency)
  labs(x = "Treatment",
       y = "Proportion of total lipoproteins (%)") +
  theme(
    axis.text.x = element_text(face = 'bold', angle = 45, hjust = 1), 
    axis.title.x = element_text(face = 'bold'),
    axis.text.y = element_text(face = 'bold'), 
    axis.title.y = element_text(face = 'bold'),
    legend.title = element_text(face = 'bold'),
    legend.text = element_text(face = 'bold'),
    strip.text.x = element_text(face = 'bold')) +
  facet_grid(~factor(LP_class, c("ZM", "VLDL", "IDL", "LDL")))
ggsave(paste0(drug_name, "_nPAGE.svg"), width = 5, height = 4, path = "./output_data/")

drug_PAGE %>%
  group_by(Treatment, LP_class) %>%
  summarise(n = n())

means_comp <- aov(data = drug_PAGE %>% filter(LP_class == "ZM"), formula = Bin_value ~ Treatment)
summary(means_comp)
Anova(means_comp, type = "3")

drug_PAGE %>% 
  filter(LP_class == "ZM") %>%
  compare_means(formula = Bin_value~Treatment,
                ref.group = "Vehicle", 
                p.adjust.method = "bonf", 
                method = "t.test")

means_comp <- aov(data = drug_PAGE %>% filter(LP_class == "VLDL"), formula = Bin_value ~ Treatment)
summary(means_comp)
Anova(means_comp, type = "3")

drug_PAGE %>% 
  filter(LP_class == "VLDL") %>%
  compare_means(formula = Bin_value~Treatment,
                ref.group = "Vehicle", 
                p.adjust.method = "bonf", 
                method = "t.test")

means_comp <- aov(data = drug_PAGE %>% filter(LP_class == "IDL"), formula = Bin_value ~ Treatment)
summary(means_comp)
Anova(means_comp, type = "3")

drug_PAGE %>% 
  filter(LP_class == "IDL") %>%
  compare_means(formula = Bin_value~Treatment,
                ref.group = "Vehicle", 
                p.adjust.method = "bonf", 
                method = "t.test")

means_comp <- aov(data = drug_PAGE %>% filter(LP_class == "LDL"), formula = Bin_value ~ Treatment)
summary(means_comp)
Anova(means_comp, type = "3")

drug_PAGE %>% 
  filter(LP_class == "LDL") %>%
  compare_means(formula = Bin_value~Treatment,
                ref.group = "Vehicle", 
                p.adjust.method = "bonf", 
                method = "t.test")
```

## Riboflavin tetrabutyrate

```{r}
#| label: LipoGlo assay with homogenization endpoint of Riboflavin tetrabutyrate treated samples
drug_name = "Riboflavin_tetrabutyrate"

drug_hom <- read_plate("Figure_3_data/homogenate/Rft_lpCount.csv", well_ids_column = "Wells")
drug_hom <- drug_hom %>%
  na.omit() %>%
  filter(Treatment != "Empty") %>%
  pivot_longer(cols = c("P1", "P2", "P3"), names_to = "Replicate", values_to = "RLU")

drug_hom_veh_summary <- drug_hom %>% 
  group_by(Replicate, Treatment) %>% 
  summarise(vehicle_mean = mean(RLU, na.rm = TRUE)) %>% 
  filter(Treatment == "Vehicle") %>%
  select(-Treatment)

drug_hom <- drug_hom %>% 
  full_join(drug_hom_veh_summary) %>%
  mutate(fold_change = RLU/vehicle_mean)

drug_hom <- drug_hom %>% 
  group_by(Treatment) %>%
  mutate(fold_change_Q1 = (quantile(fold_change, na.rm = TRUE)[[2]]), 
         fold_change_Q3 = (quantile(fold_change, na.rm = TRUE)[[4]]), 
         fold_change_IQR = IQR(fold_change, na.rm = TRUE),
         Outlier = if_else(fold_change > fold_change_Q3+1.5*(fold_change_IQR) |
                             fold_change < fold_change_Q1-1.5*(fold_change_IQR),
                           "Outlier", 
                           "Normal"))

y_lim <- drug_hom %>% 
  filter(Outlier == "Normal") %>%
  pull(fold_change) %>% 
  max(.) %>% 
  round_any(., 1, ceiling)

ggplot(data = drug_hom %>% 
  filter(Outlier == "Normal"),
       aes(x = factor(Treatment, levels = c("Vehicle",
                                            "5 µM lomitapide",
                                            "4 µM",
                                            "8 µM")), 
           y = fold_change)) +
  geom_boxplot(outlier.shape = NA) +
  geom_jitter(width = 0.2, shape = 1, alpha = 0.6) +
  theme_bw() +
  scale_y_continuous(
    labels = comma, limits = c(0,1.75), #c(lower limit, upper limit)
    breaks = seq(0,1.75,0.25)) +  #seq(lower limit, upper limit, tick frequency)
  labs(x = "Treatment",
       y = str_wrap("Fold chance in luminescense (relative to vehicle)", 
                    width = 30)) +
  theme(
    axis.text.x = element_text(face = 'bold', angle = 45, hjust = 1), 
    axis.title.x = element_text(face = 'bold'),
    axis.text.y = element_text(face = 'bold'), 
    axis.title.y = element_text(face = 'bold'),
    legend.title = element_text(face = 'bold'),
    legend.text = element_text(face = 'bold'),
    strip.text.x = element_text(face = 'bold'))
ggsave(paste0(drug_name, "_hom.svg"), width = 3, height = 4, path = "./output_data/")

drug_hom %>%
  filter(Outlier == "Normal") %>%
  group_by(Treatment) %>%
  summarise(n = n())

drug_hom %>% 
  filter(Outlier == "Normal") %>%
  compare_means(formula = fold_change~Treatment,
                ref.group = "Vehicle", 
                p.adjust.method = "bonf", 
                method = "t.test")

aov_drug_hom <- aov(data = drug_hom %>% filter(Outlier == "Normal"), 
                   formula = fold_change ~ Treatment)
summary(aov_drug_hom)

# nativePAGE
drug_PAGE <- read.csv(file = "Figure_3_data/nativePAGE/Rft_PAGEquant.csv")

drug_PAGE <- drug_PAGE %>% 
  pivot_longer(cols = c("ZM", "VLDL", "IDL", "LDL"),
               names_to = "LP_class", 
               values_to = "Bin_value")

ggplot(data = drug_PAGE,
       aes(x = factor(Treatment, c("Vehicle",
                                   "5 µM lomitapide",
                                   "4 µM",
                                   "8 µM")),
           y = Bin_value*100)) +
  geom_boxplot(outlier.shape = NA) +
  geom_jitter(width = 0.2, shape = 1, alpha = 0.6) +
  theme_bw() +
  scale_y_continuous(
    labels = comma, limits = c(0,80), #c(lower limit, upper limit)
    breaks = seq(0,80,10)) +  #seq(lower limit, upper limit, tick frequency)
  labs(x = "Treatment",
       y = "Proportion of total lipoproteins (%)") +
  theme(
    axis.text.x = element_text(face = 'bold', angle = 45, hjust = 1), 
    axis.title.x = element_text(face = 'bold'),
    axis.text.y = element_text(face = 'bold'), 
    axis.title.y = element_text(face = 'bold'),
    legend.title = element_text(face = 'bold'),
    legend.text = element_text(face = 'bold'),
    strip.text.x = element_text(face = 'bold')) +
  facet_grid(~factor(LP_class, c("ZM", "VLDL", "IDL", "LDL")))
ggsave(paste0(drug_name, "_nPAGE.svg"), width = 5, height = 4, path = "./output_data/")

drug_PAGE %>%
  group_by(Treatment, LP_class) %>%
  summarise(n = n())

means_comp <- aov(data = drug_PAGE %>% filter(LP_class == "ZM"), formula = Bin_value ~ Treatment)
summary(means_comp)
Anova(means_comp, type = "3")

drug_PAGE %>% 
  filter(LP_class == "ZM") %>%
  compare_means(formula = Bin_value~Treatment,
                ref.group = "Vehicle", 
                p.adjust.method = "bonf", 
                method = "t.test")

means_comp <- aov(data = drug_PAGE %>% filter(LP_class == "VLDL"), formula = Bin_value ~ Treatment)
summary(means_comp)
Anova(means_comp, type = "3")

drug_PAGE %>% 
  filter(LP_class == "VLDL") %>%
  compare_means(formula = Bin_value~Treatment,
                ref.group = "Vehicle", 
                p.adjust.method = "bonf", 
                method = "t.test")

means_comp <- aov(data = drug_PAGE %>% filter(LP_class == "IDL"), formula = Bin_value ~ Treatment)
summary(means_comp)
Anova(means_comp, type = "3")

drug_PAGE %>% 
  filter(LP_class == "IDL") %>%
  compare_means(formula = Bin_value~Treatment,
                ref.group = "Vehicle", 
                p.adjust.method = "bonf", 
                method = "t.test")

means_comp <- aov(data = drug_PAGE %>% filter(LP_class == "LDL"), formula = Bin_value ~ Treatment)
summary(means_comp)
Anova(means_comp, type = "3")

drug_PAGE %>% 
  filter(LP_class == "LDL") %>%
  compare_means(formula = Bin_value~Treatment,
                ref.group = "Vehicle", 
                p.adjust.method = "bonf", 
                method = "t.test")
```

## Thiethylperazine

```{r}
#| label: LipoGlo assay with homogenization endpoint of Thiethylperazine treated samples
drug_name = "Thiethylperazine"

drug_hom <- read_plate("Figure_3_data/homogenate/Tpz_lpCount.csv", well_ids_column = "Wells")
drug_hom <- drug_hom %>%
  na.omit() %>%
  filter(Treatment != "Empty") %>%
  pivot_longer(cols = c("P1", "P2", "P3"), names_to = "Replicate", values_to = "RLU")

drug_hom_veh_summary <- drug_hom %>% 
  group_by(Replicate, Treatment) %>% 
  summarise(vehicle_mean = mean(RLU, na.rm = TRUE)) %>% 
  filter(Treatment == "Vehicle") %>%
  select(-Treatment)

drug_hom <- drug_hom %>% 
  full_join(drug_hom_veh_summary) %>%
  mutate(fold_change = RLU/vehicle_mean)

drug_hom <- drug_hom %>% 
  group_by(Treatment) %>%
  mutate(fold_change_Q1 = (quantile(fold_change, na.rm = TRUE)[[2]]), 
         fold_change_Q3 = (quantile(fold_change, na.rm = TRUE)[[4]]), 
         fold_change_IQR = IQR(fold_change, na.rm = TRUE),
         Outlier = if_else(fold_change > fold_change_Q3+1.5*(fold_change_IQR) |
                             fold_change < fold_change_Q1-1.5*(fold_change_IQR),
                           "Outlier", 
                           "Normal"))

y_lim <- drug_hom %>% 
  filter(Outlier == "Normal") %>%
  pull(fold_change) %>% 
  max(.) %>% 
  round_any(., 1, ceiling)

ggplot(data = drug_hom %>% 
  filter(Outlier == "Normal"),
       aes(x = factor(Treatment, levels = c("Vehicle",
                                            "5 µM lomitapide",
                                            "2 µM",
                                            "4 µM")), 
           y = fold_change)) +
  geom_boxplot(outlier.shape = NA) +
  geom_jitter(width = 0.2, shape = 1, alpha = 0.6) +
  theme_bw() +
  scale_y_continuous(
    labels = comma, limits = c(0,1.75), #c(lower limit, upper limit)
    breaks = seq(0,1.75,0.25)) +  #seq(lower limit, upper limit, tick frequency)
  labs(x = "Treatment",
       y = str_wrap("Fold chance in luminescense (relative to vehicle)", 
                    width = 30)) +
  theme(
    axis.text.x = element_text(face = 'bold', angle = 45, hjust = 1), 
    axis.title.x = element_text(face = 'bold'),
    axis.text.y = element_text(face = 'bold'), 
    axis.title.y = element_text(face = 'bold'),
    legend.title = element_text(face = 'bold'),
    legend.text = element_text(face = 'bold'),
    strip.text.x = element_text(face = 'bold'))
ggsave(paste0(drug_name, "_hom.svg"), width = 3, height = 4, path = "./output_data/")

drug_hom %>%
  filter(Outlier == "Normal") %>%
  group_by(Treatment) %>%
  summarise(n = n())

drug_hom %>% 
  filter(Outlier == "Normal") %>%
  compare_means(formula = fold_change~Treatment,
                ref.group = "Vehicle", 
                p.adjust.method = "bonf", 
                method = "t.test")

aov_drug_hom <- aov(data = drug_hom %>% filter(Outlier == "Normal"), 
                   formula = fold_change ~ Treatment)
summary(aov_drug_hom)

# nativePAGE
drug_PAGE <- read.csv(file = "Figure_3_data/nativePAGE/Tpz_PAGEquant.csv")

drug_PAGE <- drug_PAGE %>% 
  pivot_longer(cols = c("ZM", "VLDL", "IDL", "LDL"),
               names_to = "LP_class", 
               values_to = "Bin_value")

ggplot(data = drug_PAGE,
       aes(x = factor(Treatment, c("Vehicle",
                                   "5 µM lomitapide",
                                   "4 µM",
                                   "8 µM")),
           y = Bin_value*100)) +
  geom_boxplot(outlier.shape = NA) +
  geom_jitter(width = 0.2, shape = 1, alpha = 0.6) +
  theme_bw() +
  scale_y_continuous(
    labels = comma, limits = c(0,80), #c(lower limit, upper limit)
    breaks = seq(0,80,10)) +  #seq(lower limit, upper limit, tick frequency)
  labs(x = "Treatment",
       y = "Proportion of total lipoproteins (%)") +
  theme(
    axis.text.x = element_text(face = 'bold', angle = 45, hjust = 1), 
    axis.title.x = element_text(face = 'bold'),
    axis.text.y = element_text(face = 'bold'), 
    axis.title.y = element_text(face = 'bold'),
    legend.title = element_text(face = 'bold'),
    legend.text = element_text(face = 'bold'),
    strip.text.x = element_text(face = 'bold')) +
  facet_grid(~factor(LP_class, c("ZM", "VLDL", "IDL", "LDL")))
ggsave(paste0(drug_name, "_nPAGE.svg"), width = 5, height = 4, path = "./output_data/")

drug_PAGE %>%
  group_by(Treatment, LP_class) %>%
  summarise(n = n())

means_comp <- aov(data = drug_PAGE %>% filter(LP_class == "ZM"), formula = Bin_value ~ Treatment)
summary(means_comp)
Anova(means_comp, type = "3")

drug_PAGE %>% 
  filter(LP_class == "ZM") %>%
  compare_means(formula = Bin_value~Treatment,
                ref.group = "Vehicle", 
                p.adjust.method = "bonf", 
                method = "t.test")

means_comp <- aov(data = drug_PAGE %>% filter(LP_class == "VLDL"), formula = Bin_value ~ Treatment)
summary(means_comp)
Anova(means_comp, type = "3")

drug_PAGE %>% 
  filter(LP_class == "VLDL") %>%
  compare_means(formula = Bin_value~Treatment,
                ref.group = "Vehicle", 
                p.adjust.method = "bonf", 
                method = "t.test")

means_comp <- aov(data = drug_PAGE %>% filter(LP_class == "IDL"), formula = Bin_value ~ Treatment)
summary(means_comp)
Anova(means_comp, type = "3")

drug_PAGE %>% 
  filter(LP_class == "IDL") %>%
  compare_means(formula = Bin_value~Treatment,
                ref.group = "Vehicle", 
                p.adjust.method = "bonf", 
                method = "t.test")

means_comp <- aov(data = drug_PAGE %>% filter(LP_class == "LDL"), formula = Bin_value ~ Treatment)
summary(means_comp)
Anova(means_comp, type = "3")

drug_PAGE %>% 
  filter(LP_class == "LDL") %>%
  compare_means(formula = Bin_value~Treatment,
                ref.group = "Vehicle", 
                p.adjust.method = "bonf", 
                method = "t.test")
```

# Figure 4 - Enoxolone

```{r}
#| label: LipoGlo assay with fixation endpoint of enoxolone treated samples
exl_fix <- read_plate("Figure_4_data/Enoxolone_fix_data_summary.csv", well_ids_column = "Wells")
exl_fix <- exl_fix %>%
  pivot_longer(cols = c(3:ncol(.)), names_to = "Experiment", values_to = "RLU") %>%
  filter(Treatment != "Empty") 

exl_fix_veh_summary <- exl_fix %>% 
  group_by(Experiment, Treatment) %>% 
  summarise(vehicle_mean = mean(RLU)) %>% 
  filter(Treatment == "Vehicle") %>%
  select(-Treatment)

exl_fix <- exl_fix %>% 
  full_join(exl_fix_veh_summary) %>%
  mutate(fold_change = RLU/vehicle_mean)

exl_fix <- exl_fix %>% 
  group_by(Treatment) %>%
  mutate(fold_change_Q1 = (quantile(fold_change)[[2]]), 
         fold_change_Q3 = (quantile(fold_change)[[4]]), 
         fold_change_IQR = IQR(fold_change),
         Outlier = if_else(fold_change > fold_change_Q3+1.5*(fold_change_IQR) |
                             fold_change < fold_change_Q1-1.5*(fold_change_IQR),
                           "Outlier", 
                           "Normal"))

y_lim <- exl_fix %>% 
  filter(Outlier == "Normal") %>%
  pull(fold_change) %>% 
  max(.) %>% 
  round_any(., 1, ceiling)

ggplot(data = exl_fix %>% 
  filter(Outlier == "Normal"),
       aes(x = factor(Treatment, levels = c("Vehicle",
                                            "5 µM lomitapide",
                                            "0.0625 µM",
                                            "0.125 µM",
                                            "0.25 µM",
                                            "0.5 µM",
                                            "1 µM",
                                            "2 µM",
                                            "4 µM",
                                            "8 µM")), 
           y = fold_change)) +
  geom_boxplot(outlier.shape = NA) +
  geom_jitter(width = 0.2, shape = 1, alpha = 0.6) +
  theme_bw() +
  scale_y_continuous(
    labels = comma, limits = c(0,1.75), #c(lower limit, upper limit)
    breaks = seq(0,1.75,0.25)) +  #seq(lower limit, upper limit, tick frequency)
  labs(x = "Treatment",
       y = str_wrap("Fold chance in luminescense (relative to vehicle)", 
                    width = 30)) +
  theme(
    axis.text.x = element_text(face = 'bold', angle = 45, hjust = 1), 
    axis.title.x = element_text(face = 'bold'),
    axis.text.y = element_text(face = 'bold'), 
    axis.title.y = element_text(face = 'bold'),
    legend.title = element_text(face = 'bold'),
    legend.text = element_text(face = 'bold'),
    strip.text.x = element_text(face = 'bold'))
ggsave("Enoxolone_fix.pdf", width = 4, height = 4, path = "./output_data/")

exl_fix %>%
  filter(Outlier == "Normal") %>%
  group_by(Treatment) %>%
  summarise(n = n())

exl_fix %>% 
  filter(Outlier == "Normal") %>%
  compare_means(formula = fold_change~Treatment,
                ref.group = "Vehicle", 
                p.adjust.method = "bonf", 
                method = "t.test")

aov_exl_fix <- aov(data = exl_fix %>% filter(Outlier == "Normal"), 
                   formula = fold_change ~ Treatment)
summary(aov_exl_fix)
```

```{r}
#| label: LipoGlo assay with homogenization endpoint of enoxolone treated samples
exl_hom <- read_plate("Figure_4_data/Enoxolone_hom_data_summary.csv", well_ids_column = "Wells")
exl_hom <- exl_hom %>%
  pivot_longer(cols = c(3:ncol(.)), names_to = "Experiment", values_to = "RLU") %>%
  filter(Treatment != "Empty") 

exl_hom_veh_summary <- exl_hom %>% 
  group_by(Experiment, Treatment) %>% 
  summarise(vehicle_mean = mean(RLU)) %>% 
  filter(Treatment == "Vehicle") %>%
  select(-Treatment)

exl_hom <- exl_hom %>% 
  full_join(exl_hom_veh_summary) %>%
  mutate(fold_change = RLU/vehicle_mean)

exl_hom <- exl_hom %>% 
  group_by(Treatment) %>%
  mutate(fold_change_Q1 = (quantile(fold_change)[[2]]), 
         fold_change_Q3 = (quantile(fold_change)[[4]]), 
         fold_change_IQR = IQR(fold_change),
         Outlier = if_else(fold_change > fold_change_Q3+1.5*(fold_change_IQR) |
                             fold_change < fold_change_Q1-1.5*(fold_change_IQR),
                           "Outlier", 
                           "Normal"))

y_lim <- exl_hom %>% 
  filter(Outlier == "Normal") %>%
  pull(fold_change) %>% 
  max(.) %>% 
  round_any(., 1, ceiling)

ggplot(data = exl_hom %>%
         filter(Outlier == "Normal"),
       aes(x = factor(Treatment, levels = c("Vehicle",
                                            "5 µM lomitapide",
                                            "0.0625 µM",
                                            "0.125 µM",
                                            "0.25 µM",
                                            "0.5 µM",
                                            "1 µM",
                                            "2 µM",
                                            "4 µM",
                                            "8 µM")), 
           y = fold_change)) +
  geom_boxplot(outlier.shape = NA) +
  geom_jitter(width = 0.2, shape = 1, alpha = 0.6) +
  theme_bw() +
  scale_y_continuous(
    labels = comma, limits = c(0,1.75), #c(lower limit, upper limit)
    breaks = seq(0,1.75,0.25)) +  #seq(lower limit, upper limit, tick frequency)
  labs(x = "Treatment",
       y = str_wrap("Fold chance in luminescense (relative to vehicle)", 
                    width = 30)) +
  theme(
    axis.text.x = element_text(face = 'bold', angle = 45, hjust = 1), 
    axis.title.x = element_text(face = 'bold'),
    axis.text.y = element_text(face = 'bold'), 
    axis.title.y = element_text(face = 'bold'),
    legend.title = element_text(face = 'bold'),
    legend.text = element_text(face = 'bold'),
    strip.text.x = element_text(face = 'bold'))

ggsave("Enoxolone_hom.pdf", width = 4, height = 4, path = "./output_data/")

exl_hom %>%
  filter(Outlier == "Normal") %>%
  group_by(Treatment) %>%
  summarise(n = n())

exl_hom %>% 
  filter(Outlier == "Normal") %>%
  compare_means(formula = fold_change~Treatment,
                ref.group = "Vehicle", 
                p.adjust.method = "bonf", 
                method = "t.test")

aov_clutch1 <- aov(data = exl_hom %>% filter(Outlier == "Normal"), 
                   formula = fold_change ~ Treatment)
summary(aov_clutch1)
Anova(aov_clutch1, type = "3")
```

```{r}
#| label: NanoLuc activity analysis of enoxolone treated samples
exl_nl <- read_plate("Figure_4_data/Enoxolone_NL_activity_data_summary.csv", well_ids_column = "Wells")
exl_nl <- exl_nl %>%
  pivot_longer(cols = c(3:ncol(.)), names_to = "Experiment", values_to = "RLU") %>%
  filter(Treatment != "Empty") 

exl_nl_veh_summary <- exl_nl %>% 
  group_by(Experiment, Treatment) %>% 
  summarise(vehicle_mean = mean(RLU)) %>% 
  filter(Treatment == "Vehicle") %>%
  select(-Treatment)

exl_nl <- exl_nl %>% 
  full_join(exl_nl_veh_summary) %>%
  mutate(fold_change = RLU/vehicle_mean)

exl_nl <- exl_nl %>% 
  group_by(Treatment) %>%
  mutate(fold_change_Q1 = (quantile(fold_change)[[2]]), 
         fold_change_Q3 = (quantile(fold_change)[[4]]), 
         fold_change_IQR = IQR(fold_change),
         Outlier = if_else(fold_change > fold_change_Q3+1.5*(fold_change_IQR) |
                             fold_change < fold_change_Q1-1.5*(fold_change_IQR),
                           "Outlier", 
                           "Normal"))

y_lim <- exl_nl %>% 
  filter(Outlier == "Normal") %>%
  pull(fold_change) %>% 
  max(.) %>% 
  round_any(., 0.25, ceiling)

ggplot(data = exl_nl %>%
         filter(Outlier == "Normal"),
       aes(x = factor(Treatment, levels = c("Vehicle",
                                            "400 nM NanoLuc inhibitor",
                                            "0.0625 µM",
                                            "0.125 µM",
                                            "0.25 µM",
                                            "0.5 µM",
                                            "1 µM",
                                            "2 µM",
                                            "4 µM",
                                            "8 µM")), 
           y = fold_change)) +
  geom_boxplot(outlier.shape = NA) +
  geom_jitter(width = 0.2, shape = 1, alpha = 0.6) +
  theme_bw() +
  scale_y_continuous(
    labels = comma, limits = c(0,1.75), #c(lower limit, upper limit)
    breaks = seq(0,1.75,0.25)) +  #seq(lower limit, upper limit, tick frequency)
  labs(x = "Treatment",
       y = str_wrap("Fold chance in luminescense (relative to vehicle)", 
                    width = 30)) +
  theme(
    axis.text.x = element_text(face = 'bold', angle = 45, hjust = 1), 
    axis.title.x = element_text(face = 'bold'),
    axis.text.y = element_text(face = 'bold'), 
    axis.title.y = element_text(face = 'bold'),
    legend.title = element_text(face = 'bold'),
    legend.text = element_text(face = 'bold'),
    strip.text.x = element_text(face = 'bold')) + 
  scale_x_discrete(labels = function(x) str_wrap(x, width = 15))
ggsave("Enoxolone_NL_activity.pdf", width = 4, height = 4, path = "./output_data/")

exl_nl %>%
  filter(Outlier == "Normal") %>%
  group_by(Treatment) %>%
  summarise(n = n())

exl_nl %>% 
  filter(Outlier == "Normal") %>%
  compare_means(formula = fold_change~Treatment,
                ref.group = "Vehicle", 
                p.adjust.method = "bonf", 
                method = "t.test")

aov_clutch1 <- aov(data = exl_nl %>% filter(Outlier == "Normal"), 
                   formula = fold_change ~ Treatment)
summary(aov_clutch1)
Anova(aov_clutch1, type = "3")
```

```{r}
#| label: Standard length measurements of enoxlone treated animals

exl_sl <- read.csv(file = "Figure_4_data/enoxolone_SL_summary.csv")

exl_sl <- exl_sl %>%
  pivot_longer(cols = !X, names_to = "Treatment", values_to = "sl") %>%
  mutate(Treatment = str_replace(Treatment, "X5.µM.lomitapide", "5 µM lomitapide"),
         Treatment = str_replace(Treatment, "X8.µM.enoxolone", "8 µM enoxolone"),
         Treatment = str_replace(Treatment, "X4.µM.enoxolone", "4 µM enoxolone")) %>%
  na.omit() %>%
  group_by(Treatment) %>%
  mutate(sl_Q1 = (quantile(sl)[[2]]), 
         sl_Q3 = (quantile(sl)[[4]]), 
         sl_IQR = IQR(sl),
         Outlier = if_else(sl > sl_Q3+1.5*(sl_IQR) |
                             sl < sl_Q1-1.5*(sl_IQR),
                           "Outlier", 
                           "Normal"))

y_lim <- exl_sl %>% 
  filter(Outlier == "Normal") %>%
  pull(sl) %>% 
  max(.) %>% 
  round_any(., 1, ceiling)

ggplot(data = exl_sl %>%
         filter(Outlier == "Normal"),
       aes(x = factor(Treatment, levels = c("Vehicle",
                                            "5 µM lomitapide",
                                            "8 µM enoxolone",
                                            "4 µM enoxolone")), 
           y = sl)) +
  geom_boxplot(outlier.shape = NA) +
  geom_jitter(width = 0.2, shape = 1, alpha = 0.6) +
  theme_bw() +
  scale_y_continuous(
    labels = comma, limits = c(0,4.5), #c(lower limit, upper limit)
    breaks = seq(0,4.5,0.5)) +  #seq(lower limit, upper limit, tick frequency)
  labs(x = "Treatment",
       y = str_wrap("Standard Length (mm)")) +
  theme(
    axis.text.x = element_text(face = 'bold', angle = 45, hjust = 1), 
    axis.title.x = element_text(face = 'bold'),
    axis.text.y = element_text(face = 'bold'), 
    axis.title.y = element_text(face = 'bold'),
    legend.title = element_text(face = 'bold'),
    legend.text = element_text(face = 'bold'),
    strip.text.x = element_text(face = 'bold'))
ggsave("Enoxolone_SL.pdf", width = 3, height = 4, path = "./output_data/")

exl_sl %>%
  filter(Outlier == "Normal") %>%
  group_by(Treatment) %>%
  summarise(mean = mean(sl), sd = sd(sl), n = n())

exl_sl %>% 
  filter(Outlier == "Normal") %>%
  compare_means(formula = sl~Treatment,
                ref.group = "Vehicle", 
                p.adjust.method = "bonf", 
                method = "t.test")

aov_result <- aov(data = exl_sl %>% 
                    filter(Outlier == "Normal"), 
                  formula = sl ~ Treatment)
summary(aov_result)
Anova(aov_result, type = "3")
```

```{r}
#| label: Native PAGE analysis of enoxolone treated samples
EXL_PAGE <- read.csv(file = "Figure_4_data/Enoxolone_nPAGE_data_summary.csv")

EXL_PAGE <- EXL_PAGE %>% 
  pivot_longer(cols = c("ZM", "VLDL", "IDL", "LDL"),
               names_to = "LP_class", 
               values_to = "Bin_value")

EXL_PAGE <- EXL_PAGE %>% 
  group_by(Experiment, Treatment, Replicate) %>%
  summarize(Total = sum(Bin_value)) %>%
  full_join(EXL_PAGE) %>%
  mutate(Proportion = (Bin_value/Total)*100)

ggplot(data = EXL_PAGE,
       aes(x = factor(Treatment, c("Vehicle", "5 µM lomitapide", "8 µM enoxolone")),
           y = Proportion)) +
  geom_boxplot(outlier.shape = NA) +
  geom_jitter(width = 0.2, shape = 1, alpha = 0.6) +
  theme_bw() +
  scale_y_continuous(
    labels = comma, limits = c(0,80), #c(lower limit, upper limit)
    breaks = seq(0,80,10)) +  #seq(lower limit, upper limit, tick frequency)
  labs(x = "Treatment",
       y = "Proportion of total lipoproteins (%)") +
  theme(
    axis.text.x = element_text(face = 'bold', angle = 45, hjust = 1), 
    axis.title.x = element_text(face = 'bold'),
    axis.text.y = element_text(face = 'bold'), 
    axis.title.y = element_text(face = 'bold'),
    legend.title = element_text(face = 'bold'),
    legend.text = element_text(face = 'bold'),
    strip.text.x = element_text(face = 'bold')) +
  facet_grid(~factor(LP_class, c("ZM", "VLDL", "IDL", "LDL")))
ggsave("Enoxolone_nPAGE.pdf", width = 5, height = 4, path = "./output_data/")

EXL_PAGE %>%
  group_by(Treatment, LP_class) %>%
  summarise(n = n())

means_comp <- aov(data = EXL_PAGE %>% filter(LP_class == "ZM"), formula = Proportion ~ Treatment)
summary(means_comp)
Anova(means_comp, type = "3")

EXL_PAGE %>% 
  filter(LP_class == "ZM") %>%
  compare_means(formula = Proportion~Treatment,
                ref.group = "Vehicle", 
                p.adjust.method = "bonf", 
                method = "t.test")

means_comp <- aov(data = EXL_PAGE %>% filter(LP_class == "VLDL"), formula = Proportion ~ Treatment)
summary(means_comp)
Anova(means_comp, type = "3")

EXL_PAGE %>% 
  filter(LP_class == "VLDL") %>%
  compare_means(formula = Proportion~Treatment,
                ref.group = "Vehicle", 
                p.adjust.method = "bonf", 
                method = "t.test")

means_comp <- aov(data = EXL_PAGE %>% filter(LP_class == "IDL"), formula = Proportion ~ Treatment)
summary(means_comp)
Anova(means_comp, type = "3")

EXL_PAGE %>% 
  filter(LP_class == "IDL") %>%
  compare_means(formula = Proportion~Treatment,
                ref.group = "Vehicle", 
                p.adjust.method = "bonf", 
                method = "t.test")

means_comp <- aov(data = EXL_PAGE %>% filter(LP_class == "LDL"), formula = Proportion ~ Treatment)
summary(means_comp)
Anova(means_comp, type = "3")

EXL_PAGE %>% 
  filter(LP_class == "LDL") %>%
  compare_means(formula = Proportion~Treatment,
                ref.group = "Vehicle", 
                p.adjust.method = "bonf", 
                method = "t.test")
```

# Figure 5 - BIM5078 and BI6015

```{r}
#| label: BIM5078 and BI6015 effect on ApoB levels (fixation endpoint)
BB_fix <- read_plate("Figure_5_data/BIBIM_fix_summary_data.csv", well_ids_column = "Wells")
BB_fix <- BB_fix %>%
  pivot_longer(cols = c(3:ncol(.)), names_to = "Experiment", values_to = "RLU") %>%
  filter(Treatment != "Empty") %>%
  separate(col = "Experiment", into = c("Experiment", "Drug"), sep = "-")

BB_fix_veh_summary <- BB_fix %>% 
  group_by(Experiment, Treatment, Drug) %>% 
  summarise(vehicle_mean = mean(RLU)) %>% 
  filter(Treatment == "Vehicle") %>%
  ungroup() %>%
  select(-Treatment)

BB_fix <- BB_fix %>% 
  full_join(BB_fix_veh_summary, join_by(Drug, Experiment)) %>%
  mutate(fold_change = RLU/vehicle_mean)

BB_fix <- BB_fix %>% 
  group_by(Treatment, Drug) %>%
  mutate(fold_change_Q1 = (quantile(fold_change)[[2]]), 
         fold_change_Q3 = (quantile(fold_change)[[4]]), 
         fold_change_IQR = IQR(fold_change),
         Outlier = if_else(fold_change > fold_change_Q3+1.5*(fold_change_IQR) |
                             fold_change < fold_change_Q1-1.5*(fold_change_IQR),
                           "Outlier", 
                           "Normal"))

y_lim <- BB_fix %>% 
  filter(Outlier == "Normal") %>%
  pull(fold_change) %>% 
  max(.) %>% 
  round_any(., 1, ceiling)

ggplot(data = BB_fix %>% 
  filter(Outlier == "Normal"),
       aes(x = factor(Treatment, levels = c("Vehicle",
                                            "5 µM lomitapide",
                                            "0.0625 µM",
                                            "0.125 µM",
                                            "0.25 µM",
                                            "0.5 µM",
                                            "1 µM",
                                            "2 µM",
                                            "4 µM",
                                            "8 µM")), 
           y = fold_change)) +
  geom_boxplot(outlier.shape = NA) +
  geom_jitter(width = 0.2, shape = 1, alpha = 0.6) +
  theme_bw() +
  scale_y_continuous(
    labels = comma, limits = c(0,1.75), #c(lower limit, upper limit)
    breaks = seq(0,1.75,0.25)) +  #seq(lower limit, upper limit, tick frequency)
  labs(x = "Treatment",
       y = str_wrap("Fold chance in luminescense (relative to vehicle)", 
                    width = 30)) +
  theme(
    axis.text.x = element_text(face = 'bold', angle = 45, hjust = 1), 
    axis.title.x = element_text(face = 'bold'),
    axis.text.y = element_text(face = 'bold'), 
    axis.title.y = element_text(face = 'bold'),
    legend.title = element_text(face = 'bold'),
    legend.text = element_text(face = 'bold'),
    strip.text.x = element_text(face = 'bold')) +
  facet_grid(~factor(Drug, levels = c("BIM5078", "BI6015")))

ggsave("BIBIM_fix.pdf", width = 5, height = 4, path = "./output_data/")

###BI6015
BB_fix %>%
  filter(Outlier == "Normal", 
         Drug == "BI6015") %>%
  group_by(Treatment) %>%
  summarise(n = n())

BB_fix %>% 
  filter(Outlier == "Normal", Drug == "BI6015") %>%
  compare_means(formula = fold_change~Treatment,
                ref.group = "Vehicle", 
                p.adjust.method = "bonf", 
                method = "t.test")

aov_BB_fix <- aov(data = BB_fix %>% filter(Outlier == "Normal", 
                                           Drug == "BI6015"), 
                  formula = fold_change ~ Treatment)
summary(aov_BB_fix)
Anova(aov_BB_fix, type = "3")

###BIM5078
BB_fix %>%
  filter(Outlier == "Normal", 
         Drug == "BIM5078") %>%
  group_by(Treatment) %>%
  summarise(n = n())

BB_fix %>% 
  filter(Outlier == "Normal", 
         Drug == "BIM5078") %>%
  compare_means(formula = fold_change~Treatment,
                ref.group = "Vehicle", 
                p.adjust.method = "bonf", 
                method = "t.test")

aov_BB_fix <- aov(data = BB_fix %>% filter(Outlier == "Normal",
                                           Drug == "BIM5078"), 
                  formula = fold_change ~ Treatment)
summary(aov_BB_fix)
Anova(aov_BB_fix, type = "3")
```

```{r}
#| label: BIM5078 and BI6015 effect on ApoB levels (homogenization endpoint)
BB_hom <- read_plate("Figure_5_data/BIBIM_hom_summary_data.csv", well_ids_column = "Wells")
BB_hom <- BB_hom %>%
  pivot_longer(cols = c(3:ncol(.)), names_to = "Experiment", values_to = "RLU") %>%
  filter(Treatment != "Empty") %>%
  separate(col = "Experiment", into = c("Experiment", "Drug"), sep = "-")

BB_hom_veh_summary <- BB_hom %>% 
  group_by(Experiment, Treatment, Drug) %>% 
  summarise(vehicle_mean = mean(RLU)) %>% 
  filter(Treatment == "Vehicle") %>%
  ungroup() %>%
  select(-Treatment)

BB_hom <- BB_hom %>% 
  full_join(BB_hom_veh_summary, join_by(Drug, Experiment)) %>%
  mutate(fold_change = RLU/vehicle_mean)

BB_hom <- BB_hom %>% 
  group_by(Treatment, Drug) %>%
  mutate(fold_change_Q1 = (quantile(fold_change)[[2]]), 
         fold_change_Q3 = (quantile(fold_change)[[4]]), 
         fold_change_IQR = IQR(fold_change),
         Outlier = if_else(fold_change > fold_change_Q3+1.5*(fold_change_IQR) |
                             fold_change < fold_change_Q1-1.5*(fold_change_IQR),
                           "Outlier", 
                           "Normal"))

y_lim <- BB_hom %>% 
  filter(Outlier == "Normal") %>%
  pull(fold_change) %>% 
  max(.) %>% 
  round_any(., 1, ceiling)

ggplot(data = BB_hom %>% 
         filter(Outlier == "Normal"),
       aes(x = factor(Treatment, levels = c("Vehicle",
                                            "5 µM lomitapide",
                                            "0.0625 µM",
                                            "0.125 µM",
                                            "0.25 µM",
                                            "0.5 µM",
                                            "1 µM",
                                            "2 µM",
                                            "4 µM",
                                            "8 µM")), 
           y = fold_change)) +
  geom_boxplot(outlier.shape = NA) +
  geom_jitter(width = 0.2, shape = 1, alpha = 0.6) +
  theme_bw() +
  scale_y_continuous(
    labels = comma, limits = c(0,1.75), #c(lower limit, upper limit)
    breaks = seq(0,1.75,0.25)) +  #seq(lower limit, upper limit, tick frequency)
  labs(x = "Treatment",
       y = str_wrap("Fold chance in luminescense (relative to vehicle)", 
                    width = 30)) +
  theme(
    axis.text.x = element_text(face = 'bold', angle = 45, hjust = 1), 
    axis.title.x = element_text(face = 'bold'),
    axis.text.y = element_text(face = 'bold'), 
    axis.title.y = element_text(face = 'bold'),
    legend.title = element_text(face = 'bold'),
    legend.text = element_text(face = 'bold'),
    strip.text.x = element_text(face = 'bold')) +
  facet_grid(~factor(Drug, levels = c("BIM5078", "BI6015")))

ggsave("BIBIM_hom.pdf", width = 5, height = 4, path = "./output_data/")

###BI6015
BB_hom %>%
  filter(Outlier == "Normal", 
         Drug == "BI6015") %>%
  group_by(Treatment) %>%
  summarise(n = n())

BB_hom %>% 
  filter(Outlier == "Normal", 
         Drug == "BI6015") %>%
  compare_means(formula = fold_change~Treatment,
                ref.group = "Vehicle", 
                p.adjust.method = "bonf", 
                method = "t.test")

aov_BB_hom <- aov(data = BB_hom %>% filter(Outlier == "Normal",
                                           Drug == "BI6015"), 
                  formula = fold_change ~ Treatment)
summary(aov_BB_hom)
Anova(aov_BB_hom, type = "3")

###BIM5078
BB_hom %>%
  filter(Outlier == "Normal", 
         Drug == "BIM5078") %>%
  group_by(Treatment) %>%
  summarise(n = n())

BB_hom %>% 
  filter(Outlier == "Normal", 
         Drug == "BIM5078") %>%
  compare_means(formula = fold_change~Treatment,
                ref.group = "Vehicle", 
                p.adjust.method = "bonf", 
                method = "t.test")

aov_BB_hom <- aov(data = BB_hom %>% filter(Outlier == "Normal",
                                           Drug == "BIM5078"), 
                  formula = fold_change ~ Treatment)
summary(aov_BB_hom)
Anova(aov_BB_hom, type = "3")
```

```{r}
#| label: BIM5078 and BI6015 effect on NanoLuciferase activity
BB_NL <- read_plate("Figure_5_data/BIBIM_NL_activity_summary_data.csv", well_ids_column = "Wells")
BB_NL <- BB_NL %>%
  pivot_longer(cols = c(3:ncol(.)), names_to = "Experiment", values_to = "RLU") %>%
  filter(Treatment != "Empty") %>%
  separate(col = "Experiment", into = c("Experiment", "Drug"), sep = "-")

BB_NL_veh_summary <- BB_NL %>% 
  group_by(Experiment, Treatment, Drug) %>% 
  summarise(vehicle_mean = mean(RLU)) %>% 
  filter(Treatment == "Vehicle") %>%
  ungroup() %>%
  select(-Treatment)

BB_NL <- BB_NL %>% 
  full_join(BB_NL_veh_summary, join_by(Drug, Experiment)) %>%
  mutate(fold_change = RLU/vehicle_mean)

BB_NL <- BB_NL %>% 
  group_by(Treatment, Drug) %>%
  mutate(fold_change_Q1 = (quantile(fold_change)[[2]]), 
         fold_change_Q3 = (quantile(fold_change)[[4]]), 
         fold_change_IQR = IQR(fold_change),
         Outlier = if_else(fold_change > fold_change_Q3+1.5*(fold_change_IQR) |
                             fold_change < fold_change_Q1-1.5*(fold_change_IQR),
                           "Outlier", 
                           "Normal"))

y_lim <- BB_NL %>% 
  filter(Outlier == "Normal") %>%
  pull(fold_change) %>% 
  max(.) %>% 
  round_any(., 1, ceiling)

ggplot(data = BB_NL %>% 
         filter(Outlier == "Normal"),
       aes(x = factor(Treatment, levels = c("Vehicle",
                                            "400 nM NL inhibitor",
                                            "0.0625 µM",
                                            "0.125 µM",
                                            "0.25 µM",
                                            "0.5 µM",
                                            "1 µM",
                                            "2 µM",
                                            "4 µM",
                                            "8 µM")), 
           y = fold_change)) +
  geom_boxplot(outlier.shape = NA) +
  geom_jitter(width = 0.2, shape = 1, alpha = 0.6) +
  theme_bw() +
  scale_y_continuous(
    labels = comma, limits = c(0,1.75), #c(lower limit, upper limit)
    breaks = seq(0,1.75,0.25)) +  #seq(lower limit, upper limit, tick frequency)
  labs(x = "Treatment",
       y = str_wrap("Fold chance in luminescense (relative to vehicle)", 
                    width = 30)) +
  theme(
    axis.text.x = element_text(face = 'bold', angle = 45, hjust = 1), 
    axis.title.x = element_text(face = 'bold'),
    axis.text.y = element_text(face = 'bold'), 
    axis.title.y = element_text(face = 'bold'),
    legend.title = element_text(face = 'bold'),
    legend.text = element_text(face = 'bold'),
    strip.text.x = element_text(face = 'bold')) +
  facet_grid(~factor(Drug, levels = c("BIM5078", "BI6015")))

ggsave("BIBIM_NL.pdf", width = 5, height = 4, path = "./output_data/")

###BI6015
BB_NL %>%
  filter(Outlier == "Normal", 
         Drug == "BI6015") %>%
  group_by(Treatment) %>%
  summarise(n = n())

BB_NL %>% 
  filter(Outlier == "Normal", 
         Drug == "BI6015") %>%
  compare_means(formula = fold_change~Treatment,
                ref.group = "Vehicle", 
                p.adjust.method = "bonf", 
                method = "t.test")

aov_BB_NL <- aov(data = BB_NL %>% filter(Outlier == "Normal",
                                         Drug == "BI6015"), 
                  formula = fold_change ~ Treatment)
summary(aov_BB_NL)
Anova(aov_BB_NL, type = "3")

###BIM5078
BB_NL %>%
  filter(Outlier == "Normal", 
         Drug == "BIM5078") %>%
  group_by(Treatment) %>%
  summarise(n = n())

BB_NL %>% 
  filter(Outlier == "Normal", 
         Drug == "BIM5078") %>%
  compare_means(formula = fold_change~Treatment,
                ref.group = "Vehicle", 
                p.adjust.method = "bonf", 
                method = "t.test")

aov_BB_NL <- aov(data = BB_NL %>% filter(Outlier == "Normal",
                                         Drug == "BIM5078"), 
                  formula = fold_change ~ Treatment)
summary(aov_BB_NL)
Anova(aov_BB_NL, type = "3")
```

```{r}
#| label: BIM5078 effect on lipoprotein class (native PAGE)
BIM5078_PAGE <- read.csv(file = "Figure_5_data/BIM5078_nPAGE_data_summary.csv")

BIM5078_PAGE <- BIM5078_PAGE %>% 
  pivot_longer(cols = c("ZM", "VLDL", "IDL", "LDL"),
               names_to = "LP_class", 
               values_to = "Bin_value")

BIM5078_PAGE <- BIM5078_PAGE %>% 
  group_by(Experiment, Treatment, Replicate) %>%
  summarize(Total = sum(Bin_value)) %>%
  full_join(BIM5078_PAGE) %>%
  mutate(Proportion = (Bin_value/Total)*100)

ggplot(data = BIM5078_PAGE,
       aes(x = factor(Treatment, c("Vehicle", "5 µM lomitapide", "8 µM BIM5078")),
           y = Proportion)) +
  geom_boxplot(outlier.shape = NA) +
  geom_jitter(width = 0.2, shape = 1, alpha = 0.6) +
  theme_bw() +
  scale_y_continuous(
    labels = comma, limits = c(0,80), #c(lower limit, upper limit)
    breaks = seq(0,80,10)) +  #seq(lower limit, upper limit, tick frequency)
  labs(x = "Treatment",
       y = "Proportion of total lipoproteins (%)") +
  theme(
    axis.text.x = element_text(face = 'bold', angle = 45, hjust = 1), 
    axis.title.x = element_text(face = 'bold'),
    axis.text.y = element_text(face = 'bold'), 
    axis.title.y = element_text(face = 'bold'),
    legend.title = element_text(face = 'bold'),
    legend.text = element_text(face = 'bold'),
    strip.text.x = element_text(face = 'bold')) +
  facet_grid(~factor(LP_class, c("ZM", "VLDL", "IDL", "LDL")))
ggsave("BIM5078_nPAGE.pdf", width = 5, height = 4, path = "./output_data/")

BIM5078_PAGE %>%
  group_by(Treatment, LP_class) %>%
  summarise(n = n())

means_comp <- aov(data = BIM5078_PAGE %>% filter(LP_class == "ZM"), formula = Proportion ~ Treatment)
summary(means_comp)
Anova(means_comp, type = "3")

BIM5078_PAGE %>% 
  filter(LP_class == "ZM") %>%
  compare_means(formula = Proportion~Treatment,
                ref.group = "Vehicle", 
                p.adjust.method = "bonf", 
                method = "t.test")

means_comp <- aov(data = BIM5078_PAGE %>% filter(LP_class == "VLDL"), formula = Proportion ~ Treatment)
summary(means_comp)
Anova(means_comp, type = "3")

BIM5078_PAGE %>% 
  filter(LP_class == "VLDL") %>%
  compare_means(formula = Proportion~Treatment,
                ref.group = "Vehicle", 
                p.adjust.method = "bonf", 
                method = "t.test")

means_comp <- aov(data = BIM5078_PAGE %>% filter(LP_class == "IDL"), formula = Proportion ~ Treatment)
summary(means_comp)
Anova(means_comp, type = "3")

BIM5078_PAGE %>% 
  filter(LP_class == "IDL") %>%
  compare_means(formula = Proportion~Treatment,
                ref.group = "Vehicle", 
                p.adjust.method = "bonf", 
                method = "t.test")

means_comp <- aov(data = BIM5078_PAGE %>% filter(LP_class == "LDL"), formula = Proportion ~ Treatment)
summary(means_comp)
Anova(means_comp, type = "3")

BIM5078_PAGE %>% 
  filter(LP_class == "LDL") %>%
  compare_means(formula = Proportion~Treatment,
                ref.group = "Vehicle", 
                p.adjust.method = "bonf", 
                method = "t.test")
```

```{r}
#| label: BI6015 effect on lipoprotein class (native PAGE)
BI6015_PAGE <- read.csv(file = "Figure_5_data/BI6015_nPAGE_data_summary.csv")

BI6015_PAGE <- BI6015_PAGE %>% 
  pivot_longer(cols = c("ZM", "VLDL", "IDL", "LDL"),
               names_to = "LP_class", 
               values_to = "Bin_value")

BI6015_PAGE <- BI6015_PAGE %>% 
  group_by(Experiment, Treatment, Replicate) %>%
  summarize(Total = sum(Bin_value)) %>%
  full_join(BI6015_PAGE) %>%
  mutate(Proportion = (Bin_value/Total)*100)

ggplot(data = BI6015_PAGE,
       aes(x = factor(Treatment, c("Vehicle", "5 µM lomitapide", "8 µM BI6015")),
           y = Proportion)) +
  geom_boxplot(outlier.shape = NA) +
  geom_jitter(width = 0.2, shape = 1, alpha = 0.6) +
  theme_bw() +
  scale_y_continuous(
    labels = comma, limits = c(0,80), #c(lower limit, upper limit)
    breaks = seq(0,80,10)) +  #seq(lower limit, upper limit, tick frequency)
  labs(x = "Treatment",
       y = "Proportion of total lipoproteins (%)") +
  theme(
    axis.text.x = element_text(face = 'bold', angle = 45, hjust = 1), 
    axis.title.x = element_text(face = 'bold'),
    axis.text.y = element_text(face = 'bold'), 
    axis.title.y = element_text(face = 'bold'),
    legend.title = element_text(face = 'bold'),
    legend.text = element_text(face = 'bold'),
    strip.text.x = element_text(face = 'bold')) +
  facet_grid(~factor(LP_class, c("ZM", "VLDL", "IDL", "LDL")))
ggsave("BIM6015_nPAGE.pdf", width = 5, height = 4, path = "./output_data/")

BI6015_PAGE %>%
  group_by(Treatment, LP_class) %>%
  summarise(n = n())

means_comp <- aov(data = BI6015_PAGE %>% filter(LP_class == "ZM"), formula = Proportion ~ Treatment)
summary(means_comp)
Anova(means_comp, type = "3")

BI6015_PAGE %>% 
  filter(LP_class == "ZM") %>%
  compare_means(formula = Proportion~Treatment,
                ref.group = "Vehicle", 
                p.adjust.method = "bonf", 
                method = "t.test")

means_comp <- aov(data = BI6015_PAGE %>% filter(LP_class == "VLDL"), formula = Proportion ~ Treatment)
summary(means_comp)
Anova(means_comp, type = "3")

BI6015_PAGE %>% 
  filter(LP_class == "VLDL") %>%
  compare_means(formula = Proportion~Treatment,
                ref.group = "Vehicle", 
                p.adjust.method = "bonf", 
                method = "t.test")

means_comp <- aov(data = BI6015_PAGE %>% filter(LP_class == "IDL"), formula = Proportion ~ Treatment)
summary(means_comp)
Anova(means_comp, type = "3")

BI6015_PAGE %>% 
  filter(LP_class == "IDL") %>%
  compare_means(formula = Proportion~Treatment,
                ref.group = "Vehicle", 
                p.adjust.method = "bonf", 
                method = "t.test")

means_comp <- aov(data = BI6015_PAGE %>% filter(LP_class == "LDL"), formula = Proportion ~ Treatment)
summary(means_comp)
Anova(means_comp, type = "3")

BI6015_PAGE %>% 
  filter(LP_class == "LDL") %>%
  compare_means(formula = Proportion~Treatment,
                ref.group = "Vehicle", 
                p.adjust.method = "bonf", 
                method = "t.test")
```

# Figure 6 - HNF4a rdu14 allele

```{r}
#| label: Characterization of lipoprotein levels throughout development of wild-type, heterozygous, and homozygous animals 
rdu14_summary <- read.csv("Figure_6_data/rdu14_characterization_summary.csv")

rdu14_summary <- rdu14_summary %>% 
  mutate("Fus(ApoBb.1-NLuc)" = ifelse(Reader == "Tecan Spark" & RLU > 1000000 | 
                                        Reader == "BioTek Synergy H1" & RLU > 2000, 
                                      "heterozygous", 
                                      "wild-type")) %>% 
  filter(HNF4 != "ND")

rdu14_3DPF_avg <- rdu14_summary %>%
  filter(Age == 3, HNF4 == "wild-type", `Fus(ApoBb.1-NLuc)` == "heterozygous") %>%
  group_by(Experiment, Clutch) %>%
  summarize(avg3 = mean(RLU))

rdu14_summary <- rdu14_summary %>%
  full_join(rdu14_3DPF_avg, join_by(Experiment, Clutch)) %>%
  filter(`Fus(ApoBb.1-NLuc)` == "heterozygous") %>%
  mutate(fold_change = RLU/avg3)

rdu14_summary <- rdu14_summary %>% 
  group_by(Experiment, Age, HNF4) %>%
  mutate(fold_change_Q1 = (quantile(fold_change)[[2]]), 
         fold_change_Q3 = (quantile(fold_change)[[4]]), 
         fold_change_IQR = IQR(fold_change),
         Outlier = if_else(fold_change > fold_change_Q3+1.5*(fold_change_IQR) |
                             fold_change < fold_change_Q1-1.5*(fold_change_IQR),
                           "Outlier", 
                           "Normal"))

y_lim <- rdu14_summary %>% 
  filter(Outlier == "Normal") %>%
  pull(fold_change) %>% 
  max(.) %>% 
  round_any(., 1, ceiling)

ggplot(data = rdu14_summary %>%
         filter(Outlier == "Normal"), 
       mapping = aes(x = factor(HNF4, levels = c("wild-type",
                                                 "heterozygous",
                                                 "homozygous")), 
                     y = fold_change)) +
  geom_boxplot(outlier.shape = NA) +
  geom_jitter(width = 0.2, shape = 1, alpha = 0.3) +
  theme_bw() +
  scale_y_continuous(
    name = "Relative Luminescence Units (RLU)",
    labels = comma, limits = c(0,1.5), #c(lower limit, upper limit)
    breaks = seq(0,1.5,0.25)) +  #seq(lower limit, upper limit, tick frequency)
  scale_x_discrete(name = "HNF4a genotype") +
  labs(color = "Treatment") +
  theme(
    axis.text.x = element_text(face = 'bold', angle = 45, hjust = 1), 
    axis.title.x = element_text(face = 'bold'),
    axis.text.y = element_text(face = 'bold'), 
    axis.title.y = element_text(face = 'bold'),
    legend.title = element_text(face = 'bold'),
    legend.text = element_text(face = 'bold'),
    strip.text.x = element_text(face = 'bold')) +
  facet_grid(cols = vars(Age))
ggsave("rdu14_LP_level.pdf", width = 5, height = 4, path = "./output_data/")

rdu14_summary %>%
  filter(Outlier == "Normal") %>%
  group_by(Age, HNF4) %>%
  summarise(n = n())

aov_rdu14_summary <- aov(data = rdu14_summary %>% 
                           filter(Outlier == "Normal"), 
                  formula = fold_change ~ HNF4 + Age)
summary(aov_rdu14_summary)

rdu14_summary %>% 
  filter(Outlier == "Normal") %>%
  compare_means(formula = fold_change~HNF4,
                group.by = "Age",
                ref.group = "wild-type", 
                p.adjust.method = "bonf", 
                method = "t.test")
```

```{r}
#| label: Pharmacogenetic interaction - rdu14 and enoxlone
PG_int <- read.csv(file = "Figure_6_data/rdu14_enoxlone_interaction_summary.csv")

PG_int <- PG_int %>% 
  mutate("Fus(ApoBb.1-NLuc)" = ifelse(RLU > 2000, "heterozygous", "wild-type")) %>% 
  filter(HNF4 != "ND") 

WT_averages <- PG_int %>%
  filter(HNF4 == "wild-type", 
           `Fus(ApoBb.1-NLuc)` == "heterozygous", 
           Treatment == "Vehicle") %>% 
  group_by(Experiment) %>%
  summarise(WT_average = mean(RLU))

PG_int <- PG_int %>% 
  filter(`Fus(ApoBb.1-NLuc)` == "heterozygous") %>%
  full_join(WT_averages) %>%
  mutate(nRLU = RLU/WT_average)

PG_int <- PG_int %>% 
  group_by(Experiment, Treatment, HNF4) %>%
  mutate(nRLU_Q1 = (quantile(nRLU)[[2]]), 
         nRLU_Q3 = (quantile(nRLU)[[4]]), 
         nRLU_IQR = IQR(nRLU),
         Outlier = if_else(nRLU > nRLU_Q3+1.5*(nRLU_IQR) | nRLU < nRLU_Q1-1.5*(nRLU_IQR),
                           "Outlier", 
                           "Normal"))

y_lim <- PG_int %>% 
  filter(Outlier == "Normal") %>%
  pull(nRLU) %>% 
  max(.) %>% 
  round_any(., 1, ceiling)

ggplot(data = PG_int %>%
       filter(Outlier == "Normal"),
       mapping = aes(x = factor(Treatment, levels = c("Vehicle", "Enoxolone")), 
                     y = nRLU)) +
  geom_boxplot(outlier.shape = NA) +
  geom_jitter(width = 0.2, shape = 1, alpha = 0.6) +
  theme_bw() +
  scale_y_continuous(
    name = "Fold Change",
    labels = comma, limits = c(0,1.75), #c(lower limit, upper limit)
    breaks = seq(0,1.75,0.25)) +  #seq(lower limit, upper limit, tick frequency)
  scale_x_discrete(name = "Treatment") +
  theme(
    axis.text.x = element_text(face = 'bold', angle = 45, hjust = 1), 
    axis.title.x = element_text(face = 'bold'),
    axis.text.y = element_text(face = 'bold'), 
    axis.title.y = element_text(face = 'bold'),
    legend.title = element_text(face = 'bold'),
    legend.text = element_text(face = 'bold'),
    strip.text.x = element_text(face = 'bold.italic')) + 
  facet_grid(~factor(HNF4, levels = c("wild-type", "heterozygous", "homozygous")))
ggsave("PG_int.pdf", width = 5, height = 4, path = "./output_data/")

PG_int %>%
  filter(Outlier == "Normal") %>%
  group_by(HNF4, Treatment) %>%
  summarise(n = n())

aov_PG_int <- aov(data = PG_int %>% filter(Outlier == "Normal"), 
                  formula = nRLU ~ HNF4 * Treatment)
summary(aov_PG_int)

PG_int %>% 
  filter(Outlier == "Normal") %>%
  compare_means(formula = nRLU~Treatment,
                group.by = "HNF4",
                p.adjust.method = "bonf", 
                method = "t.test")
```
````
